# Clonal memory of colitis accumulates and promotes tumor growth

**DOI:** 10.1101/2025.02.13.638099

**Authors:** Surya Nagaraja, Lety Ojeda-Miron, Ruochi Zhang, Ena Oreskovic, Yan Hu, Daniel Zeve, Karina Sharma, Roni R. Hyman, Qiming Zhang, Andrew Castillo, David T. Breault, Ömer H. Yilmaz, Jason D. Buenrostro

## Abstract

Chronic inflammation is a well-established risk factor for cancer, but the underlying molecular mechanisms remain unclear. Using a mouse model of colitis, we demonstrate that colonic stem cells retain an epigenetic memory of inflammation following disease resolution, characterized by a cumulative gain of activator protein 1 (AP-1) transcription factor activity. Further, we develop SHARE-TRACE, a method that enables simultaneous profiling of gene expression, chromatin accessibility and clonal history in single cells, enabling high resolution tracking of epigenomic memory. This reveals that inflammatory memory is propagated cell-intrinsically and inherited through stem cell lineages, with certain clones demonstrating dramatically stronger memory than others. Finally, we show that colitis primes stem cells for amplified expression of regenerative gene programs following oncogenic mutation that accelerate tumor growth. This includes a subpopulation of tumors that have exceptionally high AP-1 activity and the additional upregulation of pro-oncogenic programs. Together, our findings provide a mechanistic link between chronic inflammation and malignancy, revealing how long-lived epigenetic alterations in regenerative tissues may contribute to disease susceptibility and suggesting potential therapeutic strategies to mitigate cancer risk in patients with chronic inflammatory conditions.

## Introduction

Inflammation is a major risk factor for cancer, whether it is due to autoimmune disease, long-term infections or environmental exposures^1,2^, and risk often increases with the duration and severity of disease^3,4^. While diverse mechanisms may explain this connection, including the acquisition of DNA mutations^1,2^, we hypothesize that inflammation may cause lasting epigenetic alterations that lower the threshold for oncogenesis. In support, the role of the epigenome as a causal driver in cancer has become clear^5–7^. The epigenome is dynamically regulated as cells respond to environmental challenges by making new regions of their DNA accessible and active, directing transcription factor (TF) proteins to these sites, and unlocking the expression of new genes and cellular functions. During regeneration and immunity, alterations to the epigenome can persist and accumulate following repeated exposure^8–10^, improving subsequent responses to secondary stimuli^7,11–17^. While this epigenetic “memory” is largely described as adaptive, evidence suggests that it may also carry maladaptive consequences and increase future risk of disease^7,16,18^. Here, we look to study how epigenetic memory accumulates within cells, is inherited across clones, and influences predisposition for cancer. These epigenetic mechanisms may prove to be of central importance to cancer biology, providing missing mechanisms connecting lifestyle and exposure to health.

The gastrointestinal tract is an intriguing system in which to study epigenetic memory given its immense exposure to the environment^19^. There exist numerous instances of inflammation in the gut promoting malignancy, including the well-established association between ulcerative colitis (UC) and colorectal cancer (CRC). Patients with UC are at two- to five-fold more likely to develop cancer, with those diagnosed during childhood or with pancolitis carrying substantially higher risk^3,4^. The presumed cell-of-origin for CRC is the colonic stem cell^20^, a long-lived progenitor residing in the crypt base that is responsible for regenerating the entire differentiated epithelium every few days^21,22^. We hypothesized that exposure history of the intestine would be encoded in this cell population, representing a clear target for the study of how epigenetic memory may influence tissue health. To test this, we use a murine model of colitis^23^, where the colonic epithelium experiences multiple rounds of inflammation, damage and repair, to study inflammatory memory within the gut and its influence on subsequent oncogenesis.

In this work, we find that following resolution of colitis, colonic epithelia encode a memory of inflammatory damage and repair their epigenome, regulating processes such as proliferation, cell adhesion and cytoskeletal remodeling. By combining single cell multi-omics with clonal lineage tracing in *ex vivo* organoid cultures, we find that this memory is heritable and propagated through individual stem cell clonal lineages. We find the most prominent epigenetic change to be a cumulative gain in the accessibility of activator protein 1 (AP-1) transcription factor binding sites and identify disease-specific partner TFs. Finally, we demonstrate that memory of colitis and AP-1 gain is maintained through neoplastic transformation and carries the maladaptive consequence of priming colonic epithelium for increased tumor growth.

### Persistent alterations to intestinal stem cells following recovery from chronic colitis

We hypothesized that the colonic epithelium would demonstrate persistent changes in the transcriptome and epigenome following the resolution of inflammation and healing of damage. To test this, we used a murine model of chronic colitis where the colon is repeatedly injured through low-dose dextran sodium sulfate (DSS) administration^23^ (Fig. 1a). We defined three states of disease progression that reflect histologic features seen in human colitis tissue - acute injury (1 cycle of DSS), chronic injury (3 cycles of DSS) and recovery (Fig. 1b). We found that the vast majority of animals recovered or exceeded their starting body weight and epithelial crypt structures were reformed within 21 days (Extended Data Fig, 1a,b). Further, the mucosa and submucosa demonstrated an absence of immune infiltrate and CD45+ immune cells returned to levels matching that of healthy colonic tissue (Extended Data Fig. 1c-f), altogether indicating histological and morphological recovery at the cellular and organismal levels.

**Figure 1.**
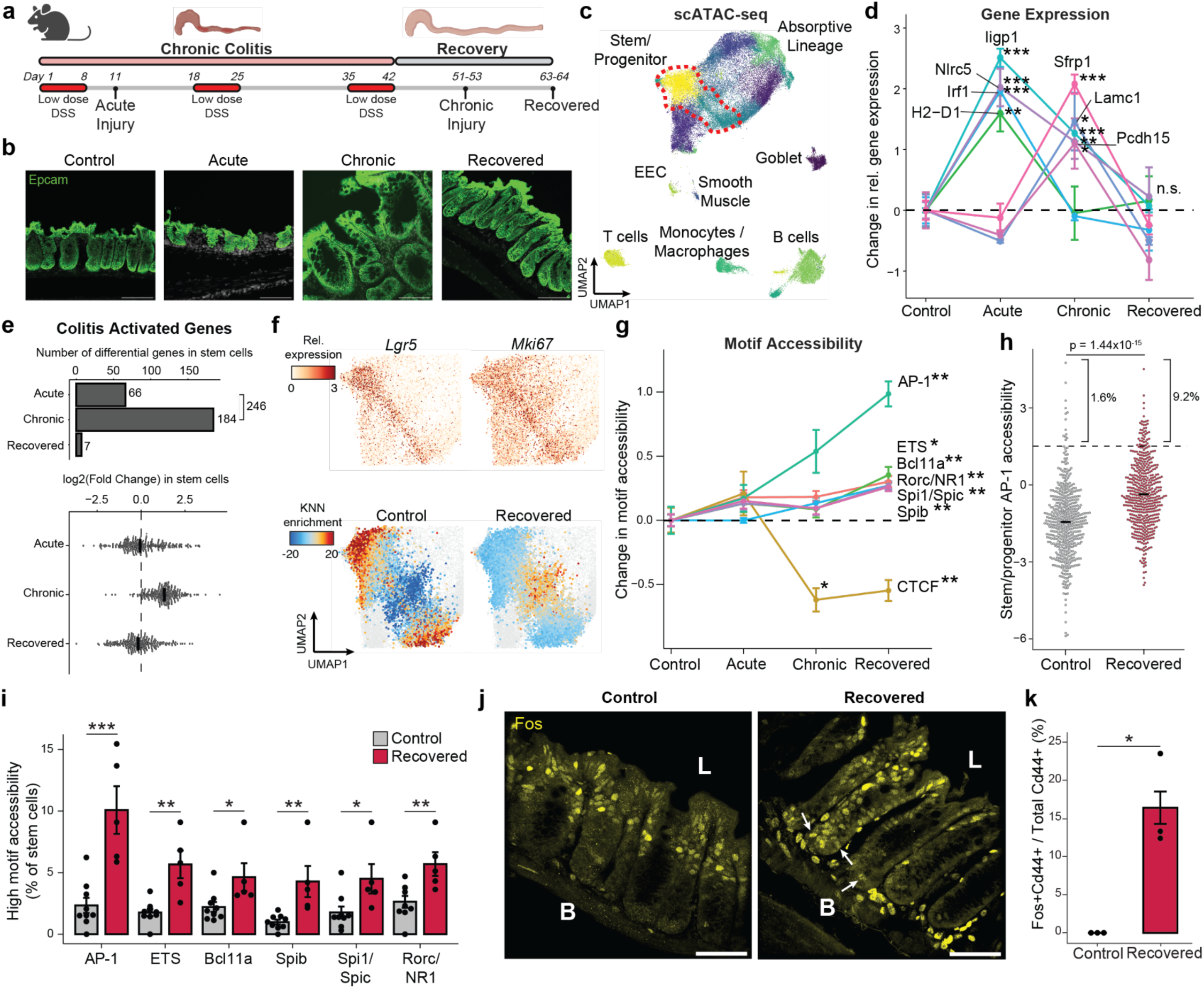
Single cell chromatin and transcriptome profiling reveals inflammatory memory in colonic stem cells. a, Paradigm for studying colitis memory. b, Immunofluorescence for Epcam at each stage of colitis. Scale bar 100 μm. c, UMAP embedding of scATAC-seq data colored by cell cluster. Stem/progenitor cells are outlined in red. d, Gene expression changes across colitis disease progression. Y-axis represents change in relative expression for each gene as compared to control. Each point represents average expression in stem cells across all animals. e, Change in stem cells for all upregulated genes during acute or chronic injury. Top, number of significant genes relative to controls. Bottom, fold change relative to controls where each point represents a gene. f, Top, expression of stem markers. Bottom, colitis stage enrichment over expected in single cell k-NN networks. g, Change in motif accessibility across colitis progression. h, AP-1 motif accessibility in single stem cells. i, Quantification of stem cells with high accessibility for each family. Each point represents the fraction of cells per animal. j, Immunofluorescence for Fos protein. Basal (B) and luminal (L) sides of crypts are marked. Arrows indicate crypt basal cells with high Fos levels. Scale bar 50 μm. k, Quantification of Cd44 and Fos co-localization per animal by immunofluorescence. All errors bars are s.e.m. (*) p < 0.05, (**) p < 0.01, (***) p < 0.001

We jointly profiled chromatin accessibility and gene expression (SHARE-seq^24^) to identify memory signatures of chronic inflammation, covering the time-course of acute, chronic and recovered tissue states (Extended Data Fig. 2a-d). Measuring 52,540 single cells across 23 animals, we identified known populations of the colonic epithelium, including *Lgr5*+ intestinal stem and progenitor cells (hereafter referred to as stem cells), absorptive enterocytes, secretory epithelial cells, and multiple immune cell populations (Fig. 1c and Extended Data Fig. 2e-g). Consistent with the recovery of the tissue, we observed no significant difference in the proportions of stem cells or cells in the absorptive lineage between recovered and control animals (Extended Data Fig. 2h).

First characterizing the stem cell response to acute injury, we found that genes related to interferon signaling and immunomodulation (e.g., *Irf1*, *Iigp1*, *H2-D1*, *Nlrc5*) were upregulated (Fig. 1d and Extended Data Fig. 3a), consistent with prior studies of epithelial inflammation^25,26^. Chronic injury further activated genes related to wound healing, involving processes such as cell junction reformation and extracellular matrix reconstruction (e.g., *Pcdh15*, *Lamc1*). Examining transcriptional memory more broadly, we found that the vast majority (>97%) of the 246 genes transcriptionally activated in stem cells during either stage of disease returned to baseline during recovery (Fig. 1e and Extended Data Fig. 3b).

Given the transcriptome demonstrated minimal changes following recovery and DSS is a poor carcinogen^27^, we hypothesized that molecular memory would be more apparent within the epigenomes of stem cells. To quantify whether cells were distinct in their overall epigenomic states, we constructed a low dimensional embedding (cisTopics^28^), found the k-nearest neighbors (k=100) of each stem cell and, for each neighborhood, quantified the proportion belonging to each stage of colitis (Extended Data Fig. 3c). Intriguingly, cells from recovered tissue were epigenomically distinct from cells derived from control tissues (Fig. 1f and Extended Data Fig. 3d,e). To more precisely define this memory, we grouped TFs by binding motif sequence similarity (n=299, Supplemental Table 1) and quantified changes in accessibility associated with these motifs^29^, revealing persistent changes following recovery (FDR<0.05, Fig. 1g). The most prominent of these was activator protein 1 (AP-1), whose motif sites cumulatively gained accessibility through colitis progression and recovery (FDR=1.27x10^-3^). We also observed memories of motifs in the ETS family (ETS, FDR=0.012; Spib, FDR=7.85x10^-3^; Spi1, FDR=5.85x10^-3^). AP-1^30–32^ and ETS^33,34^ have known oncogenic roles and are upregulated in response to ulcerative colitis. In contrast, CTCF sites showed a significant loss in accessibility during chronic colitis and recovery (FDR=8.79x10^-3^). Altogether this suggests that cells cumulatively restructure their epigenomes in response to injury and maintain these changes after recovery.

While AP-1 factors, such as Fos and Jun, are known to be activated by a diverse set of stimuli, including damage, growth and stress^35,36^, these TFs have previously been shown to be mediators of epigenetic memory^13–15^. However, these characterizations have largely been described across bulk populations of cells^14,16^. Interestingly, we quantified motif accessibility in single stem cells and identified substantial heterogeneity in TF memory representing an exceptionally high level of AP-1 motif accessibility following recovery from colitis (9.2% vs 1.6%, P=1.44x10^-15^, Fig. 1h,i and Extended Data Fig. 3f). To validate the presence of this subpopulation, we quantified Fos protein levels by immunofluorescence. Although luminal epithelial cells showed high Fos expression in healthy control tissue, we found stem and progenitor cells at the crypt base (Cd44+) with elevated Fos protein only following recovery from colitis (16.4% vs 0%, P=0.047, Fig. 1j,k). These findings highlight that while the average stem cell shows moderate epigenomic memory of colitis, approximately 10% of cells carry a prominent AP-1 memory of injury following colitis recovery.

### Epigenetic states are cell-intrinsic and clonally inherited

Immune and other non-epithelial cells are known to mediate the progression and resolution of colitis through local and systemic production of signaling factors^37,38^. We next sought to test whether colitis memory observed in colonic stem cells was cell-intrinsic or maintained by cues from surrounding cells. We reasoned that only cell-intrinsic changes would persist during the *ex vivo* culture of stem cells and, therefore, we derived organoids from colitis tissue during chronic injury. Remarkably, we found that colitis organoids progressively obtained a regenerative and hyperplastic morphology^39–41^ over 34 days of culture (Fig. 2a). Consistently, colitis-derived organoids were more proliferative (Fig. 2b), potentially representing an adaptation to repeated cycles of regenerating wounded tissue. Together, this demonstrates that cellular memory of colitis was maintained within colonic stem cells following removal from the tissue microenvironment.

**Figure 2.**
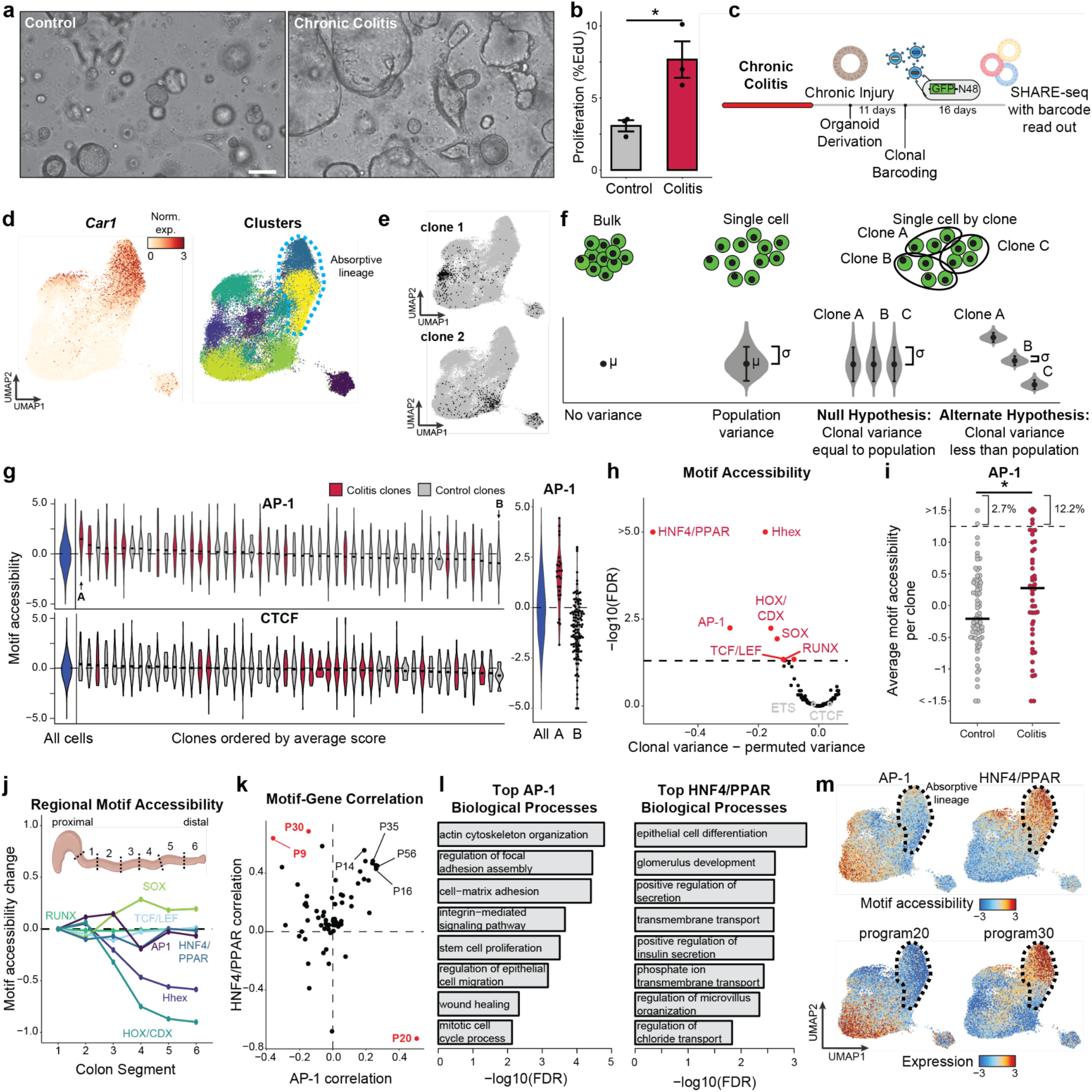
Cell-intrinsic maintenance of epigenetic states through clonal lineages. a, Organoid morphology at 34 days in culture. Scale bar 150 μm. b, Proliferation in colitis-derived and control organoids at 9 days of culture. Each point represents an organoid line. c, Schematic for lineage tracing in mouse organoids. d, UMAP embedding of scATAC-seq data in mouse organoids displaying expression of enterocyte differentiation marker *Car1* (left) and cell cluster (right). e, Examples of select clones. f, Schematic for distinguishing features that display clonal memory. g, Left, distribution of AP-1 and CTCF motif accessibility in representative organoid clones where points represent median values per clone. Right, AP-1 motif accessibility for top and bottom clones, where each point is a clone. h, Clonal memory of motif accessibility. i, Average AP-1 motif accessibility per clone. j, Motif accessibility relative to position in the colon. k, Gene program relation to motif accessibility. l, Gene ontology biological programs for AP-1 related genes (program 20, left) and HNF4/PPAR related genes (programs 9 and 30, right). m, UMAP embeddings showing motif accessibility (top) and gene program expression (bottom).

We next sought to test whether cellular memory is clonally heritable and mediated by the epigenome. Recent advances in single-cell lineage tracing provide new opportunities to measure fate transitions^42–44^. These tools exogenously introduce unique barcodes that are delivered to individual cells, propagated through cell division and transcriptionally expressed for detection through single-cell RNA-seq^42^. Inspired by this, we created SHARE-TRACE (SHARE-seq with Clonal Tracing) to simultaneously measure a cell’s clonal lineage history, gene expression, differentiation state, and chromatin accessibility. To do this, we modified the cell DNA barcode technology^42^ to improve nuclear retention and improve its compatibility with SHARE-seq (Extended Data Fig. 4a). This approach takes advantage of the scalability of SHARE-seq, which enabled us to profile 52,564 cells from six control and chronic colitis organoid lines and map the transcriptomic and epigenomic states of 172 clones (Fig. 2c-e).

The identification of hundreds of clones provided the opportunity to more sensitively quantify clonal heritability of epigenomic states. We reasoned that for any clonally remembered state, cells within a clone would more closely resemble one another than a random selection of cells. We therefore developed a permutation-based statistical approach to test heritability across TF motifs (Fig. 2f,g and Extended Data Fig. 4b,c). While *in vivo* SHARE-seq showed memory associated with ETS and CTCF motifs, we did not see clonal heritability for these motifs, suggesting that memory of these factors may be propagated by cell extrinsic mechanisms (Fig. 2h). However, we did find clonal memory of AP-1 with this approach that, as we saw with *in vivo* stem cell memory, includes both a mean shift in AP-1 activity (P=0.019) and a subpopulation of colitis-derived clones that demonstrate exceptionally high AP-1 motif accessibility (12.2% vs 2.7%; Fig. 2i). The resemblance of this distribution to that seen *in vivo* (Fig. 1h) suggests that heterogeneity in memory is maintained through clonal lineages and that stem cells particularly impacted by inflammation propagate this insult to their progeny.

Interestingly, this more-sensitive approach also uncovered clonal memory of HOX, HHEX and SOX transcription factors. We hypothesized that the stem cells within the colon may remember their developmental history and position within the colon. To test this, we divided the mouse colon in six proximal to distal segments and performed bulk ATAC-seq. Indeed, the accessibility of HOX, HHEX and SOX TFs were variable along the length of the colon (Fig. 2j and Extended Data Fig. 4d) and therefore their clonal heritability was likely attributable to the position within the colon from which the organoid was derived. Of the remaining clonal motifs, the strongest memory reflected AP-1 and the nuclear receptors Hepatocyte Nuclear Factor 4 (HNF4) and Peroxisome Proliferator-activated Receptor (PPAR), which have similar binding motif sequences. Importantly, both of these TF families were induced in colitis-derived organoids over healthy controls (Extended Data Fig. 4e-g). Overall, these analyses reveal that colitis memory is heritable and mediated by AP-1.

We next wanted to understand how the memory of AP-1 and HNF4/PPAR relate to biological functions. To this end, we derived transcriptional programs composed of co-regulated genes and, measuring clonality analogously to TFs, we observed that a subset of gene expression programs display heritability across clones (Extended Data Fig. 4h,i). To identify the target processes of clonal TFs AP-1 and HNF4/PPAR, we correlated motif accessibility with gene programs across clones and found program 20 (P20) was strongly correlated to AP-1 (r=0.50, Fig. 2k). P20 represented genes related to epithelial repair (Supplemental Table 2), including cytoskeletal remodeling, cell junction reformation and proliferation (Fig. 2l and Extended Data Fig. 4j). By contrast, HNF4/PPAR strongly correlated with P9 (r=0.64) and P30 (r=0.69), representing processes involved in colonic enterocyte maturation. Further, we found P30 genes were expressed in differentiated cells (Fig. 2m), suggesting clonal memory of HNF4/PPAR sites may represent differences in differentiation capability and stemness retention^45^.

Overall, we demonstrate that cellular memory can be maintained within clonal lineages and result in cell populations with exceptionally altered epigenetic states. Considering our finding that colitis memory promotes proliferation, the heritability of these states raises the intriguing prospect that clonal memory of chronic inflammation may give individual stem cells and their progeny a fitness advantage. Such a process may create fields within tissues with altered epigenetic states that affect future responses to stimuli and the development of disease.

### AP-1 cooperates with TFs to maintain altered cell states in colitis

Given the central importance of AP-1 accessibility in the cellular memory of colitis, we turned to understand its mechanism of action *in vivo*. We first examined whether AP-1 associated genes (P20) were expressed following tissue recovery from colitis. Unlike the accessibility of AP-1 motifs, which increased through recovery, AP-1/P20 gene expression peaked during chronic injury and declined during recovery (Fig. 3a). Consistent with this observation, we found that chromatin remained open at many individual genes despite transcription returning to baseline levels (Fig. 3b). As AP-1 is expressed broadly across tissues^46^, we sought to understand how AP-1 may be directed to have stem cell specific functions. Prior studies have reported that AP-1 interacts with tissue-specific factors to establish memory at specific genomic regions^13–15,47^. To uncover binding partners of AP-1 in colonic stem cells *in vivo*, we used seq2PRINT^48^, an approach that combines TF footprinting with deep learning to *de novo* discover sequence motifs of DNA binding proteins and localize these binding events to regulatory elements (Fig. 3c). Training seq2PRINT on chromatin accessibility footprints of *in vivo* colitis progression yielded a total of 1,838 motifs, representing 890 known motifs and 948 unknown motifs (Supplementary Data 1). This TF motif repertoire uncovered DNA sequences related to dimerization, co-binding and stability not annotated in existing motif databases (Extended Data Fig. 5a).

**Figure 3.**
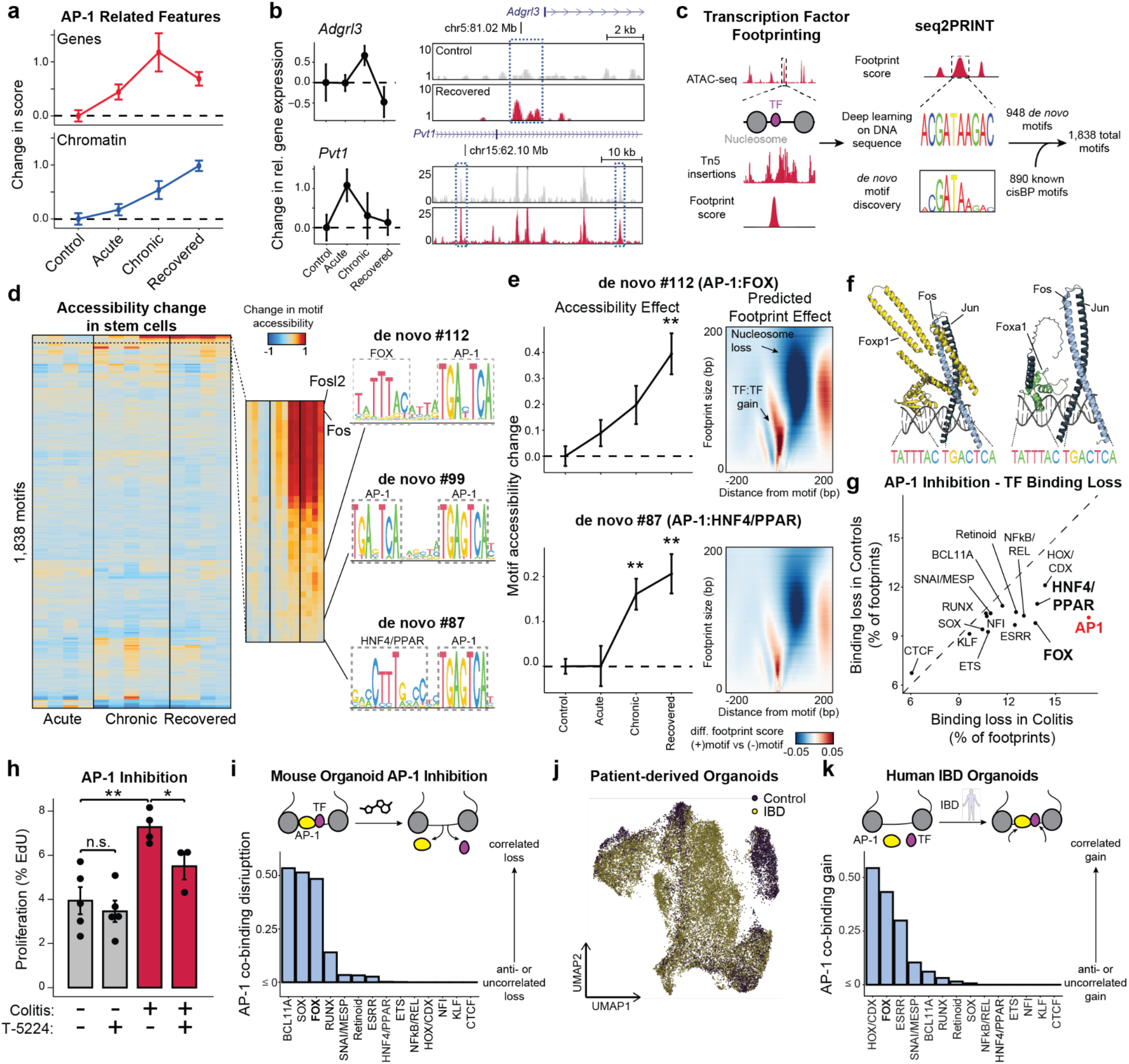
AP-1 maintains memory TF networks and altered cell states. a, Comparison of AP-1 associated gene expression scores (top) and AP-1 motif accessibility (bottom) through disease progression. b, Gene expression (left) and chromatin accessibility loci (right) at AP-1 associated genes *Adgrl3* and *Pvt1*. c, Schematic for derivation of *de novo* motifs from footprinting data. d, Accessibility of all *de novo* and known motifs. Left, each column represents the average motif score change from control for stem cells from a single animal. Right, sequence information content of *de novo* derived AP-1 composite motifs. e, Chromatin accessibility at AP-1 composite motif sites through colitis progression (left) and predicted effect of motif presence on footprint (right). For footprint effects, the x-axis represents distance from motif, the y-axis represents size of footprint and color indicates predicted change of footprint score when motif is added. f, AlphaFold3 predicted structures for Fos-Jun dimer, composite motif and either Foxa1 or Foxp1. g, Comparison of change footprint score between colitis and control organoids following 24 hours of AP-1 inhibition (T-5224, 10 μM). h, Organoid proliferation following 24 hours of AP-1 inhibition or matched vehicle control. Each point represents an organoid line. i, Disruption of co-binding with AP-1 for select TF families following 24 hours of AP-1 inhibition. Co-binding score quantifies the enrichment of binding change over chance. j, UMAP embedding of scRNA-seq data from patient-derived IBD organoids or healthy controls. k, Co-binding of select TF families at IBD-specific AP-1 footprints in human organoids. All error bars represent s.e.m. (*) p < 0.05, (**) p < 0.01, (***) p < 0.001

While this unsupervised approach once again uncovered AP-1 as the most prominent regulator of memory *in vivo*, it also identified composite motifs of AP-1 and additional TF families (Fig. 3d and Extended Data Fig. 5b). Most notable among these were associations with the forkhead box (FOX) and HNF4/PPAR motifs. Both of these composite motifs were associated with memory of colitis and predicted to increase accessibility by displacing nucleosomes (Fig. 3e). To further substantiate these associations, we identified FOX TFs expressed in colonic stem cells and modeled binding with the AP-1 heterodimer of Fos and Jun on DNA using AlphaFold3^49^ (Fig. 3f and Extended Data Fig. 5c,d). This successfully predicted that multiple FOX TFs were capable of binding the composite motif sequence adjacently to AP-1 and directly interacting with the heterodimer. Intriguingly, the manner of these predicted interactions were different across members of the FOX TF family^50^ (Extended Data Fig. 5e). For instance, Jun interacted with a portion of the DNA binding domain of Foxa1 whereas both AP-1 proteins interacted with Foxp1 at multiple domains, including the leucine zipper typically associated with homodimerization^51^. Altogether, as FOX family members have been suggested to cooperatively bind with AP-1 factors^52,53^ and play roles in intestinal wound healing as well as CRC oncogenesis^54,55^, our findings uncover a disease-specific role for FOX TFs in colonic inflammation.

To functionally interrogate the relationship between AP-1 and tissue-specific TFs, we treated organoids with T-5224^56^, a chemical inhibitor that interferes with AP-1 DNA binding, and used seq2PRINT to quantify changes in TF footprinting. Notably, inhibition of AP-1 led to the increased loss of AP-1 footprints in colitis-derived organoids as compared to controls (Fig. 3g). We also found that AP-1 inhibition preferentially disrupted binding of FOX and HNF4/PPAR, both of which were predicted to co-bind with AP-1. Functionally, AP-1 inhibition selectively blocked proliferation in colitis-derived organoids (Fig. 3h), reinforcing the causal role of AP-1 in the hyperproliferative phenotype described above.

We reasoned that the disruption in TF binding following AP-1 inhibition could be through primary interactions with AP-1 or a consequence of secondary effects associated with downstream changes to gene expression. To resolve this, we developed a computational approach to measure direct or indirect binding. We began by validating binding predictions made by seq2PRINT by performing CUT&Tag for Fos and found that differences in AP-1 footprint scores determined by seq2PRINT were more than 70% accurate in predicting differential Fos binding (Extended Data Fig. 5f-j). Encouraged by this, we calculated a co-binding score that quantified how frequently a given TF family bound at the same regulatory element as AP-1. Applying this to footprint changes following AP-1 inhibition, we found that HNF4/PPAR loss was most often distal to AP-1 binding sites (Fig. 3i), demonstrating that HNF/PPAR changes are largely secondary effects and the predicted association with AP-1 represent a minority of binding events. By contrast, we found that loss in FOX footprints strongly correlated with AP-1 loss in the same regulatory element, further supporting the prominence of this relationship and indicating that FOX binding at these disease-specific sites required the presence of AP-1.

We next extended this analysis to the gains in AP-1 binding predicted in inflammatory memory. Consistent with our observations in organoids, we found that AP-1 footprints formed in primary tissue after recovery from colitis correlated most strongly with adjacent (< 20 bp) gains in FOX binding (Extended Data Fig. 5k-m). To determine the relevancy of this association to human disease, we performed SHARE-seq in organoids derived from patients with inflammatory bowel disease (IBD) or healthy controls (Fig. 3j and Extended Data Fig. 6). Prominently, our analysis predicted that novel AP-1 binding sites formed in IBD were strongly associated with proximal FOX binding as well (Fig. 3k), revealing a role for FOX TFs in memory of intestinal inflammation in multiple model systems and across species. Taken together, these findings demonstrate AP-1 drives restructuring of gene regulatory networks with tissue-specific factors in inflammatory memory and facilitates altered cellular phenotypes.

### Memory of chronic colitis is preserved through transformation and promotes adenoma growth

We next hypothesized that chronic colitis, mediated by clonal heritability of AP-1 accessibility, may prime the colonic epithelium for malignant transformation. To test this hypothesis, we induced adenomas through *APC* loss (*Cdx2*:*CreER*^T2^;*APC^fl/fl^*)^57^ in mice recovered from colitis and naive controls (Fig. 4a). Strikingly, we found that colitis-associated adenomas were larger in size as compared to controls (P=0.042), with a greater fraction of tumors greater than 1 mm in diameter (P=0.002, Fig. 4b,c and Extended Data Fig. 7a).

**Figure 4.**
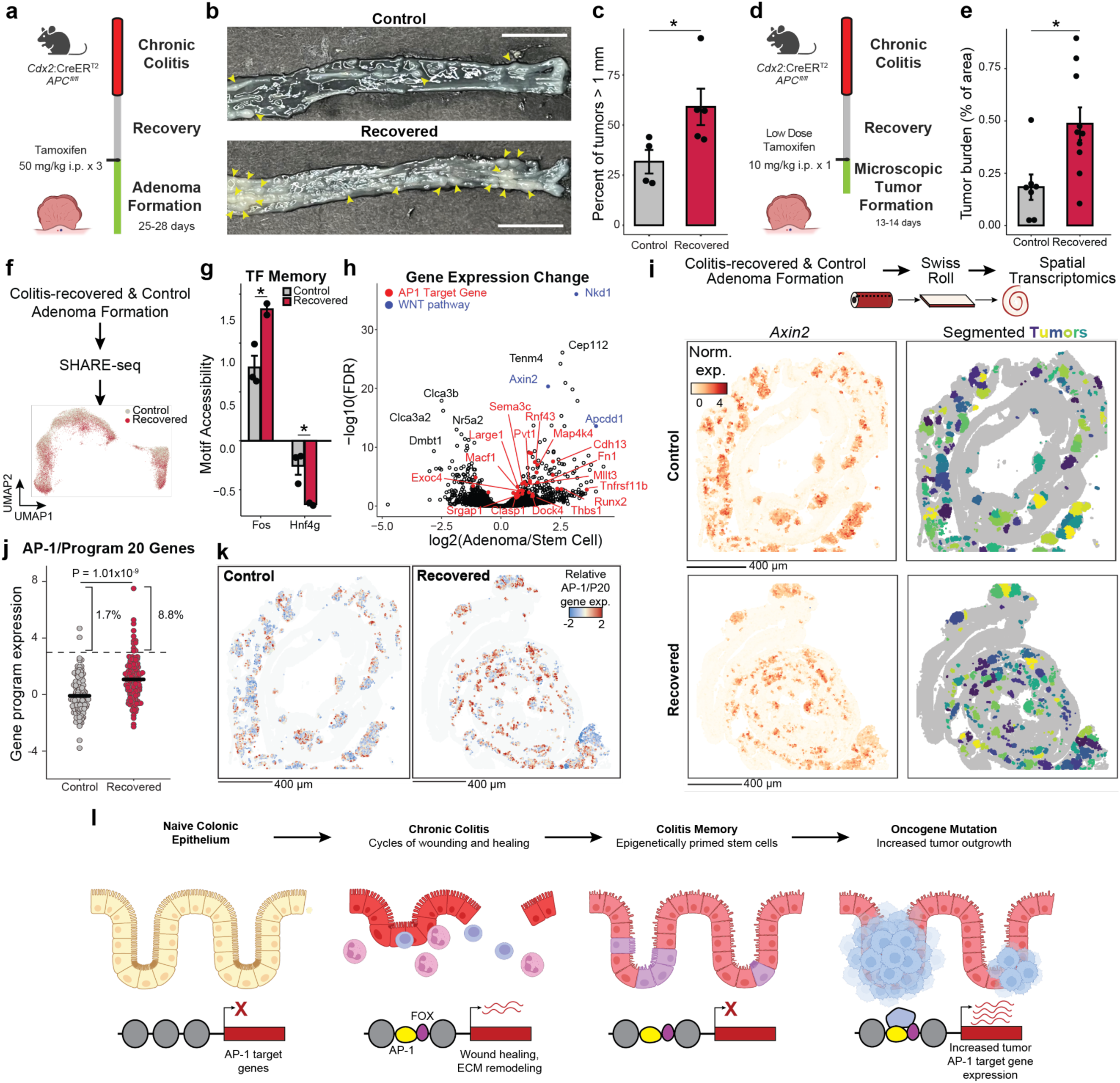
Epigenetic memory of colitis promotes tumor growth. a, Schematic for adenoma induction following colitis recovery. b, Gross images of tumor formation in recovered and control animals. Scale bar 1 cm. c, Quantification of large adenomas. Each point represents an animal. d, Schematic for measuring tumor initiation following low dose tamoxifen administration. e, Quantification of microscopic tumor area following low dose initiation. Each point represents an animal. f, Schematic for performing SHARE-seq. g, Transcription factor memory in adenoma cells following transformation. h, Gene expression changed following transformation. i, Left, spatial *Axin2* expression. Right, microscopic tumor identification with each tumor colored by a distinct color. Scale bar 400 μm. j, Expression scores of AP-1/P20. Each point represents expression in an individual tumor. k, Spatial expression of AP-1/P20 expression score in identified tumor cells. Scale bar 400 μm. l, Model of colitis memory priming epithelium for increased tumor growth. Error bars are s.e.m. (*) p < 0.05, (**) p < 0.01, (***) p < 0.001

Surprisingly, colitis recovered animals did not carry higher numbers of macroscopic tumors and adenomas did not demonstrate higher proliferation or clone-forming potential (Extended Data Fig. 7b-d). We hypothesized that epigenetic memory following colitis promoted initial tumor outgrowth and grossly larger tumors occurred due to the close proximity and clustering of many microscopic tumors. Therefore, we treated mice with a single low dose of tamoxifen to induce sparse tumor initiation and harvested tissue prior to gross adenoma formation, allowing the assessment of early microscopic growth^58^ (Fig. 4d and Extended Data Fig. 7e,f). Consistent with colitis recovered epithelium being primed for tumorigenesis, we found increased tumor burden at this early time point (P=0.012, Fig. 4e). Furthermore, we did not observe a greater number of initiation events in animals recovered from colitis but instead that each microscopic tumor was larger (P=1.79x10^−5^, Extended Data Fig. 7g,h). Similar to the heterogeneity observed in clonal memory of AP-1 accessibility (Fig. 2i), we find a greater proportion of these microscopic lesions to be excessively large in colitis recovered tissue (8.7% vs 2.5%), raising the intriguing possibility that these represent clonal fields expanded from stem cells with strong memory of inflammation. Altogether, these findings indicate that the pro-tumorigenic effect of chronic colitis was due to an acceleration of growth following initial oncogene mutation. While previous studies have found that colitis injury after inducing oncogenic mutations increases tumor formation^59,60^, we show here that the pro-oncogenic effect of colitis is maintained even after recovery and resolution.

To characterize the molecular differences associated with this phenomenon, we performed SHARE-seq on control and colitis-associated tumors, identifying both adenoma cells and adjacent non-neoplastic epithelium (n=38,382 cells; Fig. 4f and Extended Data Fig. 8a-c). Interestingly, we found that colitis-associated adenoma cells displayed higher accessibility at AP-1 motifs (P=0.025, Fig. 4g and Extended Data Fig. 8d) and decreased accessibility at HNF4/PPAR sites (P=0.048). Predicted AP-1 binding changes in adenoma cells continued to demonstrate cooperativity with FOX factors but additionally gained novel co-binding relationships with ESRR and HOX/CDX factors (Extended Data Fig. 8e,f), indicating that memory TFs may acquire novel roles and functions following oncogenic transformation. Altogether, these findings show that AP-1 memory of colitis is preserved through oncogenic transformation. Intriguingly, we found many genes associated with AP-1 to be upregulated in adenomas relative to normal stem cells (Fig. 4h), suggesting that memory of AP-1 may be oncogenic. To relate these molecular differences to individual tumors, we performed spatial RNA sequencing following adenoma formation. As *APC* loss drives unrestrained WNT pathway activation, we were able to identify 254 distinct tumors based on *Axin2* expression and image segmentation (Fig. 4i and Extended Data Fig. 9a). Across the 117 control and 137 recovered tumors, we found that colitis-associated adenomas had elevated levels of AP-1/P20 gene expression (P=1.01x10^-9^, Fig. 4j,k and Extended Data Fig. 9b). In line with the exceptionally large AP-1 memory that we observed in a subpopulation of stem cells *in vivo* (Fig. 1h) and organoid clones (Fig. 2i), we found that a fraction of colitis-associated tumors exhibited particularly strong expression of AP-1 associated genes (8.8% vs 1.7%). In addition to expected expression differences in repair-related genes associated with P20, these exceptional tumors upregulated oncogenic programs related to platelet-derived growth factor (PDGF) signaling^61^, proliferation and vasculogenesis (Extended Data Fig. 9c,d). Together, these findings demonstrate that recovery from colitis primes tumors for increased AP-1 associated gene expression and an exceptionally high AP-1 subpopulation robustly activates additional pro-oncogenic programs.

Overall, these findings suggest a model where chronic inflammation encodes an epigenetic memory of repair in colonic stem cells that promotes tumor growth through progressive gain of AP-1 and tissue-specific TF accessibility at pro-proliferative genes (Fig. 4l). Rather than by increasing tumor initiation, inflammatory memory promotes tumorigenesis by promoting malignant outgrowth once a stem cell acquires an oncogenic mutation, thereby contributing to the increased incidence of cancer associated with chronic inflammation.

## Discussion

Overall, we reveal that chronic inflammatory disease creates cellular memory through the accumulation of epigenetic changes. Following repeated cycles of inflammation, damage and healing, colonic stem cells heterogeneously encode a memory of regeneration in their epigenome, including the emergence of exceptionally high AP-1 cells, that lowers the threshold for tumor formation. Furthermore, we find that this high AP-1 state is propagated clonally and maintained following neoplasia, promoting a hyperproliferative tumor state. These findings provide an epigenetic mechanism that connects inflammatory diseases with malignancy.

Our observation of macroscopically larger tumors strongly suggests that clonal fields of epigenetically primed cells emerge in close proximity to one another, consistent with a model of “field cancerization”^62^. While this framework has typically been used to describe fields of somatic mutations, this work suggests a similar model where chronic colitis creates clonal fields of cells carrying fitness-conferring “epi-mutations”. In line with this, recent work demonstrated that colons of UC patients consist of a “patchwork” of millimeter-sized clonal fields^63^, representing massive expansions of single stem cells when compared to healthy colonic epithelium. However, these fields did not show positive selection for typical CRC driver mutations or mutations enriched in colitis-associated tumors^64–66^. Our findings indicate this may instead be explained by heritable epigenetic alterations, largely maintained by subpopulations of stem clones bearing exceptional AP-1 accessibility, following chronic inflammation and the pro-proliferative effect of this memory.

We find that the pro-oncogenic memory of AP-1 following repeated damage is mediated by an interaction with FOX transcription factors. Previous studies have identified AP-1, and additional binding partners, as regulators of memory in inflammation, repair and therapeutic resistance^13–16,67^. Integrating these studies, AP-1 appears to be a central regulator of memory across tissues and functions^68^. Motivated by this, we suspect that diverse lifestyle and environmental factors such as diet, infection and age^69^ may all encode memory through AP-1, cumulatively influencing health and disease risk.

With rates of inflammatory bowel diseases^70^ and the incidence of early-onset colorectal cancer rising globally^71,72^, our findings carry both diagnostic and therapeutic implications for patients. Memory-related epigenetic signatures linked to cancer could allow for tracking of oncogenic risk in patients prior to the formation of visible neoplastic lesions. Similarly, inhibition of AP-1 factors or other associated transcription factor families could erase pathologic cellular memory and mitigate its maladaptive consequences, offering a promising avenue for disease prevention in patients with chronic diseases.

## Acknowledgments

We are grateful to the Buenrostro Laboratory for helpful feedback throughout this body of work. We would like to thank K. McKinley for advice and support throughout this project. We also would like to thank the Harvard Stem Cell & Regenerative Biology Histology Core (C. MacGillivray, D. Faria) for tissue processing and staining and the Broad Spatial Technology Platform (M. Lipinski, T. Aleksanyan, S.L. Farhi) for performing Visium HD. Finally, we would like to thank M. Greenberg for the generous donation of the Fos antibody for CUT&Tag experiments.

## Author contributions

S.N. and J.D.B. conceived of the project and designed experiments. S.N., L.O.M., and E.O. performed *in vivo* animal work and *ex vivo* mouse organoid experiments. S.N., L.O.M., R.Z. and Y.H. performed data analysis. Q.Z. performed clonogenicity experiments with support of O.H.Y. D.Z. and K.S. performed human organoid work with support of D.T.B. R.R.H. and A.C. assisted with animal care and experimentation. J.D.B. supervised all aspects of the study. S.N. and J.D.B. wrote the manuscript with input from all authors.

## Data and materials availability

As of February 13, 2025, raw data has been provided to the Impact of Genomic Variation on Function (IGVF) consortium and may be accessed at https://tinyurl.com/nagaraja-buenrostro-2025. We are in the process of uploading processed data files to the same location and code to GitHub. This preprint will soon be updated accordingly. Access to data will be made available upon request.

## Methods

### Animals and Cell Lines

#### Animal work

C57BL/6J (Strain #:000664) and *Cdx2*:CreER^T2^;*APC^f^*^l/fl^;*Kras*^WT^ (Strain #:035169)^57^ were obtained from The Jackson Laboratory. Mice were housed in individually ventilated cages at a maximum density of five mice per cage with ad libitum access to food and water. The colony room was kept on a 12 h/12 h light–dark cycle. All animal handling and experiments were conducted in accordance with procedures approved by the Institutional Animal Care and Use Committee at Harvard University.

#### Mouse organoid derivation and culture

Colitis organoids were derived from whole colonic tissue 11 days following cessation of the third cycle of DSS. Animals were anesthetized with 2,2,2-tribromoethanol (Sigma #T48402-25G) and cardiac perfusion was performed with PBS to remove peripheral immune cells. Epithelium was removed by incubating colonic tissue in EDTA Solution (see “Colon tissue processing and cell sorting” below) supplemented with 100 ug/mL primocin (Invitrogen #ant-pm-05) for 20 to 30 minutes and scraping the luminal surface with a glass slide. Epithelial fragments were washed once with ADMEM and resuspended in Crypt Basal (ADMEM, 10 mM HEPES, 1X GlutaMax (Thermo #35050061), 1X Pen-Strep (Thermo #15140122), 1X N2 Supplement (Thermo #17502048), 1X B27 Supplement (Thermo #17504044), 1 mM N-acetylcysteine (Sigma #A9165-5G)) before mixing with an equal volume of Matrigel (Corning #47743-722). Crypts were plated as ∼30 uL domes in a six-well plate and allowed 10 to 15 minutes to polymerize. Colon organoids were grown in WENR media: 50% ENR (Crypt Basal with 50 ng/mL EGF (Thermo #PMG8041), 100 ng/mL Noggin (Peprotech #250-38), 1:100 Rspondin conditioned media) and 50% Wnt conditioned media. Conditioned media was generated in-house from L Wnt-3a cells (ATCC #CRL 2647) or HA-R-Spondin1-Fc 293T Cells (R&D #3710-001-01). Organoids were passaged every 7 to 10 days by mild dissociation in TrypLE (Thermo #12604039) for 8-10 minutes, triturating every 4 to 5 minutes and quenching with 10% FBS in ADMEM. When collected for SHARE-seq, organoids were dissociated to near single-cell for 15 minutes in TrypLE and quenched before treating with 1:100 recombinant DNase (Roche #04716728001) in ADMEM at RT for 5 minutes to reduce dead cell DNA contamination. Cells were then washed and frozen in CryoStor at -80C prior to SHARE-seq.

#### Human organoid derivation and culture

Human organoid lines were derived from de-identified biopsies from grossly unaffected tissue in patients undergoing endoscopy at Boston Children’s Hospital. Informed consent and developmentally appropriate assent were obtained at Boston Children’s Hospital from the donors’ guardian and the donor, respectively. All methods were approved and carried out in accordance with the Institutional Review Board of Boston Children’s Hospital (Protocol number IRB-P00000529).

Organoids were derived from biopsies as previously described^73^. Briefly, intestinal crypts were isolated from frozen tissue and then resuspended and plated in 40μL Matrigel domes. Once established, human rectal organoids were sustained in specialized growth media (GM) that has been previously described^73^. Media changes occurred every two days during expansion, with organoids being passaged once every 6-8 days as necessary. To induce differentiation, organoids were grown in GM for 2 days post passage to allow for stem cell expansion; after which, the organoids were transitioned to differentiation media (DM). Media was changed every two days for the length of the experiment, with organoids being collected for analysis after 10 total days.

### Experimental Procedures

#### Colitis induction

Male mice aged between 8 and 15 weeks were administered dextran sulfate sodium (VWR #IC16011080) in drinking water at 1-1.5% final concentration to induce chronic colitis. Animals were weighed every day during DSS administration and every two to three days during rest periods. On the fourth day of each DSS administration, stool was tested for occult blood (VWR #10012-002) to ensure successful induction of colitis. DSS concentrations were reduced if excess disease severity was observed through any of the following metrics: frank blood in the stool at any point, weight of loss of greater than 10% before the 9th day of any cycle, failure to recover back to 90% of starting weight before the next cycle or poor body condition.

Acute injury timepoints were collected 3 days after the end of the first DSS cycle (day 11), chronic injury 9-11 days after the third cycle (days 51-53) and recovery 21-22 days after the third cycle (days 63-64).

#### Colon tissue processing and cell sorting

For *in vivo* colitis memory SHARE-seq experiments, animals were anesthetized and perfused as previously described. Entire colons were dissected, lumens were exposed and tissue was transferred to EDTA Dissociation Solution (10% FBS, 4 mM EDTA, 10 mM HEPES in PBS). Following rotation for 20 to 30 minutes at RT, epithelium was coarsely removed by scraping the luminal surface with a glass slide and remaining muscle and submucosal were crudely chopped with scissors. Both epithelial and tissue fragments were then dissociated to single cells in Advanced DMEM/F12 (ADMEM; Fisher #12-634-028) with 10 mM HEPES, 0.4 mg/mL collagenase (Millipore #C9263-25MG), 1.25 U/mL dispase (Millipore #D4693-1G), 1 U/mL DNase (Worthington Biochemical #LS002004), and 5 μM Y-27632 (R&D #1254). Cells were washed with 0.1% BSA/PBS, stained with Calcein Red-AM (BioLegend #425205) then with antibodies for Epcam (Fisher #501129753), Cd45 (BioLegend #103116), Ly6g (BioLegend #127605), and SiglecF (BioLegend #155503). DAPI (Fisher #62248) dead cell staining was performed prior to sorting.

Stained cells were sorted on a BD FACSAria for epithelial (Epcam+Cd45-) and non-granulocyte (Epcam-Cd45+Ly6g-SiglecF-) populations into ADMEM with 0.2% BSA, 0.1 U/uL Enzymatics RNase inhibitor (Qiagen #Y9240L) and 15 μM Y-27632. Cells were pelleted, resuspended in CryoStor CS10 (StemCell Technologies #07959) and stored at -80 C.

#### Histology and colitis scoring

Animals were anesthetized and perfused as described above. After colonic tissue was dissected and the luminal surface was exposed, swiss rolls or tissue fragments were fixed in 4% PFA/PBS overnight at 4C and then placed in 70% ethanol prior to dehydration and paraffin embedding. Hameotoxylin-eosin staining was performed for general histology evaluation.

Colitis scoring was performed as described in Remke et al^74^ with researchers blinded to sample identity. Immune infiltration was scored as follows: mucosa, 0 = normal, 1 = mildly increased immune infiltrate, 2 = modest infiltration, 3 = severe infiltration. Submucosa, 0 = normal, 1 = mild to modest immune infiltration, 2 = severe infiltration. Muscularis, 0 = normal, 1 = modest to severe.

Immunohistochemistry was performed for CD45 using Abcam #ab10558 and total CD45+ cells were counted in the mucosa and submucosa in all images before normalizing to total tissue area assessed. Researchers were blinded to conditions during imaging and quantification.

#### Immunofluorescence

Tissue was extracted, fixed with 4% PFA/PBS and subsequently cryoprotected in 30% sucrose/PBS before OCT embedding. Following sectioning, tissue was washed with PBS then permeabilized and blocked for 1 hour in PBS with 3% normal donkey serum (Jackson Immuno #017-000-121) and 0.5% Triton X-100. Sections were incubated overnight at 4C in Antibody Diluent (PBS with 1% NDS, 0.3% Triton X-100) with primary antibodies for Epcam (Abcam #ab213500), Fos (Synaptic System # 226 308), and/or Cd44 (BioLegend #103001). Excess antibody was removed with three PBS washes and secondary antibodies (Jackson Immuno #712-605-15, 706-586-148, 711-545-152) were added in Antibody Diluent. Following 1 to 4 hours of incubation at RT, excess antibody was washed away with two PBS washes. Nuclei were counterstained with DAPI and slides were mounted with Prolong Gold (Thermo #P36934). Imaging was performed on a Andor CR-DFLY-201-40 confocal spinning disk coupled to a Nikon Ti-E microscope. For Fos+Cd44+ co-staining analysis, Fos positivity in crypt epithelial Cd44+ cells was measured and researchers were blinded during both imaging and quantification.

#### SHARE-seq

SHARE-seq was performed with minor modifications to the protocol described in Ma et al. 2020 (https://www.protocols.io/view/share-seq-v1-6qpvrdexpgmk/v1) . For sorted cell and organoid experiments, frozen cells in CryoStor were briefly thawed (∼2 minutes) at room temperature before diluting with ice-cold PBS supplemented with 0.04% BSA, 0.1 U/uL Enzymatics RNase Inhibitor and 0.05 U/uL SUPERase RNase inhibitor (Thermo #AM2696). Cells were pelleted and supernatant was discarded prior to lysis with HLB - H-RSB (10 mM HEPES, 10 mM NaCl, 3 mM MgCl2) with 0.1% NP-40 (Thermo #28324), 0.04% BSA, 0.1 U/uL Enzymatics RNase Inhibitor and 0.05 U/uL SUPERase RNase inhibitor. Following a 5 minutes incubation on ice, buffer was diluted with HDT-2RI (H-RSB, 0.04% BSA, 0.1% Tween-20, 0.01% digitonin (Thermo #300410), 0.1 U/uL Enzymatics RNase Inhibitor and 0.05 U/uL SUPERase RNase inhibitor) and nuclei were pelleted. Supernatant was discarded and nuclei were resuspended in HDT-2RI at a density of 1 M/mL and fixed with 0.2% formaldehyde for 5 minutes at RT. Fixation was quenched with 140 mM glycine, 50 mM Tris pH 8.0 and 0.1% BSA on ice for 5 minutes. Fixed nuclei were washed once with HDT-2RI, once without SUPERase RNase inhibitor and stored at -80C until SHARE-seq was performed.

For adenoma tissue, nuclei were isolated from OCT embedded tissue. Two to four 40 μm sections were collected from each tissue block, removing excess peripheral OCT and placing into 1.5 mL tubes on dry-ice, not allowing tissue to thaw. Tubes were allowed to briefly warm before resuspending in 200 uL H-RSB with 0.1% NP-40, 0.04% BSA, Enzymatics RNase inhibitor and SUPERase RNase inhibitor. Tissue was dissociated by triturating with a P1000 for 20 strokes then a P200 for 80 strokes. Nuclei were diluted with HDT-2RI, pelleted, resuspended in 500 uL HDT-RI and filtered through a 40 μm filter (Millipore #BAH136800040-50EA) to remove large fragments of undissociated tissue. Nuclei were then fixed as described above.

Fixed nuclei were transposed as previously described^24^ with Protease Inhibitor Cocktail (Sigma #P8340) and 0.1% NP-40. Reverse transcription was performed as described in Ma et al. 2020 except using 1X Smart-seq3 Buffer (40 mM DTT, 125 mM Tris pH 8.0, 5 mM GTP, 150 mM NaCl, 12.5 mM MgCl2) in place of Maxima RT Buffer. Washes and split-pool barcoding were performed with 0.1% Tween-20 and 0.01% digitonin instead of NP-40. Sub-library generation, reverse crosslinking, cDNA pulldown and ATAC library preparation were all performed as previously described. Template switching was performed with 1X Smart-seq3 Buffer in place of Maxima RT Buffer. cDNA amplification and tagmentation was performed as previously described.

#### Organoid proliferation and AP-1 inhibition

Organoids were passaged once prior to AP-1 inhibition to remove dead or dying cells from primary plating. Three days following passaging, media was supplemented with 10 μM T-5224 or equal volume DMSO. After 24 hours, 10 μM EdU was added for 3 hours prior to cell dissociation as described above. A portion of cells were banked for ATAC-seq and footprinting while the rest were fixed with 4% PFA/PBS for 10 min at room temperature, washed with PBS then permeabilized with 0.5% Triton X-100 in PBS. Following two washes with 3% BSA/PBS, EdU staining was performed with Click-iT EdU Assay kit (Life Tech #C10340) and cells were counterstained with DAPI. Percent EdU was measured on an Attune CytPix cytometer as EdU+ cells over total DAPI+ cells. For baseline EdU differences between colitis and control organoids, EdU assays were performed at 9 days of culture.

#### Barcode vector cloning and library construction

The pLARRY empty vector (Addgene #140025) was first modified to insert a TruSeq sequencing adaptor (ACACTCTTTCCCTACACGACGCTCTTCCGATCT) upstream of the barcode insertion site to allow for direct amplification. Additional sequence, including a mouse U1 hairpin, was introduced downstream of the barcode site to promote nuclear translocation of RNA transcripts and more efficient SHARE-seq capture. For nuclear localization validation, lentivirus was generated (see method below), HEK293T cells were infected and sorted for a pure GFP+ culture. Plasmids will be deposited in Addgene upon publication.

Fluorescence in situ hybridization was performed using anti-GFP probes (LGC Biosearch, #VSMF-1014-5) and imaging was performed on an Andor CR-DFLY-201-40 confocal spinning disk coupled to a Nikon Ti-E microscope.

For constructing barcode libraries, the following oligonucleotides were ordered from IDT: Forward oligo:

CCTATAGTGAGTCGTATTAGAGACATNNNNCTNNNNACNNNNTCNNNNGTNNNNTGNNNNCANNNNATNNNNGCATCATCAAGATCGGAAGAGCGTCGTG

Reverse oligo:

CACGACGCTCTTCCGATCTTGATG

The two oligos were annealed with program:

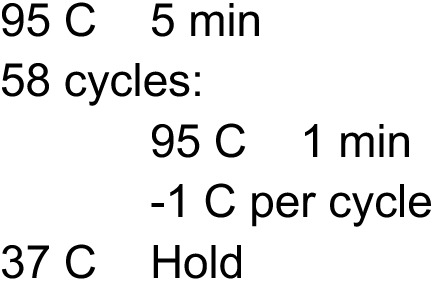

Double stranded barcode inserts were then generated. Extension was performed by adding 1 U/uL Exo-Klenow (NEB #M0212L) and 1 mM dNTPs and incubating at 37 C for 2 hours, followed by enzyme inactivation at 75 C for 20 min. The resulting annealed barcodes were purified into 20 uL of 10 mM Tris pH 8.0. The empty barcode vector (2 ug) was digested and dephosphorylated with BamHI (NEB #R3136T), XbaI (NEB #R0145S) and FastAP (Thermo #EF0652) for 4 to 12 hours at 37 C. Following purification, 1 ug of digested plasmid and 60 ng of annealed barcodes were assembled using NEBuilder 2X master mix (NEB #E2621S) for 1 to 4 hours at 50 C. The resulting product was purified into 10 uL of water, electroporated into Stbl4 ElectroMAX cells (Thermo # 11635018) and plated onto bioassay plates (Sigma CLS431111-16EA) with carbenicillin. After growth at 30C for 20 to 24 hours, colonies were scraped into media and grown for an additional 1 to 4 hours before plasmid purification.

#### Organoid lentiviral infection

Lentivirus was generated by transfecting LentiX cells with 2nd generation packaging constructs. Viral supernatant was concentrated by overnight incubation with 300 mM NaCl and 8% PEG-6000 (Millipore #8074911000) and centrifugation at 3000xg for 30 minutes. Concentrated virus was resuspended in PBS prior to cell treatment.

Murine organoids were pretreated for 24 hours with 5 μM Y-27632 before dissociation to near single cells. Cells were pre-incubated in Infection Media (WENR, 10 μM Y-27632, 10 μg/mL polybrene) for 15 minutes before addition of concentrated virus at less than or equal to 50% the volume of Infection Media. An approximate ratio of 750 μL concentrated virus to 25,000 cells was used. Spinoculation was performed by centrifuging at 600xg for 1 hour at 32 C and the cell-viral mixture was incubated at 37C for 3 to 4 hours before organoids were replated. A MOI of less than 0.3 was used and verified by GFP expression.

#### SHARE-TRACE

Nuclear isolation, fixation and transposition were performed as described above. For clonal barcode capture from SHARE-seq, barcode specific RT primer was spiked into the reverse transcription reaction at 10% general RT primer concentration: (/5Phos/GTCTCGTGGGCTCGGAGATGTGTATAAGAGACAGNNNNNNNNNN/iBiodT/CTCATTCAGCCACGG TGG)

Split-pool barcoding, ligation and reverse cross-linking were all performed without modification. ATAC-seq libraries were generated as above. During cDNA PCR amplification, a barcode specific forward primer mix was spiked into the reaction at 2 μM final concentration. The primer mix consisted of an equimolar mixture of:

ACACTCTTTCCCTACACGACGCTCTTCCGATC

ACACTCTTTCCCTACACGACGCTCTTCCGATCTNTAGACAT

ACACTCTTTCCCTACACGACGCTCTTCCGATCTNNTAGACAT

ACACTCTTTCCCTACACGACGCTCTTCCGATCTNNNTAGACAT

where N represent a random base mixture, introducing a frameshift in the amplification products and increased sequencing diversity of the barcode library

Following total cDNA amplification and purification, 0.5 uL of product was removed and further amplified with the above primer mix and P7 primer for 10 cycles and Ampure purified into 10 uL. A barcode-enriched library was then generated by amplifying 5 uL of these products with P7 and a barcode-specific index primer: AATGATACGGCGACCACCGAGATCTACAC(index5)ACACTCTTTCCCTACACGACGCTCTTCCGATCT

#### Regional colon motif accessibility

The entirety of the colon (between cecum and rectum) from a healthy control mouse was extracted as previously described and then cut into six equal length fragments (each approximately 1 cm). These fragments were then individually treated with EDTA solution, crudely scraped and dissociated to single cells as previously described. Cells were resuspended in Advanced DMEM, counted, and 10,000 cells per segment were collected. Bulk ATAC-seq and library preparation were performed in the same manner as SHARE-seq.

#### Fos CUT&Tag

Epithelium was crudely scraped from control or recovered animal colons as described above and nuclei were isolated by resuspension in HLB supplemented with PIC. Following a 5 minute incubation on ice, nuclei were diluted with Working NE Buffer - Nuclear Extraction Buffer (20 mM HEPES pH 8, 10 mM KCl, 0.1% Triton-X 100, 20% glycerol) supplemented with 0.5 mM spermidine and PIC. Following another 5 minute incubation, nuclei were centrifuged and 500K nuclei were resuspended in 100 uL Working NE Buffer. Concanavalin A beads (Bangs Laboratories, #BP531) were activated by washing twice in Bead Activation Buffer (20 mM HEPES pH 8, 10 mM KCl, 1 mM CaCl2, 1mM MnCl2) and 10 uL of beads were added per sample. Nuclei were bound to beads for 10 min at RT, magnetized and supernatant was removed. Bound nuclei were resuspended in 50 uL of Digitonin150 Buffer (20 mM HEPES, 150 mM NaCl, 0.5 mM spermidine, 0.01% digitonin, PIC) supplemented with 2 mM EDTA, 1 ug of anti-Fos antibody (AE-1059 from the lab of Dr. Michael Greenberg) was added and samples were rotated overnight at 4C.

Beads were magnetized and resuspended in 50 uL of Digitonin150 Buffer prior to addition to anti-rabbit secondary (Novus, #NBP1-72763). Samples were rotated for 30 min at RT and washed twice with Digitonin150 Buffer. Nuclei were then resuspended in 50 uL of Digitonin300 Buffer (20 mM HEPES pH 8, 300 mM NaCl, 0.5 mM spermidine, PIC, 0.01% digitonin) and 2.5 uL of CUTANA pAG-Tn5 was added (EpiCypher #15-1017). Samples were rotated for 1 hour at RT and washed twice with 200 uL of Digitonin300 Buffer. Nuclei were resuspended in Tagmentation Buffer (20 mM HEPES pH 8, 300 mM NaCl, 0.5 mM spermidine, PIC, 10 mM MgCl2) and placed on a thermocycler at 37C for 1 hour. Tagmented nuclei were magnetized, washed with 50 uL of TAPS Buffer (10 mM TAPS, 0.2 mM EDTA) and resuspended in 5 uL of SDS Release Buffer (10 mM TAPS, 0.1% SDS). Following an incubation at 58C for 1 hour, 15 uL of SDS Quench Buffer (0.67% Triton-X 100) was added and libraries were prepared using NEBNext HiFi 2x Master Mix.

#### Adenoma induction, macroscopic quantification and proliferation assessment

Mice were ordered from Jackson Laboratory (#035169) with genotype wildtype for Kras<TM4TYJ>, homozygous for Apc<TM1TNO>, homozygous for Tg(CDX2-cre/ERT2)752Erf. Adenomas were induced at 21 to 23 days following DSS cessation. For macroscopic tumors, tamoxifen dissolved in corn oil was administered at 50 mg/kg by intraperitoneal injection on three consecutive days and animals were sacrificed 25 to 28 days following the first injection. The entire length of colon was removed and imaged. Due to higher expression of the CDX2-cre driver of this mouse model in the proximal colon^57^, more tumors form in the 1 cm of colon adjacent to the cecum and adenomas were quantified in the distal 5 cm of colon for more accurate counting. Adenoma diameter was measured in ImageJ along the longest axis of each tumor and scaled to millimeters using a ruler placed in the same image. Researchers were blinded during quantification.

To measure proliferation, immunohistochemistry for Ki67 (Abcam #ab15580) was performed and proliferating fraction was quantified in only adenoma areas, as identified by H&E morphology on adjacent sections. Researchers were blinded during quantification.

#### Ex vivo adenoma organoid clonogenicity

Organoids were derived from adenoma tissue 23 days following the first tamoxifen administration. Tissue was crudely scraped following EDTA treatment and dissociated to near single cell as previously described. Cells were plated in Crypt Basal with 50 ng/mL EGF and 100 ng/mL Noggin and allowed to grow for at least one passage to purify culture. Clonogenicity (colony-forming efficiency) was calculated on the secondary organoids by plating 1,000 cells passaged from the primary organoids and assessing organoid formation 7 days after initiation of cultures.

#### Microscopic tumor initiation and quantification

Adenomas were induced 21 days following DSS cessation with a single dose of tamoxifen at 10 mg/kg and sacrificed 13-14 days following. The entire length of colon was removed, fixed and paraffin embedded as a swiss roll as previously described. Immunohistochemistry for beta-catenin (BD, #610153) was performed. The entire section was first imaged at low magnification (2x) to quantify total tissue area, then microscopic tumors were identified by high nuclear beta-catenin staining and imaged under high magnification (20x). Both total tissue and individual tumor areas were quantified using ImageJ, using scale bars as reference, and tumor area summed across all tumors and reported as percent of total tissue. Researchers were blinded during imaging and quantification.

#### Spatial transcriptomics on adenoma tissue

Spatial RNA-sequencing was performed on fresh frozen swiss rolls of colons with adenomas induced at 50 mg/kg x 3. The Visium HD Kit was used according to standard protocol with the following modifications: the OCT-embedded swiss rolls were sectioned on a cryostat at 10 μm thickness and mounted on a Fisherbrand™ Superfrost™ Plus glass slide. The slides were fixed using 4% PFA and stained using H&E method. During that process the wash buffers were supplemented with either ribonucleoside vanadyl complex or a commercial RNAse inhibitor. After imaging of the H&E staining, the samples were destained and permeabilized using 1% SDS and then pre-chilled 70% methanol. Tissue processed this way was then analyzed using a standard 10X Visium HD method described in the User Guide CG000685 (Rev A).

### Data Processing and Analysis

#### SHARE-seq raw data processing

Raw SHARE-seq data was processed as previously described^24^ with minor modifications (code available at https://github.com/masai1116/SHARE-seq-alignmentV2/). Briefly, raw fastq files were demultiplexed using custom Python scripts. ATAC-seq reads were aligned to the mm10 or hg38 genome using bowtie2^75^, removing fragments with length longer than 2 kb. RNA-seq reads were aligned using STAR^76^, removing reads with greater than 20 alignments or score less than 0.3. Both library types were further filtered to remove mitochondrial reads and reads with MAPQ < 30, and chrY reads were removed from ATAC libraries. Filtered ATAC reads were then deduplicated and further filtered for cell barcodes with at least 100 raw reads. Filtered RNA reads were assigned to genes using featureCounts^77^ using only primary mapping coordinates and UMIs were counted using umi_tools^78^, removing those consisting of only G’s. Libraries were then filtered for RNA cell barcodes with at least 300 UMIs or ATAC cell barcodes with a library size of 500 reads.

#### Single cell RNA-seq processing

All filtered cell barcodes were normalized by the total number of transcripts detected. The top 5000 most variable genes were selected and PCA was performed on the log2+1 transformed values of these genes. Library sizes were smoothened over the 20 k-nearest neighbors in this space and clumps of cells were manually identified as those barcodes with extremely high smoothened library sizes. Once clumps were removed, the remaining barcodes were processed with scrublet^79^ to identify doublets and barcode collisions, where doublet score thresholds were manually selected. The remaining singlet filtered barcodes were again normalized, PCA was performed again as described and the resulting top 20 PCs were batch corrected using harmony^80^. Corrected PCs with high correlation to barcode library size were removed prior to UMAP and k=5 k-NN analysis for Louvain graph-based clustering. Gene expression was visualized after using k=5 k-NN for smoothening normalized values and capping maximum z-score values at 3.

#### Single cell ATAC-seq processing

Peaks were called on a merged set of fragments from all sub-libraries for each dataset (in vivo tissue memory, ex vivo organoid culture or adenoma tissue) using MACS2^81^. Fragments were counted within peaks for each cell barcode and barcodes with low library size (in vivo tissue memory and ex vivo organoids 1000 fragments, adenoma tissue 2000 fragments) or fraction of reads in peaks (in vivo tissue memory and adenoma tissue FRIP < 0.2, ex vivo organoids FRIP < 0.25) were removed. cisTopic^28^ models were generated on all filtered cell barcodes for 10*n topics for n = 1 to 9, with 150 iterations and burnin=120. Library sizes were smoothened over the 20 k-NN in this space and clumps of cells were manually identified as those barcodes with extremely high smoothened library sizes. These clump barcodes, as well as clump and doublet barcodes identified in the matched RNA cell barcodes, were removed to identify singlet cell barcodes. Using the value of n selected on all cell barcodes, cisTopic models were generated again on singlet barcodes using 10*(n-1), 10*n, 10*(n+1) topics. UMAP, k=5 k-NN and Louvain graph-based clustering analysis were performed in the resulting topic space. Gene expression visualization was performed by matching ATAC cell barcodes to corresponding RNA cell barcodes to get smoothened normalized expression values. For genome track visualization, fragments were separated by colitis stage, normalized to 10 million total reads and Tn5 insertion sites were counted.

#### Cell type identification

For each scATAC-seq Louvain cluster, the fraction of cells expressing each marker gene and the average normalized expression within those cells was computed and plotted. Clusters with high expression of *Ptprc* or *Acta2* were designated as non-epithelial cells. Clusters with high expression of *Muc1*, *Chga/b* or *Dclk1* were designated as secretory cells and subdivided as goblet, enteroendocrine or tuft cells when possible. In non-neoplastic tissue, the clusters with high expression of *Lgr5* and *Lrig1* were assigned as stem/progenitor cells and those with high expression of *Car4*, *Car1*, *Lypd8* or *Aqp8* were assigned as differentiated absorptive enterocytes. The remaining clusters with moderate expression of those genes were assigned as intermediate absorptive enterocytes. Analogous assignment was performed for ex vivo organoid clusters. For adenoma experiments, the clusters with high expression of *Axin2* and high motif scores for Lef1, in conjunction with high expression of *Lgr5*, *Lrig1* and *Mki67*, were assigned as adenoma cells. The remaining epithelial non-secretory cells were then partitioned as described for non-neoplastic tissue.

#### Gene expression change analysis

For differential analysis, raw UMI counts were first pseudobulked by cell type in each animal then tested using DESeq2^82^. For *in vivo* memory analysis, genes were filtered for those with a minimum reads per million (RPM) of 10 in at least one pseudobulk, then each disease stage was compared to controls and an adjusted p-value of 0.05 was used for significance. Log2(fold change) values were taken from DESeq2 output. For adenoma gene activation, all adenoma pseudobulks were tested against all stem/progenitor pseudobulks, regardless of colitis condition. When plotting individual genes, raw UMI counts were pseudobulked across stem/progenitor cells within each animal and RPM values were calculated using total assigned UMIs. Z-scores were calculated across all animals for each gene and change was calculated by subtracting average value across control pseudobulks. Gene ontology analysis^83^ was done with all differential genes.

#### k-NN enrichment analysis

For the *in vivo* memory dataset, cells were subsetted to stem/progenitors only and 100 k-NN were determined in the cisTopic space defined above. For each condition (control, acute injury, chronic injury, recovery), the expected value of k-NN was calculated for random assignment to the 100 k-NN and therefore was the fraction of cells in each condition. Enrichment for each condition was then calculated as the (observed % k-NN) - (expected % k-NN). The analogous procedure was used for human organoids except using all cells.

#### Motif accessibility analysis

Peaks were annotated as containing a motif using motifmatchr for known cisBP^84^ motifs. For *de novo* derived motifs, peak annotation was performed as described below under the “Footprinting and seq2PRINT” method section. Single cell motif deviations and accessibility scores were calculated with chromVAR^29^ using 250 background peaks across all single cells within each dataset (in vivo memory, ex vivo organoids or adenoma-induced tissue). A motif similarity matrix was calculated on all known and de novo motifs using Tomtom^85^ and a q-value cutoff of 0.05 was used to group motifs into families based on sequence similarity. This step ensures reliable motif-motif comparisons for downstream analysis. During the bagging process, motifs are sorted based on their variation across cells and those with highest variation were retained as “leaders”, while other motifs with high similarity scores to these representatives are merged into their respective “families,” effectively consolidating similar motifs into unified groups. These motif families were created using the in vivo tissue memory dataset and held constant across all other murine tissue and organoid experiments. Motif families were derived analogously in the human organoid dataset.

For motif accessibility testing, the single cell scores were averaged across all cells of a given type within each animal or all cells in an organoid line. For in vivo memory, p-values were computed for the 50 most variable families by t-test and adjusted using the Benjamini-Hochberg method. Motif accessibility change was defined as the difference between the score for each replicate and the average across control animals or organoid lines. When visualizing the *in vivo* change per mouse as a heatmap, samples with less than 200 stem cells were excluded.

Heterogeneity in single cell motif accessibility was evaluated by randomly downsampling each condition to 500 stem/progenitor cells and computing p-values for selected motifs with a Kolmogorov–Smirnov test. Activated cells were defined as stem/progenitor cells with a score greater than 1.5 and the fraction of activated cells was calculated for each animal using all stem/progenitor cells.

#### SHARE-TRACE clone assignment

Demultiplexed reads were processed with custom python scripts to search for common barcode vector sequence TAGACAT, allowing at most 1 mismatch. This sequence and all preceding base pairs were trimmed, and any reads with UMIs consisting of only Gs were removed. The remaining 48 bp of barcode sequences were validated by checking for staggered invariant sequences every 4 bases (CT,AC,TC,GT,TG,CA,AT,GC), removing off-target PCR products. To account for sequencing base call errors, the number of reads for each cell-UMI-barcode triple were counted and those with less than 5 reads were removed.

To identify clonal barcode sequences, a Levenshtein distance matrix was calculated across all remaining barcode sequences. For each barcode sequence, all other sequences within a distance of 4 were found and the barcode was assigned to the most abundant of those sequences. Distance between members of this set of most abundant sequences was computed once more and any sequences within a distance of 2 were collapsed to the more abundant sequence, generating a consensus set of clonal barcodes. The remaining reads were used to assign each cell-UMI pair to a clone.

To account for transcript mixing that occurs during SHARE-seq split-pooling, we leveraged the fact that each clone should be unique to organoids generated from a single mouse. Cell-clone assignments were matched to validated ATAC or RNA singlets and each clone was assigned to the animal from which the most cells were present (typically >95% of barcode reads). Only clones with at least 5 cells were used for subsequent analysis.

#### SHARE-TRACE clonal variance calculation

For each feature (motif families or gene programs), the standard deviation of the single cell scores were calculated across all cells in a clone and the observed clonal variance was then defined as the median of these values across clones to the second power. Cell-clone assignments were then randomly permuted and the shuffled clonal variance was computed analogously. This process was repeated 1000 times and the mean and standard deviation of the resulting distribution of randomized clonal variance was used to compute a p-value: Z = (observed_clonal_variance - mean_shuffled_clonal_variance) / (stdev_shuffed_clonal_variance) P-value = 2*pnorm(-abs(Z))

The FDR was then calculated using the Benjamini-Hochberg method of p-value adjustment.

For comparisons between colitis and control organoids, the standard deviation and median value of the scores for all cells belonging to each clone was computed. Variance was compared between conditions by performing a Kolmogorov–Smirnov test on the standard deviations. To compare change in clonal score and variance, the medians of those values were then computed across clones and the value for control clones was subtracted from colitis clones. Clones with high AP-1 accessibility were defined as those with a median score > 1.25.

#### Gene expression program derivation and scoring

For gene expression program modeling, the input was a cell-by-gene count matrix. Unlike the cistopic approach for scATAC-seq data^28^, this matrix was not binarized but remains in raw count format. Latent Dirichlet Allocation (LDA), implemented in the Mallet package, was used to infer: 1)the probability distribution of topics for each cell, and 2)the probability distribution of genes for each topic. LDA models were trained with a range of topic numbers (30–90), and the model with the highest log-likelihood was selected, following a procedure similar to cisTopic. To score single cells on these programs, we adapted the chromVAR algorithm for RNA topics. The input cell-by-peak accessibility matrix was replaced with the cell-by-gene transcription matrix, and the motif-by-peak matching matrix is substituted with the topic-by-gene probability distribution matrix inferred by LDA. The rest of the calculations remain identical to the original chromVAR workflow. Background genes were generated by grouping all genes in 20 bins of equivalent size based on average expression and 250 background genes were chosen for each gene in the annotated set. The resulting cell-by-topic scores represented the activity levels of RNA topics in each cell while controlling for sequencing depth, gene expression level, and other biases.

To identify AP-1 and HNF4/PPAR associated gene programs, the mean motif score and gene program score across all cells within each clone was calculated. These mean values were then correlated across clones to get motif-gene program correlation values. The top programs were selected for each motif family (gene program 20 for AP-1, and gene programs 9 and 30 for HNF4/PPAR) and the top 150 genes by weight of contribution to the program were selected for Gene Ontology analysis and subsequent gene program scoring. For plotting scores on UMAP projections, single cell score values were capped at -3 and 3.

#### Footprinting and seq2PRINT

Footprint scores were calculated using the scPrinter package^48^. The data used were as follows: in vivo memory - all epithelial cells; AP-1 inhibition - all bulk ATAC-seq reads; human organoids - all cells; mouse adenomas - only adenoma cells. Briefly, the seq2PRINT model was trained to take DNA sequences spanning a cCRE of interest and its surrounding regions (+/- 920 bp) as input and predict multiscale footprints derived using the PRINT method. After training, DeepLIFT is employed to extract sequence attribution scores, which represent the contribution of each base pair in the input sequence to the predicted footprints. These sequence attribution scores enable highly accurate TF binding predictions through a neural network model trained on ChIP-seq data^48^ and are referred to as seq2PRINT footprint scores. In this study, raw seq2PRINT footprint scores were binned at a 10-bp resolution for all cCREs. Bins with maximum seq2PRINT footprint scores below 0.2 across all conditions are excluded from further analysis. To facilitate comparisons across different conditions, the scores were quantile-transformed for each condition using the quantile_transform function in scikit-learn (n_quantiles=100000, target_distribution=’uniform’).

Sequence attribution scores from seq2PRINT were further analyzed to identify de novo motifs. TF-MoDISco is used to align and cluster seqlets (local sub-sequences with high sequence attribution scores) into groups of de novo motifs. To assign these motifs to candidate cis-regulatory elements (cCREs), the software finemo was used, which takes the output de novo motifs from TF-MoDISco and the sequence attribution scores as input for motif matching.

Motif accessibility change in *de novo* motifs was calculated as described above, where single cells scores were averaged across all stem/progenitor cells in a given animal, p-values computed by t-test as compared to control animals and change was defined as the difference from the control animal average.

To visualize how the seq2PRINT model learns the association between input DNA sequences and output multi-scale footprints, we generated marginalized predictions (referred to as delta effects) for given motifs using the delta_effects_seq2print function from the scprinter psackage. Briefly, we randomly selected 10,000 CRE sequences, inserted the consensus sequence of a specific motif at the center, and averaged the difference in model predictions with and without the inserted motif across the 10,000 CREs.

#### AlphaFold3-based structure prediction

The full amino acid sequences and features of murine Fos (P01101), Jun (P05627), Foxp1 (P58462), Foxa1 (P35582), Foxn2 (E9Q7L6), and Foxj2 (Q9ES18) were downloaded from UniProt. Structures were predicted using Alphafold3 through Alphafold Server by combining nucleotide sequences and amino acid sequences. Each structure included the motif nucleotide sequence, its reverse complement, and the amino acid sequences of Fos, Jun, and Fox proteins. PyMOL (ver. 3.1.3) was used to select for interacting residues and structure visualization. To reduce the amount of disordered regions in each structure and to focus on protein-to-protein and protein-to-DNA interactions, the beginning and end of the protein sequences were truncated to only include regions containing residues within 3.5 Å of neighboring protein and DNA structures. Alphafold3 was once again used to predict the truncated structures containing the complete DNA sequence, truncated Fos chain, truncated Jun chain and truncated Fox chain. Fox residues within 3.5 Å of either Fos or Jun were highlighted in the truncated structures as an estimate of interaction between proteins. The location of the interaction residues were compared to known features for each Fox protein. For visualization purposes, structures were modified to only include regions with a b-factor greater than 50.

#### CUT&Tag processing and analysis

Fos CUT&Tag reads were aligned to the mm10 genome annotation and processed analogously to SHARE-seq (ignoring the single cell barcode considerations). Fos binding signal analysis was performed by counting CUT&Tag reads within ATAC-seq peaks called from SHARE-seq data and summing reads across replicates for control or recovered samples. Reads per million (RPM) was calculated at each peak and variable peaks were identified as the top 10,000 by standard deviation across RPM values after filtering for peaks with a minimum of 20 raw reads in at least one sample.

For footprinting prediction performance, peaks were first called on Fos CUT&Tag data using MACS2. Reads were counted within peaks, normalized to 10 million total reads and a t-test was performed between recovered and control samples. Differential peaks were defined as having a p-value of less than 0.05. Normalized footprint scores were then condensed by motif, where overlapping AP-1 motifs were collapsed to one site and change in footprint score was calculated between recovered and control. Performance was then evaluated by calculating the sensitivity and specificity of identifying differential peaks based on varying thresholds of footprint score difference.

#### Co-binding Analysis

Change in footprint score was calculated as the difference between acute, chronic or recovered scores and their batch matched controls. Memory footprints were defined as those showing a minimum change of 0.2. Using all pairs across the selected families, peaks were labeled as containing a memory motif for the first family only, second family only, both families or neither family and an odds ratio of the resulting contingency table was calculated. Co-binding scores were then calculated by log2 transforming these odds ratios, performing 10th-90th quantile normalization across all pairs on positive and negative ratios separately, and linearly scaling positive values between 0 and 1 and negative values between 0 and -1. For primary tissue memory, this normalization and scaling was performed on values across all timepoints. For human IBD organoids and mouse adenomas, footprint differences were calculated as (IBD - Control) and (Recovered adenoma - Control adenoma), respectively. For AP-1 co-binding loss, the analysis was performed analogously using a decrease in footprint score of 0.2 (T-5224 - DMSO). Distance between TFs was calculated by finding the midpoints of all memory sites and calculating all distances between sites within the same peak.

Motif families were identified as above under “Motif accessibility analysis” with the following modifications: ETS - ETS family with Spi1, Spib, and Spic added.

RUNX - Runx1, Runx2, Runx3.

SNAI/MESP - Snai2, Snai3, Mesp1, Mesp2, Tcf3, Tcf4.

ESRR - Esrra, Esrrb, Esrrg.

Retinoid - Rxra, Rxrb, Rxrg, Rara, Rarb.

#### Visium HD data processing and visualization

Raw BCL files were converted to fastqs using spaceranger mkfastq (v3.0.1) and gene expression values were computed using spaceranger count using the Visium_Mouse_Transcriptome_Probe_Set_v2.0 and mm10 genome reference data. Bins of size 16 μm were used. For plotting of individual genes, beads were first filtered for a minimum of 300 reads, and then normalized to average read depth across all remaining beads. PCA was performed on the log2+1 transformation of these values and the top 20 PCs were used to find k-NN (k=20) for smoothening prior to plotting of Z-scored expression values (capped at -3 and 3).

#### Spatial transcriptomic adenoma identification and gene scoring

To identify tumor cells, the PC smoothened *Axin2* expression values were Z-scored and adenoma cells were defined as beads with a minimum value of 1. Tumors were then called by finding k=5 k-NN using spatial coordinates of adenomas cells only and performing Louvain graph-based clustering.

For AP-1 associated genes, the top 150 genes from program 20 were overlapped with probes present in the Visium output and then this set of genes was scored analogously to single cells as above, treating bins as cells. In single bin spatial plots, scores were plotted without smoothening and capped at -2 and 2. For whole tumor analysis, raw expression values were pseudobulked by tumor call and these pseudobulks were scored analogously as cells. Tumors with high AP-1 associated gene expression were identified as those with scores > 1.5. For analysis of single AP-1 related genes, reads per million (RPM) values were computed on single bins. Visualization of these genes in adenomas only was performed by subsetting bins to only tumor cell calls, renormalizing and resmoothening expression values in only these cells, smoothening once more across the k=20 x-y coordinate k-NN and capping Z-scores at -3 and 3 before plotting.

## Extended Data

**Extended Data Figure 1.**
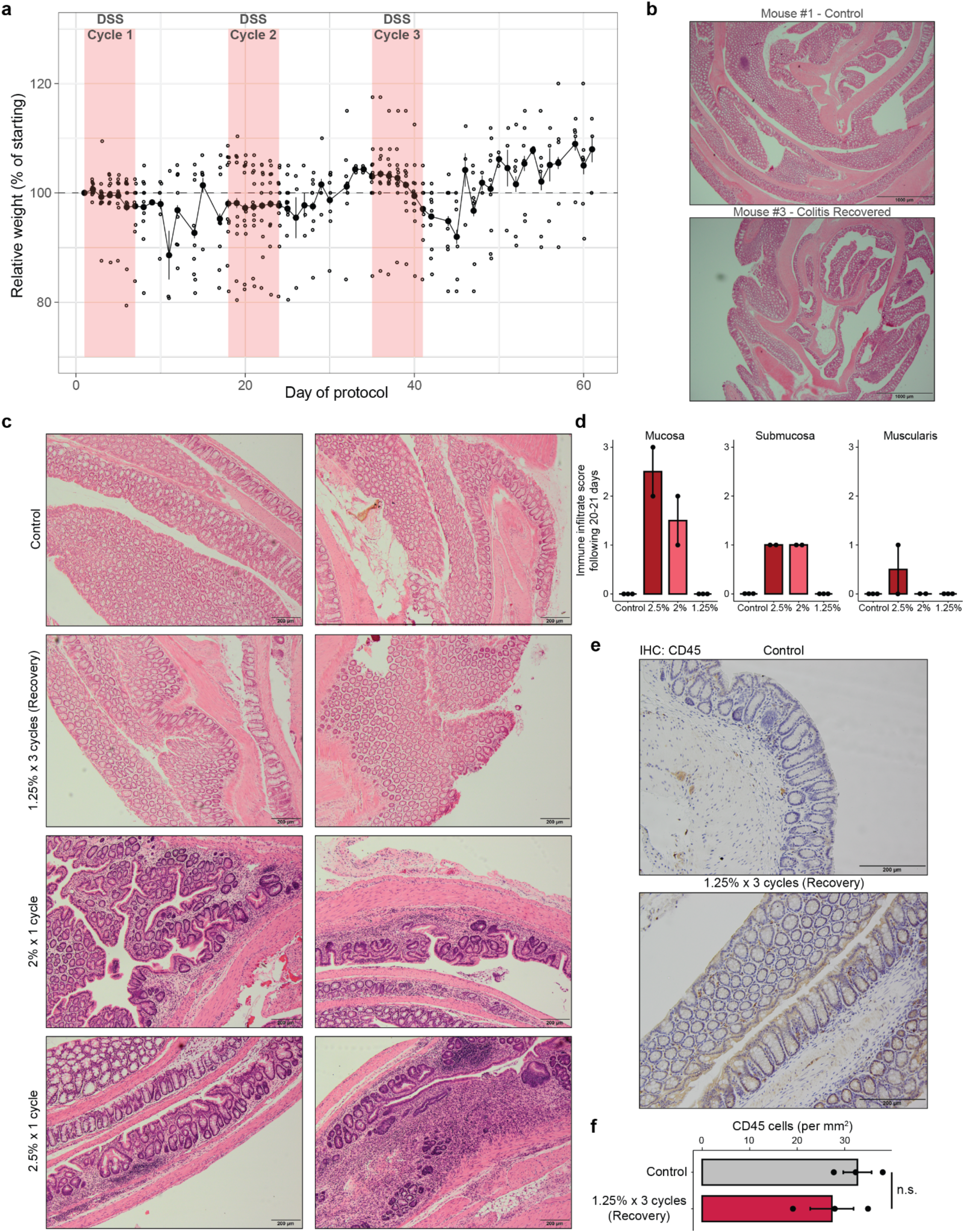
Tissue and organismal recovery following colitis. a, Mouse weights, normalized to day 1 weight, through chronic colitis and recovery. Each point represents an animal. b, Colon swiss rolls of mice 21 days following three cycles of 1.25% DSS (Recovery timepoint) and healthy controls. Scale bars 1 mm. c, Representative images of distal colon immune infiltration in healthy controls, 21 days following three cycles of 1.25% DSS, 20 days following one cycle of 2% DSS, and 20 days following one cycle of 2.5% DSS. Scale bars 200 μm. d, Colitis grading of distal colon immune infiltration per Remke et al.^74^ Each point represents tissue from one animal. e, Representative images of CD45 immunohistochemistry in the distal colon 21 days following three cycles of 1.25% DSS. f, Quantification of CD45 cell density in the mucosa and submucosa of the distal colon at 21 days of recovery. All error bars are s.e.m. (*) p < 0.05, (**) p < 0.01, (***) p < 0.001

**Extended Data Figure 2.**
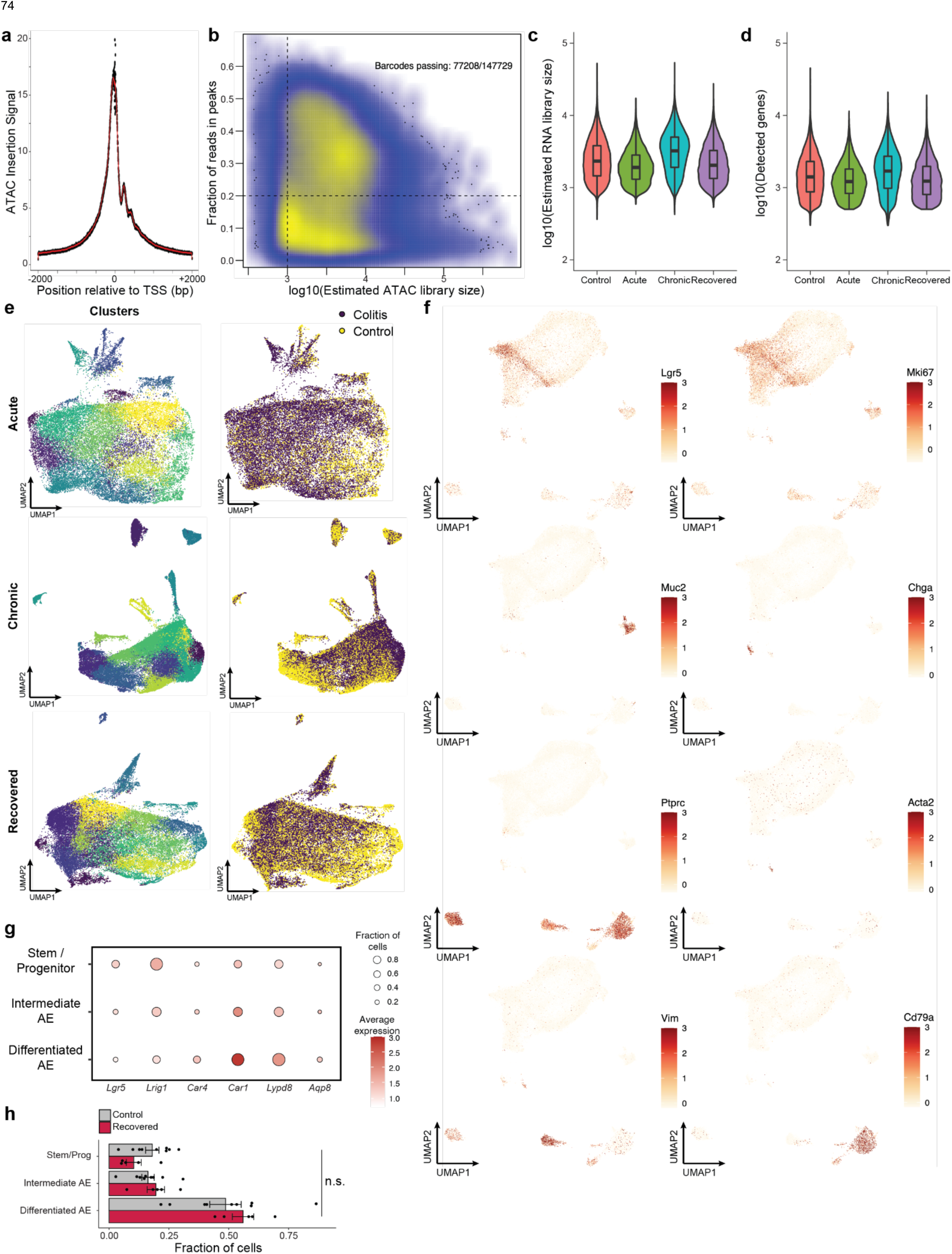
Quality control and identification of cell types. a, TSS enrichment of scATAC-seq data. The x-axis represents distance from the center of all transcription start sites and the y-axis shows signal normalized to -2 kb. b, Density plot of library size vs FRIP for all scATAC-seq barcodes. Barcodes passing filter indicate number before matching with scRNA-seq data. c, Estimated library size per cell by sample for scRNA-seq data. d, Number of unique genes detected per cell for scRNA-seq data. e, UMAP embedding of scRNA-seq data per timepoint. Each colitis sample is co-embedded with its corresponding batch controls. Left, cells colored by clusters called. Right, cells colored by sample condition. f, Key marker gene expression shown on scATAC-seq UMAP embedding. Relative expression was computed for each gene independently. g, Marker gene expression for each cell type identified within the absorptive lineage. For each gene, the size of the circle indicates the fraction of cells in which at least one transcript was detected and the color of the circle indicates average expression within the cell type. h, Absorptive epithelial cell type distribution between colitis recovered and control mice. Each circle represents one animal. All error bars are s.e.m.

**Extended Data Figure 3.**
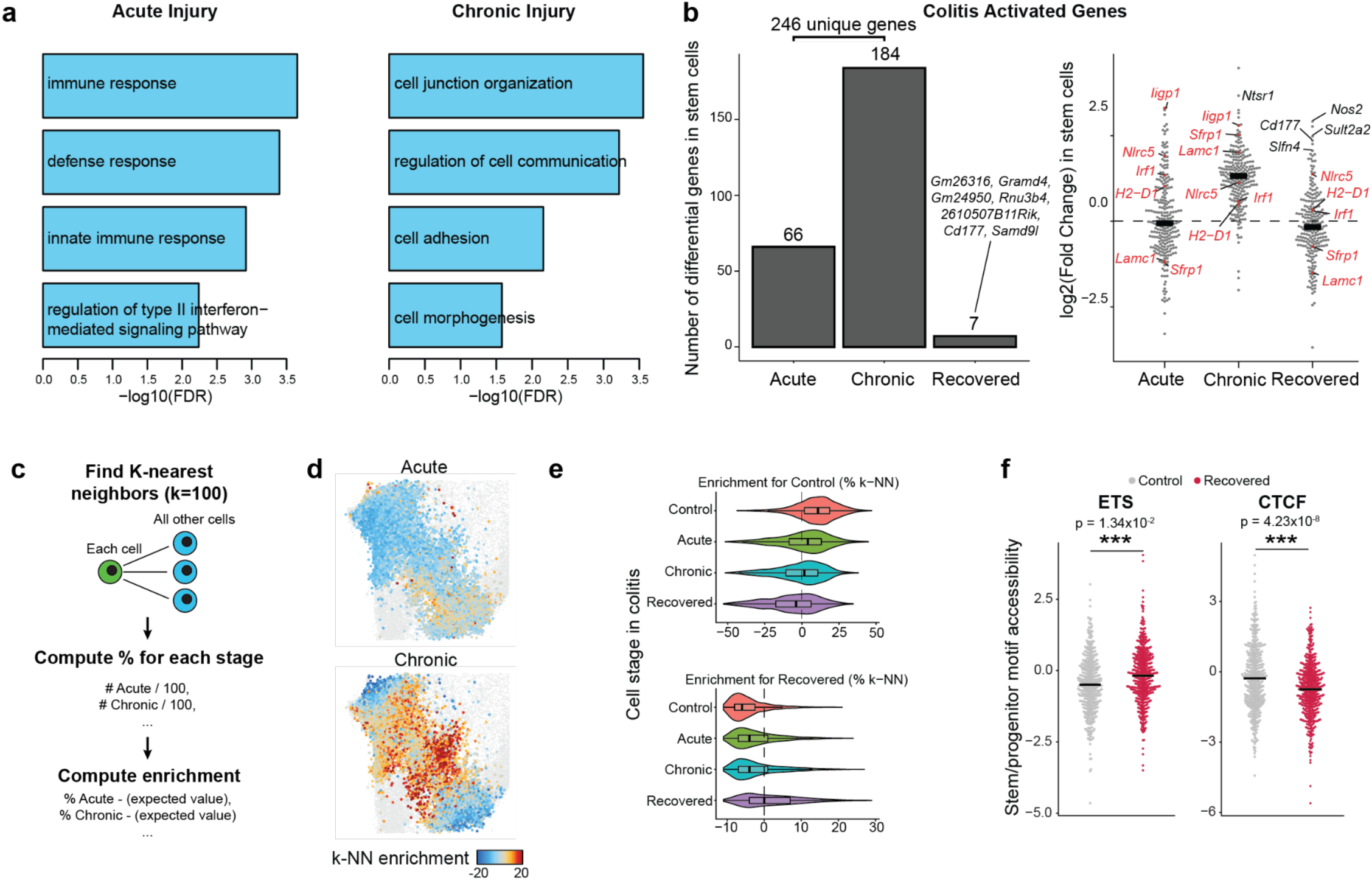
Cellular memory following chronic colitis. a, Gene ontology for genes upregulated in stem cells at acute or chronic injury. B, Change in stem cells for all upregulated genes during acute or chronic injury. Left, number of significant genes relative to controls. Right, fold change relative to controls where each point represents a gene. c, Schematic for computing neighborhood enrichment by sample. d, UMAP embedding showing enrichment for each colitis stage in k-NN networks, with each point representing a single cell colored by the enrichment of its k-NN network for a given stage. e, k-NN stage enrichment for all stem cells. The y-axis represents the stage of colitis from which the cell was acquired and the x-axis represents the enrichment of either control or recovered k-NN. f, Single cell motif accessibility for select motif families. Each point represents a single stem cell.

**Extended Data Figure 4.**
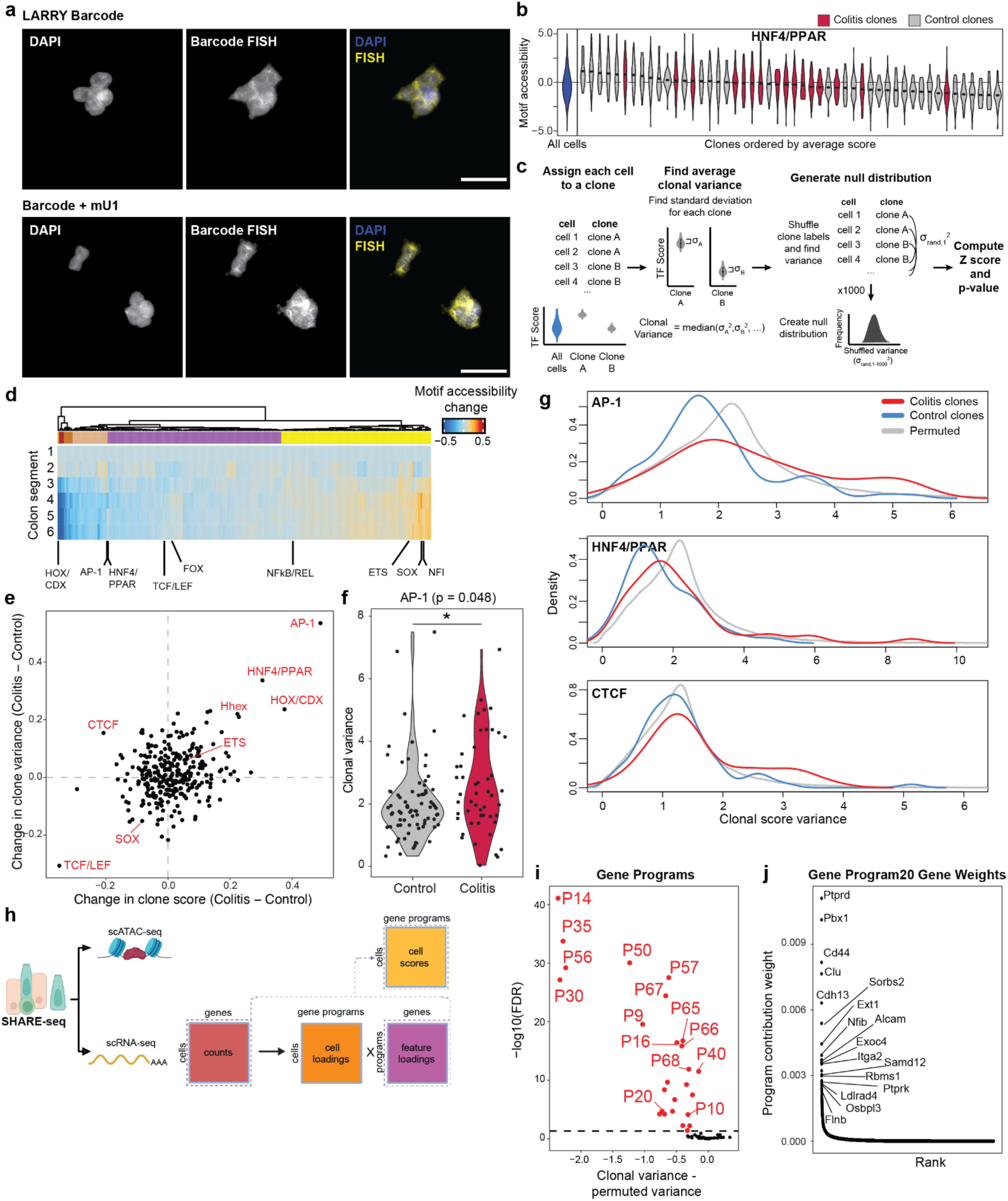
Clonal inheritance of epigenetic states. a, Fluorescent in situ hybridization for clonal barcode transcripts in HEK293T cells. Top, cells infected with original LARRY barcode from Weinreb et al.^42^ Bottom, cells infected with LARRY barcode modified by addition of mU1 hairpin sequence. Scale bar 50 μm. b, Distribution of motif accessibility in representative organoid clones for HNF4/PPAR. The y-axis represents motif score and x-axis shows clones ranked by average score. c, Framework for computing clonal variance and identifying clonal features. d, Accessibility of all TF motif families relative to position in the colon. Segment 1 is colon adjacent to cecum and segment 6 is adjacent to rectum. e, Change in clonal signal and variability of TF motif accessibility between control and colitis organoids. The x-axis represents the difference between average motif score across clones and the y-axis the difference between average motif variance across clones. f, Distribution of AP-1 motif score variance in clones. Each point represents a single clone. g, Distribution of motif score variance in clones as compared to random permutation of clonal identity. h, Schematic for deriving gene expression programs. i, Clonal memory of gene programs. j, Individual gene contribution weights to program 20. (*) p < 0.05, (**) p < 0.01, (***) p < 0.001

**Extended Data Figure 5.**
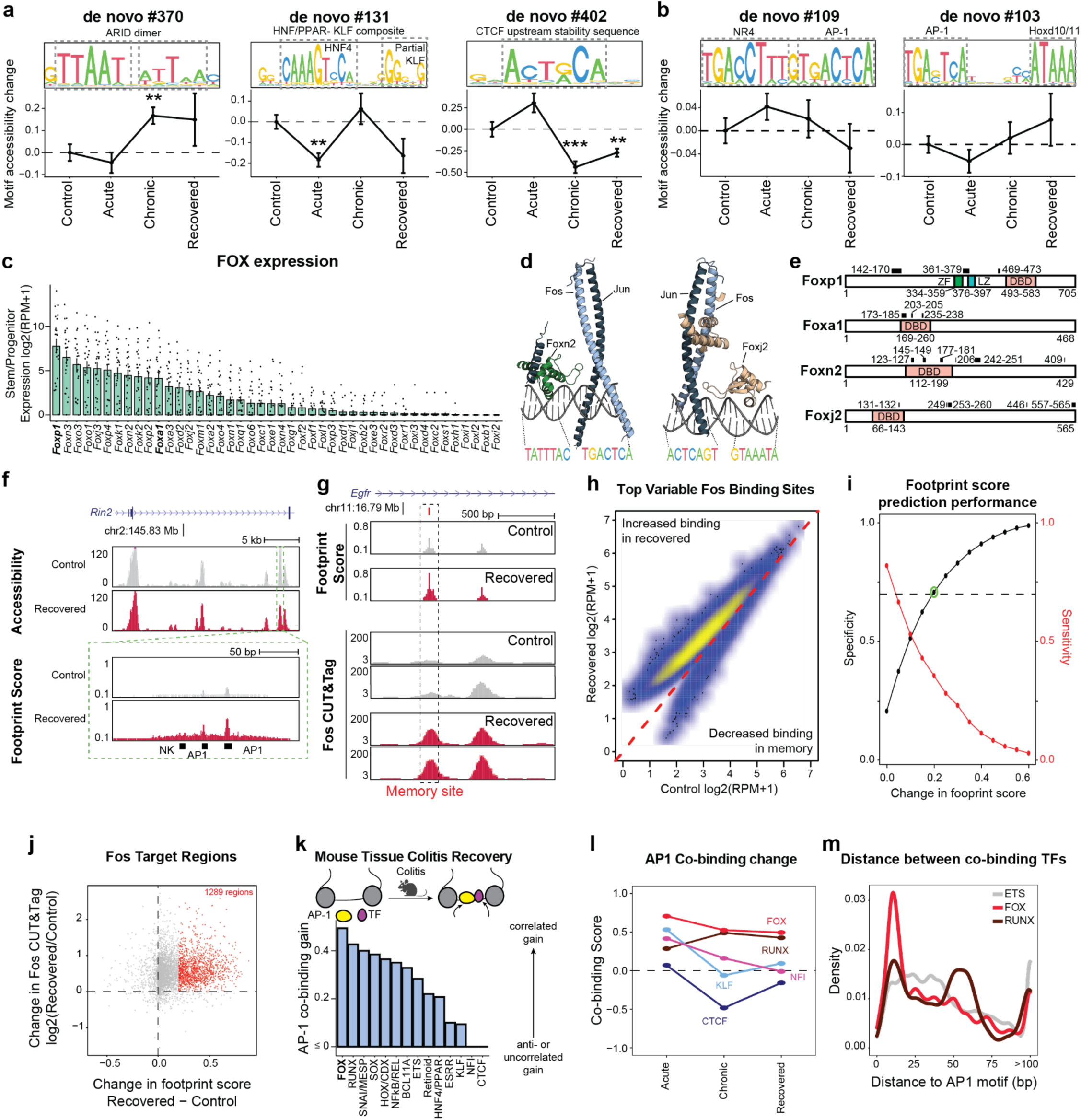
Transcription factor networks in colitis memory. a, Examples of *de novo* derived motifs. The y-axis on motif sequences show information content. Line plots show change in motif accessibility in stem cells through colitis progression. Each point represents average accessibility in stem cells across all animals. b, AP-1 compositie motifs. c, Gene expression of FOX family transcription factors in all stem cells in primary tissue. Each point represents RPM value after pseudobulking by mouse. d, AlphaFold predicted structures for Fos-Jun dimer, composite motif and either Foxn2 or Foxj2. e, UniProt domain annotation for Foxa1 and Foxp1 with black boxes indicating regions predicted to interact with AP-1. Numbers indicate amino acid positions. DBD = DNA binding domain, ZF = Zinc finger, LZ = Leucine zipper. f, Genome tracks showing high resolution footprinting in epithelial cells. Horizontal position represents genomic location and vertical height represents either ATAC-seq accessibility (top) or footprint score (bottom). g, Genome tracks of a Fos memory site showing footprint score and respective Fos CUT&Tag. h, Fos CUT&Tag signal in control and recovered tissue at the top 10,000 variable sites following normalization to read depth. i, Differential footprint score performance in predicting CUT&Tag signal. The x-axis represents varying thresholds for difference in AP-1 footprint score andb the y-axis shows sensitivity and specificity for detecting Fos CUT&Tag signal difference between control and recovered tissue. j, Comparison of AP-1 footprint score difference and Fos CUT&Tag signal difference at all identified Fos binding sites. Red points represent sites called as different by footprint score difference of 0.2. k, Co-binding scores with AP-1 factors in primary epithelial tissue following colitis recovery. l, TF family co-binding change across colitis progression. m, Distance from each TF family memory footprint to the nearest AP-1 memory footprint. All error bars are s.e.m. (*) p < 0.05, (**) p < 0.01, (***) p < 0.001

**Extended Data Figure 6.**
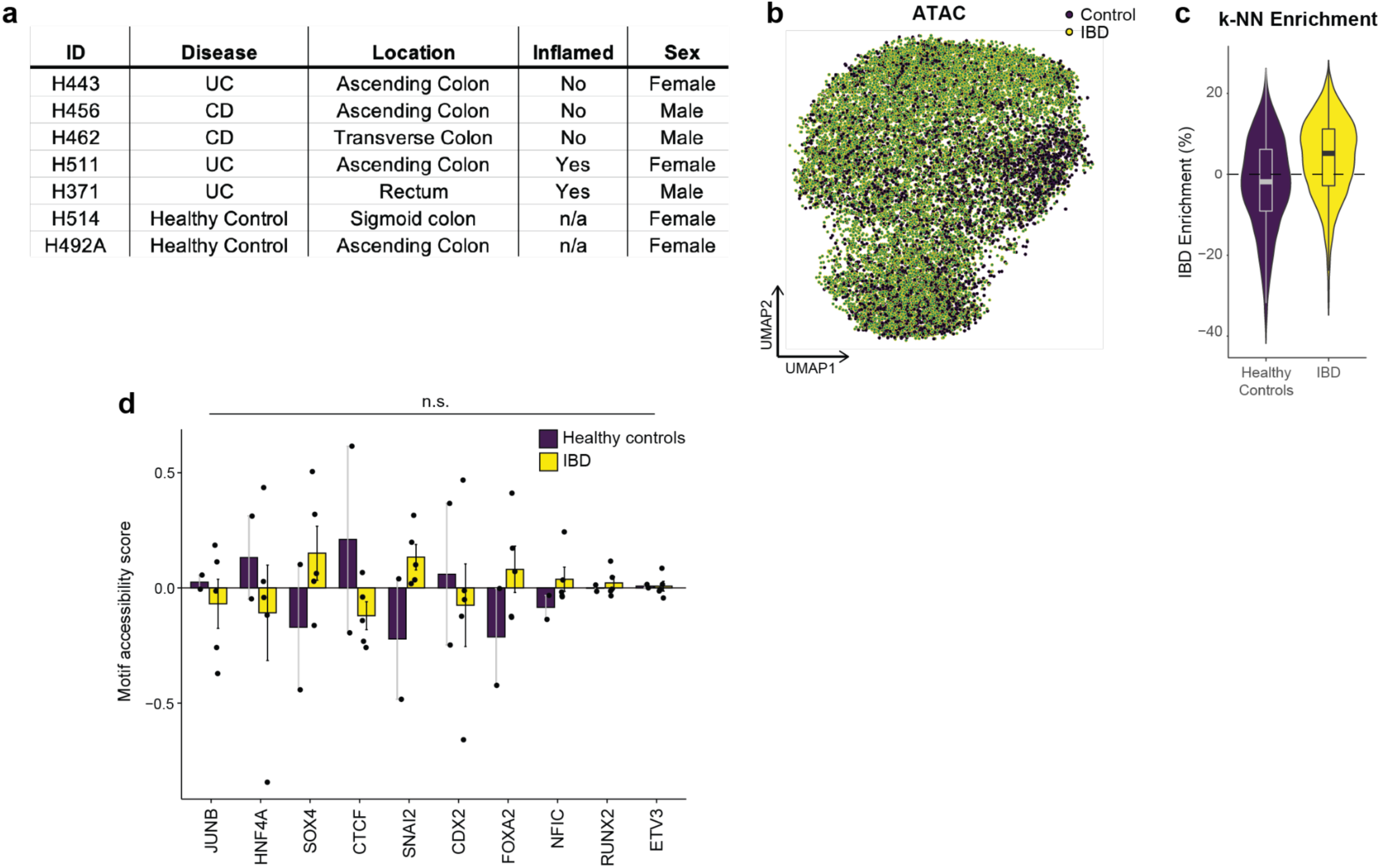
SHARE-seq in organoids derived from inflammatory bowel disease patients. a, Clinical information for each patient-derived organoid line b, UMAP embedding of scATAC-seq data colored by patient disease state. c, Distribution of k-NN network enrichment for IBD cells. d, Motif memory in patient-derived organoids. Each point represents the average motif score per organoid line. All error bars are s.e.m.

**Extended Data Figure 7.**
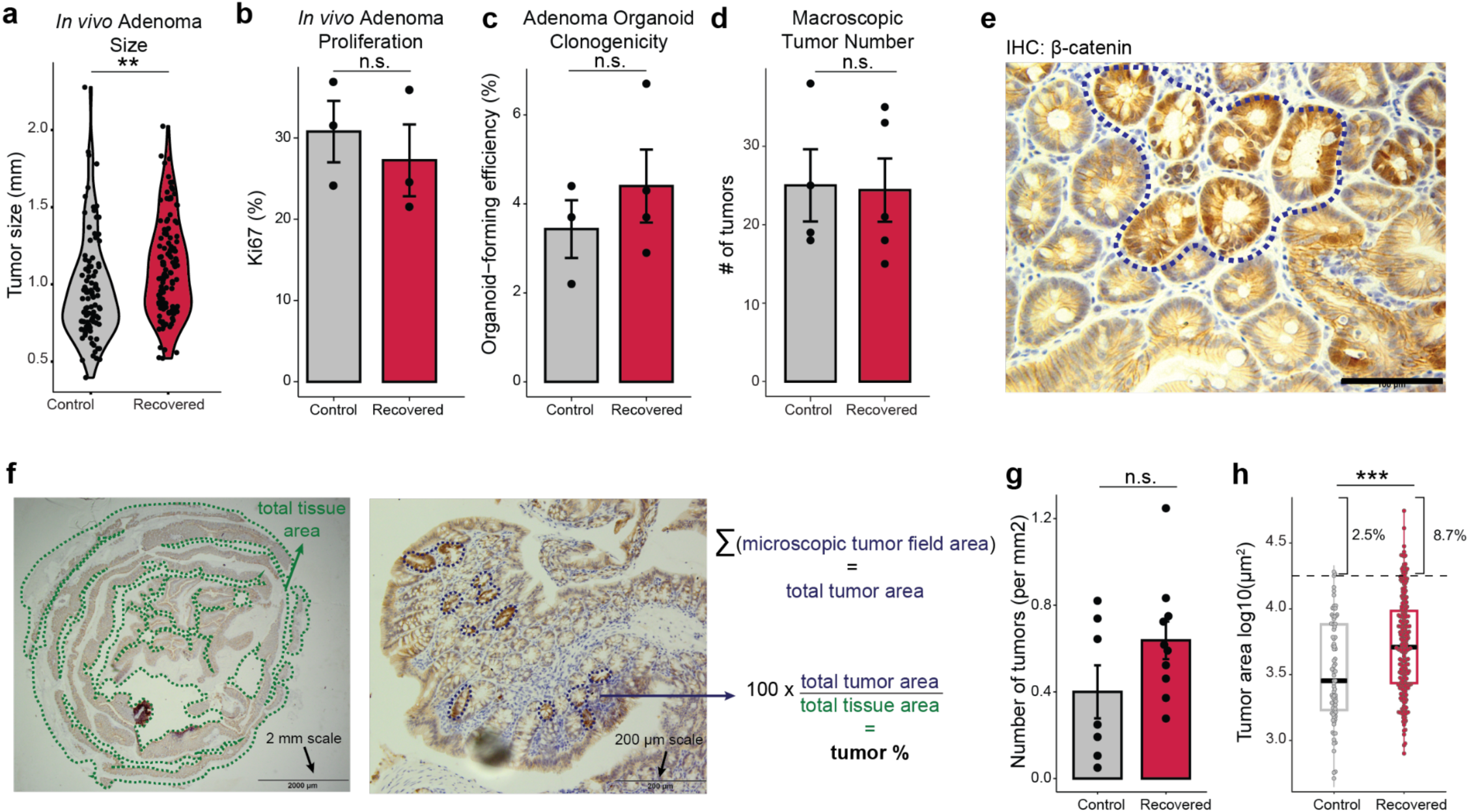
Features of adenoma following colitis. a, Distribution of macroscopic adenoma size between control and colitis recovered animals. Each point represents a single tumor. b, Proliferation in adenomas following high dose tamoxifen induction (50 mg/kg x 3). Each point represents a mouse. c, Clonogenicity in adenoma-derived organoid lines. Each point represents an organoid line. d, Visible adenoma number following high dose tamoxifen induction. Each point represents a mouse. e, Representative example of microscopic tumor following low dose tamoxifen induction (10 mg/kg x 1). Scale bar 100 μm. f, Method for computing tumor area. Left, low magnification image used to compute total tissue area. Middle, high magnification image where individual tumors are individually traced. Right, equation for computing percent tumor area. g, Microscopic tumor number normalized to total tissue area. Each point represents a mouse. h, Area of individual tumors. Each point represents a tumor. All error bars are s.e.m. (*) p < 0.05, (**) p < 0.01, (***) p < 0.001

**Extended Data Figure 8.**
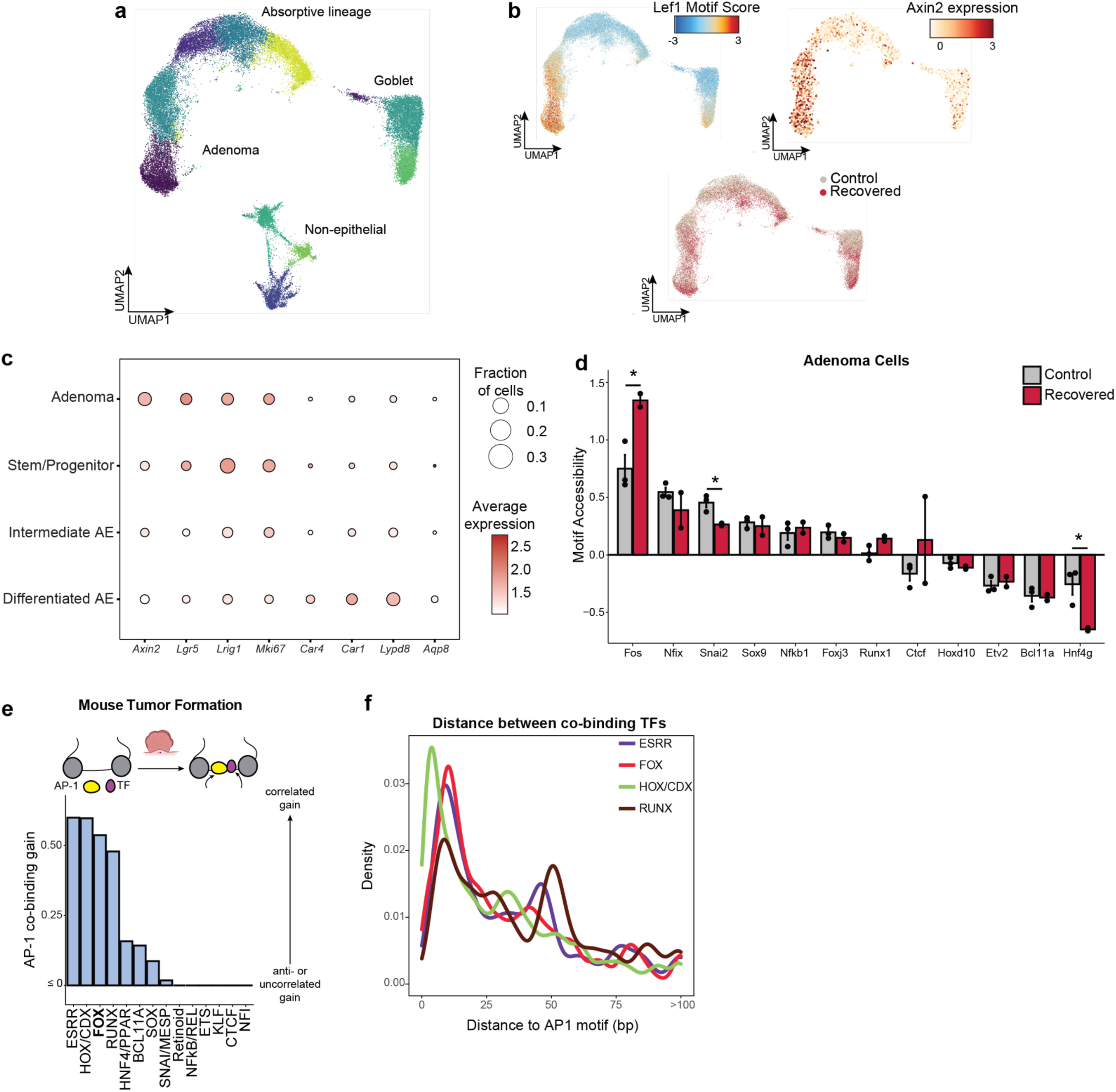
SHARE-seq in colitis recovered adenomas. a, UMAP embedding of scATAC-seq data of primary tissue following adenoma induction with cells colored by cluster. b, UMAP embeddings showing Lef1 motif accessibility (top left), *Axin2* relative gene expression (top right), and sample condition (bottom). c, Marker gene expression for each epithelial cell type identified. For each gene, the size of the circle indicates the fraction of cells in which at least one transcript was detected and the color indicates average expression within the cell type. d, Transcription factor memory following transformation. Each point represents the average motif score in adenoma cells in an animal. e, Co-binding scores with AP-1 factors in primary adenoma cells. f, Distance from each TF family colitis-associated footprint to the nearest AP-1 colitis-associated footprint in the same regulatory element. All error bars are s.e.m. (*) p < 0.05, (**) p < 0.01, (***) p < 0.001

**Extended Data Figure 9.**
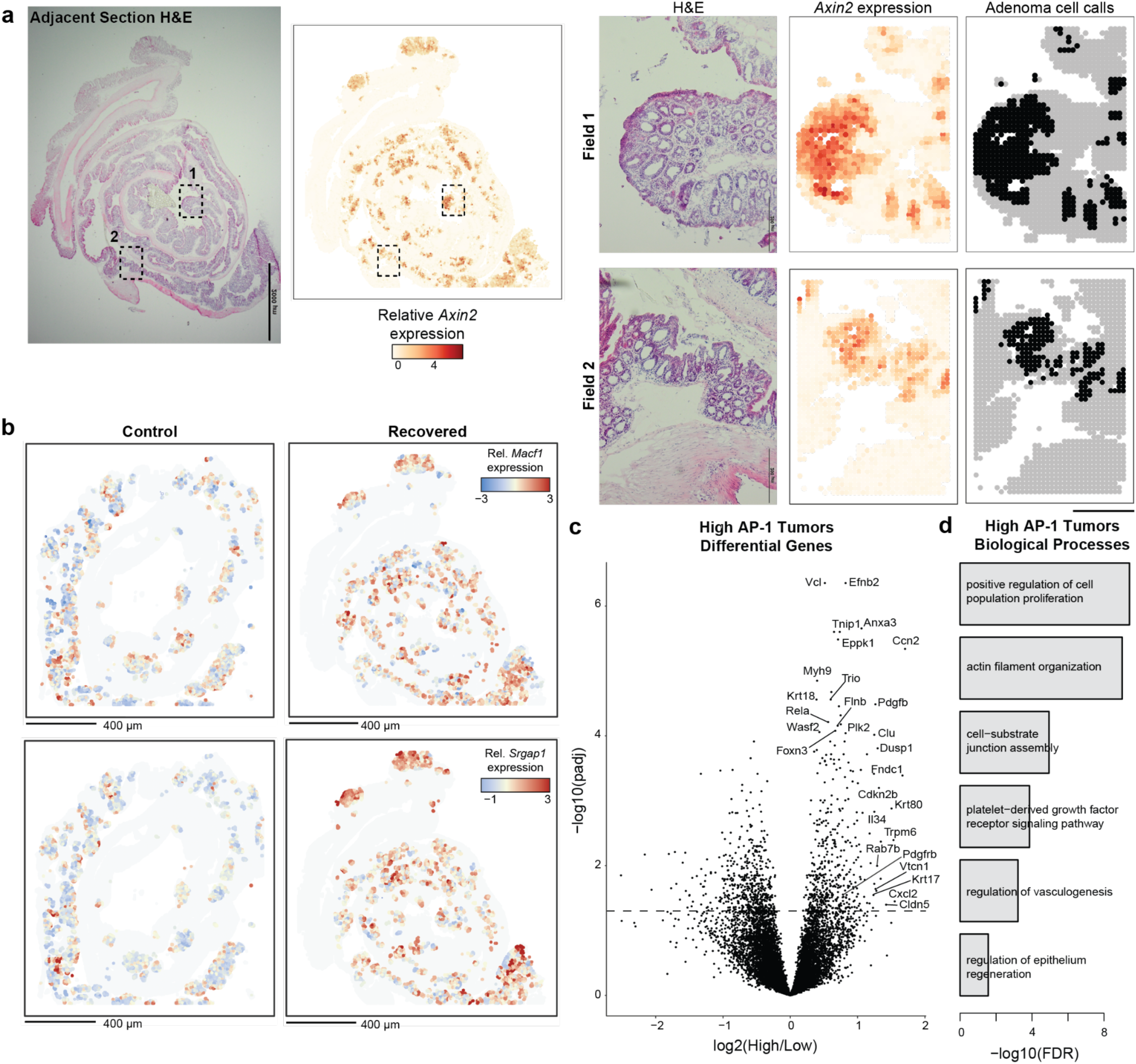
Spatial transcriptomics of adenomas. a, *Axin2* based tumor cell calls. Left, H&E of adjacent section. Scale bar 2 mm. Middle, spatial expression of *Axin2*. Right, boxed regions showing alignment of H&E morphology, *Axin2* expression and individual tumor cell calls. Scale bar 50 μm. b, Spatial expression of AP-1 associated genes in identified tumor cells. Scale bars 400 μm. c, Differential gene expression between tumors with high AP-1 associated gene expression (score > 3) and tumors with low expression. d, Biological processes associated with genes upregulated in high AP-1 tumors.

