## Supplemental Data 1 for "Clonal memory of colitis accumulates and promotes tumor growth"

| pattern | num_seqlets | modisco_cwm_fwd | modisco_cwm_rev | delta_effects | match0 | qval0 | match0_logo | match1 | qval1 | match1_logo | match2 | qval2 | match2_logo |
|---|---|---|---|---|---|---|---|---|---|---|---|---|---|
| de novo #1 | 37 | | | | Spi1 | 1.000000e+00 | | Spic | 1.000000e+00 | | Erf | 1.000000 | |
| de novo #2 | 822 | | | | Dbx1 | 2.321360e-02 | | Pou4f1 | 2.321360e-02 | | Tlx1 | 2.321360e-02 | |
| de novo #3 | 429 | | | | Zfp287 | 1.772060e-04 | | Foxj1 | 7.364640e-03 | | Onecut3 | 1.304850e-02 | |
| de novo #4 | 1048 | | | | Zfp287 | 5.913510e-06 | | Prdm6 | 1.290500e-04 | | Zfp422 | 5.815600e-02 | |
| de novo #5 | 598 | | | | Zfp287 | 9.573920e-08 | | Prdm6 | 1.070780e-06 | | Zfp422 | 1.100520e-04 | |
| de novo #6 | 1448 | | | | Zfp422 | 2.528330e-04 | | Zfp287 | 6.258960e-02 | | Zfp182 | 1.212740e-01 | |
| de novo #7 | 51 | | | | Onecut3 | 1.200830e-01 | | Zfp287 | 1.393100e-01 | | Onecut2 | 0.211331 | |
| de novo #8 | 201 | | | | Stat5b | 1.843740e-01 | | Stat5a | 1.000000e+00 | | Nfatc2 | 1.000000e+00 | |
| de novo #9 | 591 | | | | Foxj1 | 1.060450e-02 | | Zfp287 | 2.122260e-02 | | Ahctf1 | 2.127700e-02 | |
| de novo #10 | 122 | | | | Stat5b | 1.290560e-01 | | Nfatc2 | 1.290560e-01 | | Lcorl | 0.129056 | |
| de novo #11 | 502 | | | | Dbx1 | 9.037290e-03 | | Pou4f2 | 1.022360e-02 | | Tlx1 | 1.022360e-02 | |
| de novo #12 | 3704 | | | | Zfp422 | 8.096640e-02 | | Zfp653 | 9.223390e-01 | | Hmga1 | 9.223390e-01 | |
| de novo #13 | 2227 | | | | Zfp287 | 4.970410e-02 | | Prdm6 | 3.663160e-01 | | Zfp422 | 3.663160e-01 | |
| de novo #14 | 189 | | | | Lcorl | 9.101520e-02 | | Hmg20a | 1.252270e-01 | | Hmg20b | 1.252270e-01 | |
| de novo #15 | 1002 | | | | Zfp422 | 4.450230e-01 | | Prdm6 | 4.450230e-01 | | Mef2c | 5.741100e-01 | |
| de novo #16 | 3152 | | | | Zfp287 | 2.526300e-10 | | Prdm6 | 7.235480e-06 | | Zfp422 | 9.543510e-06 | |

| pattern | num_seqlets | modisco_cwm_fwd | modisco_cwm_rev | delta_effects | match0 | qval0 | match0_logo | match1 | qval1 | match1_logo | match2 | qval2 | match2_logo |
|---|---|---|---|---|---|---|---|---|---|---|---|---|---|
| de novo #17 | 4842 | | | | Zfp422 | 6.847590e-01 | | Hmga1 | 8.756110e-01 | | Hmga1rs1 | 8.756110e-01 | |
| de novo #18 | 1782 | | | | Zfp422 | 6.325630e-03 | | Phf21a | 1.527410e-01 | | Onecut3 | 1.558990e-01 | |
| de novo #19 | 840 | | | | Prdm6 | 1.904400e-04 | | Zfp287 | 3.265320e-04 | | Zfp422 | 3.866320e-03 | |
| de novo #20 | 53 | | | | Foxl1 | 1.673410e-02 | | Foxs1 | 1.673410e-02 | | Foxl2 | 2.029320e-02 | |
| de novo #21 | 31 | | | | Foxl2 | 4.869560e-02 | | Foxl1 | 4.869560e-02 | | Foxs1 | 4.869560e-02 | |
| de novo #22 | 84 | | | | Aire | 7.374560e-01 | | Pdx1 | 7.374560e-01 | | Fosb | 7.374560e-01 | |
| de novo #23 | 654 | | | | Hmg20a | 3.179440e-02 | | Hmg20b | 3.179440e-02 | | Dbx1 | 3.179440e-02 | |
| de novo #24 | 69 | | | | Nfatc2 | 9.180730e-01 | | Nfatc3 | 9.180730e-01 | | Hmga1 | 0.918073 | |
| de novo #25 | 2928 | | | | Zfp287 | 6.254440e-02 | | Dmrtc2 | 3.953860e-01 | | Zfp422 | 3.953860e-01 | |
| de novo #26 | 1811 | | | | Zfp422 | 4.094140e-03 | | Phf21a | 2.559790e-02 | | Hmga1 | 8.141000e-02 | |
| de novo #27 | 499 | | | | Zfp287 | 3.351640e-05 | | Foxj1 | 7.942150e-03 | | Onecut3 | 7.942150e-03 | |
| de novo #28 | 1166 | | | | E2f3 | 7.450860e-02 | | E2f2 | 1.233520e-01 | | Hmga2 | 1.233520e-01 | |
| de novo #29 | 1537 | | | | Zfp422 | 2.691930e-02 | | Hmga1 | 4.812690e-02 | | Hmga1rs1 | 4.812690e-02 | |
| de novo #30 | 43 | | | | Lcorl | 5.553020e-01 | | Lcor | 6.489340e-01 | | Mef2a | 1.000000 | |
| de novo #31 | 1442 | | | | Zfp422 | 2.549470e-02 | | Phf21a | 5.806890e-02 | | Onecut3 | 5.806890e-02 | |

| pattern | num_seqlets | modisco_cwm_fwd | modisco_cwm_rev | delta_effects | match0 | qval0 | match0_logo | match1 | qval1 | match1_logo | match2 | qval2 | match2_logo |
|---|---|---|---|---|---|---|---|---|---|---|---|---|---|
| de novo #32 | 80 | | | | Foxb1 | 3.181580e-01 | | Foxb2 | 3.181580e-01 | | Foxa1 | 3.181580e-01 | |
| de novo #33 | 7011 | | | | Zfp287 | 3.006580e-09 | | Prdm6 | 1.766260e-05 | | Zfp422 | 2.402780e-05 | |
| de novo #34 | 181 | | | | Cdx2 | 1.408320e-05 | | Hoxb9 | 1.511210e-03 | | Hoxd10 | 0.001823 | |
| de novo #35 | 219 | | | | Zfp287 | 1.243550e-05 | | Ahctf1 | 5.099930e-03 | | Onecut3 | 1.725870e-02 | |
| de novo #36 | 2925 | | | | Zfp287 | 6.432160e-10 | | Prdm6 | 8.085120e-06 | | Zfp422 | 8.085120e-06 | |
| de novo #37 | 70 | | | | Foxd3 | 1.228630e-02 | | Foxj3 | 1.228630e-02 | | Foxa2 | 1.228630e-02 | |
| de novo #38 | 1832 | | | | Stat5b | 1.000000e+00 | | Zbtb26 | 1.000000e+00 | | Onecut3 | 1.000000e+00 | |
| de novo #39 | 178 | | | | Zfp287 | 1.047040e-08 | | Prdm6 | 2.810660e-06 | | Zfp422 | 0.000015 | |
| de novo #40 | 84 | | | | Zfp287 | 1.087240e-03 | | Foxj1 | 1.419960e-02 | | Prdm6 | 1.798950e-01 | |
| de novo #41 | 72 | | | | Arid3b | 2.650430e-03 | | Hmga1 | 2.650430e-03 | | Hmga1rs1 | 0.002650 | |
| de novo #42 | 995 | | | | Zfp422 | 8.619030e-05 | | Zfp287 | 3.300600e-03 | | Ahctf1 | 4.891120e-03 | |
| de novo #43 | 5209 | | | | Zfp422 | 1.000000e+00 | | Arid3a | 1.000000e+00 | | Stat5b | 1.000000e+00 | |
| de novo #44 | 5719 | | | | Zfp287 | 2.245740e-09 | | Prdm6 | 1.062610e-05 | | Zfp422 | 1.772310e-05 | |
| de novo #45 | 103 | | | | Prdm6 | 7.685300e-04 | | Zfp287 | 1.149070e-03 | | Hmga1 | 1.415110e-01 | |
| de novo #46 | 1932 | | | | Zfp287 | 2.865800e-09 | | Prdm6 | 1.761310e-06 | | Zfp422 | 1.124950e-04 | |

| pattern | num_seqlets | modisco_cwm_fwd | modisco_cwm_rev | delta_effects | match0 | qval0 | match0_logo | match1 | qval1 | match1_logo | match2 | qval2 | match2_logo |
|---|---|---|---|---|---|---|---|---|---|---|---|---|---|
| de novo #47 | 540 | | | | Zfp287 | 1.013040e-05 | | Ahctf1 | 5.816930e-03 | | Onecut3 | 5.816930e-03 | |
| de novo #48 | 56 | | | | Foxa1 | 5.033160e-04 | | Foxb1 | 5.033160e-04 | | Foxb2 | 0.000503 | |
| de novo #49 | 1153 | | | | Stat5b | 9.863860e-01 | | Hmga1 | 9.863860e-01 | | Hmga1rs1 | 9.863860e-01 | |
| de novo #50 | 1043 | | | | Hmg20a | 3.422490e-03 | | Hmg20b | 3.422490e-03 | | Arid3b | 4.663100e-02 | |
| de novo #51 | 214 | | | | Zfp287 | 1.163460e-04 | | Prdm6 | 1.219970e-03 | | Zfp422 | 2.901820e-03 | |
| de novo #52 | 845 | | | | Dbx1 | 1.599150e-02 | | Tlx1 | 2.179860e-02 | | Tlx2 | 2.179860e-02 | |
| de novo #53 | 492 | | | | Zfp422 | 1.118690e-02 | | Zfp287 | 4.042720e-02 | | Hmga1 | 9.814870e-02 | |
| de novo #54 | 149 | | | | Zfp287 | 8.284720e-04 | | Zfp879 | 2.646410e-02 | | Onecut3 | 0.031811 | |
| de novo #55 | 4030 | | | | Zfp422 | 9.031700e-02 | | Hmga1 | 8.601250e-01 | | Hmga1rs1 | 8.601250e-01 | |
| de novo #56 | 113 | | | | Hoxb9 | 3.496850e-02 | | Hoxd12 | 3.496850e-02 | | Cdx1 | 3.496850e-02 | |
| de novo #57 | 543 | | | | Zfp287 | 1.432580e-06 | | Zfp422 | 2.500160e-05 | | Prdm6 | 4.701770e-05 | |
| de novo #58 | 602 | | | | Zfp422 | 1.643460e-05 | | Zfp287 | 1.070010e-03 | | Prdm6 | 1.522820e-03 | |
| de novo #59 | 138 | | | | Lcorl | 2.937400e-02 | | Lcor | 1.070470e-01 | | Stat5b | 5.853570e-01 | |
| de novo #60 | 1677 | | | | Zfp287 | 4.305550e-06 | | Zfp422 | 8.882290e-06 | | Prdm6 | 7.783480e-05 | |
| de novo #61 | 102 | | | | Cdx2 | 6.084670e-04 | | Hoxd10 | 9.032200e-04 | | Hoxb9 | 2.316980e-03 | |

| pattern | num_seqlets | modisco_cwm_fwd | modisco_cwm_rev | delta_effects | match0 | qval0 | match0_logo | match1 | qval1 | match1_logo | match2 | qval2 | match2_logo |
|---|---|---|---|---|---|---|---|---|---|---|---|---|---|
| de novo #62 | 2447 | | | | Zfp287 | 9.068750e-09 | | Prdm6 | 4.181820e-06 | | Zfp422 | 9.032410e-06 | |
| de novo #63 | 1307 | | | | Zfp287 | 1.202760e-06 | | Prdm6 | 7.505940e-03 | | Zfp182 | 1.222340e-02 | |
| de novo #64 | 733 | | | | Zfp422 | 1.755540e-02 | | Prdm6 | 1.755540e-02 | | Zfp287 | 1.755540e-02 | |
| de novo #65 | 82 | | | | Foxj3 | 9.184690e-03 | | Foxc2 | 9.184690e-03 | | Foxa1 | 0.009185 | |
| de novo #66 | 6447 | | | | Arid3a | 9.083160e-01 | | Hmga1 | 9.083160e-01 | | Hmga1rs1 | 9.083160e-01 | |
| de novo #67 | 859 | | | | Zfp422 | 2.730660e-05 | | Zfp287 | 2.117900e-04 | | Prdm6 | 1.114120e-02 | |
| de novo #68 | 2727 | | | | Zfp287 | 3.946280e-01 | | Zfp422 | 3.946280e-01 | | Bcl6b | 8.281590e-01 | |
| de novo #69 | 3023 | | | | Hmga1 | 5.997750e-01 | | Hmga1rs1 | 5.997750e-01 | | Arid3b | 5.997750e-01 | |
| de novo #70 | 267 | | | | Hmg20a | 1.896330e-02 | | Hmg20b | 1.896330e-02 | | Phf21a | 1.896330e-02 | |
| de novo #71 | 121 | | | | Hmga1 | 9.657970e-04 | | Hmga1rs1 | 9.657970e-04 | | Tbp | 2.017060e-03 | |
| de novo #72 | 546 | | | | Zfp287 | 3.162490e-04 | | Prdm6 | 6.699180e-03 | | Zfp422 | 1.053430e-02 | |
| de novo #73 | 112 | | | | Phf21a | 4.855030e-01 | | Hmga1 | 4.855030e-01 | | Hmga1rs1 | 4.855030e-01 | |
| de novo #74 | 456 | | | | Hmga1 | 5.958860e-01 | | Hmga1rs1 | 5.958860e-01 | | Phf21a | 0.595886 | |
| de novo #75 | 4534 | | | | Zfp287 | 1.291410e-09 | | Prdm6 | 6.742950e-06 | | Zfp422 | 1.313530e-05 | |
| de novo #76 | 547 | | | | Zfp287 | 1.720780e-07 | | Prdm6 | 2.203830e-06 | | Zfp422 | 0.000016 | |

| pattern | num_seqlets | modisco_cwm_fwd | modisco_cwm_rev | delta_effects | match0 | qval0 | match0_logo | match1 | qval1 | match1_logo | match2 | qval2 | match2_logo |
|---|---|---|---|---|---|---|---|---|---|---|---|---|---|
| de novo #77 | 74 | | | | Zfp879 | 2.667670e-01 | | Phf21a | 2.667670e-01 | | Lcorl | 0.266767 | |
| de novo #78 | 33 | | | | Lcorl | 1.635620e-01 | | Lcor | 4.738640e-01 | | Mef2a | 0.811895 | |
| de novo #79 | 437 | | | | Zfp422 | 8.533530e-05 | | Zfp287 | 5.374730e-04 | | Ahctf1 | 1.078770e-01 | |
| de novo #80 | 1869 | | | | Zfp287 | 5.241820e-02 | | Prdm6 | 7.345790e-02 | | Zfp422 | 2.182690e-01 | |
| de novo #81 | 269 | | | | Prdm6 | 9.192990e-02 | | Kdm2b | 9.192990e-02 | | Hmg20a | 1.290550e-01 | |
| de novo #82 | 14083 | | | | Fosl2 | 2.123640e-03 | | Fos | 2.123640e-03 | | Bach2 | 2.794930e-03 | |
| de novo #83 | 75 | | | | Fosb | 2.758940e-01 | | Jund | 2.758940e-01 | | Mga | 2.758940e-01 | |
| de novo #84 | 53 | | | | Fosb | 1.000000e+00 | | Mafa | 1.000000e+00 | | Mafk | 1.000000e+00 | |
| de novo #85 | 11965 | | | | Fosl2 | 3.390240e-04 | | Fos | 3.381150e-03 | | Fosb | 0.003381 | |
| de novo #86 | 7678 | | | | Bach2 | 2.271530e-03 | | Fosl2 | 2.271530e-03 | | Fos | 2.271530e-03 | |
| de novo #87 | 28 | | | | Ppard | 4.615580e-03 | | Nr2c2 | 4.615580e-03 | | Fosb | 4.615580e-03 | |
| de novo #88 | 210 | | | | Fosb | 1.093940e-03 | | Fos | 6.420790e-03 | | Jund | 0.007821 | |
| de novo #89 | 6318 | | | | Bach2 | 2.255230e-03 | | Fosl2 | 2.255230e-03 | | Fos | 2.255230e-03 | |
| de novo #90 | 9638 | | | | Fosl2 | 3.152090e-03 | | Fos | 3.152090e-03 | | Bach2 | 3.614410e-03 | |
| de novo #91 | 46 | | | | Mga | 2.358490e-01 | | Tbx4 | 2.358490e-01 | | Tbx5 | 0.235849 | |

| pattern | num_seqlets | modisco_cwm_fwd | modisco_cwm_rev | delta_effects | match0 | qval0 | match0_logo | match1 | qval1 | match1_logo | match2 | qval2 | match2_logo |
|---|---|---|---|---|---|---|---|---|---|---|---|---|---|
| de novo #92 | 7638 | | | | Bach2 | 2.659940e-03 | | Fosl2 | 2.659940e-03 | | Fosb | 2.659940e-03 | |
| de novo #93 | 56 | | | | Fosb | 1.000000e+00 | | Mafa | 1.000000e+00 | | Jund | 1.000000e+00 | |
| de novo #94 | 8313 | | | | Fosl2 | 3.216590e-03 | | Fos | 3.216590e-03 | | Bach2 | 3.226330e-03 | |
| de novo #95 | 63 | | | | Arntl | 1.144040e-02 | | Tfe3 | 1.144040e-02 | | Tfeb | 1.144040e-02 | |
| de novo #96 | 10258 | | | | Fosl2 | 1.761910e-03 | | Fos | 1.761910e-03 | | Bach2 | 2.495220e-03 | |
| de novo #97 | 49 | | | | Fosb | 9.999980e-01 | | Mafa | 9.999980e-01 | | Tbx15 | 9.999980e-01 | |
| de novo #98 | 108 | | | | Fosb | 3.471140e-01 | | Jund | 3.940460e-01 | | Junb | 0.745869 | |
| de novo #99 | 22 | | | | Fosb | 1.290610e-04 | | Jund | 6.236450e-02 | | Fosl2 | 7.204690e-02 | |
| de novo #100 | 100 | | | | Hoxd11 | 8.195540e-02 | | Jund | 8.195540e-02 | | Hoxc10 | 8.195540e-02 | |
| de novo #101 | 26 | | | | Nr4a2 | 7.795930e-03 | | Nr4a3 | 7.795930e-03 | | Junb | 1.529980e-01 | |
| de novo #102 | 7800 | | | | Bach2 | 2.478920e-03 | | Fosl2 | 2.478920e-03 | | Fos | 2.478920e-03 | |
| de novo #103 | 71 | | | | Hoxd10 | 4.665250e-02 | | Hoxd11 | 4.665250e-02 | | Jund | 4.665250e-02 | |
| de novo #104 | 23 | | | | Nr4a2 | 1.075660e-02 | | Nr4a3 | 1.075660e-02 | | Junb | 1.846900e-01 | |
| de novo #105 | 64 | | | | Jund | 9.707080e-02 | | Hoxd11 | 9.707080e-02 | | Hoxd10 | 9.707080e-02 | |
| de novo #106 | 122 | | | | Hoxd11 | 9.414860e-02 | | Hoxd10 | 9.414860e-02 | | Hoxc10 | 9.414860e-02 | |

| pattern | num_seqlets | modisco_cwm_fwd | modisco_cwm_rev | delta_effects | match0 | qval0 | match0_logo | match1 | qval1 | match1_logo | match2 | qval2 | match2_logo |
|---|---|---|---|---|---|---|---|---|---|---|---|---|---|
| de novo #107 | 11581 | | | | Fosl2 | 2.181910e-03 | | Fos | 2.181910e-03 | | Bach2 | 0.002871 | |
| de novo #108 | 52 | | | | Mga | 2.438900e-01 | | Tbx4 | 2.438900e-01 | | Tbx5 | 2.438900e-01 | |
| de novo #109 | 23 | | | | Nr4a2 | 7.703990e-04 | | Nr4a3 | 7.703990e-04 | | Rarg | 0.271699 | |
| de novo #110 | 32 | | | | Mafa | 6.076560e-03 | | Fosb | 1.338000e-02 | | Bach2 | 4.036000e-02 | |
| de novo #111 | 12835 | | | | Fosl2 | 2.152640e-03 | | Fos | 2.152640e-03 | | Bach2 | 2.821790e-03 | |
| de novo #112 | 500 | | | | Batf3 | 1.410160e-02 | | Foxb1 | 1.520540e-02 | | Foxb2 | 1.520540e-02 | |
| de novo #113 | 6974 | | | | Fosl2 | 2.934750e-03 | | Fos | 2.934750e-03 | | Bach2 | 3.051290e-03 | |
| de novo #114 | 52 | | | | Nr1h3 | 3.141360e-03 | | Nr2c2 | 9.300220e-03 | | Pparg | 9.300220e-03 | |
| de novo #115 | 38 | | | | Hnf4g | 5.211040e-03 | | Nr1h3 | 9.307750e-03 | | Hnf4a | 3.346690e-02 | |
| de novo #116 | 12534 | | | | Hnf4g | 8.803960e-09 | | Hnf4a | 1.308760e-05 | | Nr1h3 | 2.267470e-05 | |
| de novo #117 | 39 | | | | Nr1h3 | 1.720710e-03 | | Nr2c2 | 1.720710e-03 | | Pparg | 0.004057 | |
| de novo #118 | 182 | | | | Hnf4g | 1.587790e-08 | | Nr1h3 | 2.561520e-05 | | Hnf4a | 0.000255 | |
| de novo #119 | 83 | | | | Hnf4g | 3.238640e-03 | | Rarg | 9.019830e-03 | | Hnf4a | 8.759430e-02 | |
| de novo #120 | 27 | | | | Nr2c2 | 7.812650e-03 | | Ppard | 7.812650e-03 | | Hnf4g | 7.812650e-03 | |
| de novo #121 | 11793 | | | | Hnf4g | 1.486930e-07 | | Nr1h3 | 1.063210e-05 | | Hnf4a | 1.063210e-05 | |

| pattern | num_seqlets | modisco_cwm_fwd | modisco_cwm_rev | delta_effects | match0 | qval0 | match0_logo | match1 | qval1 | match1_logo | match2 | qval2 | match2_logo |
|---|---|---|---|---|---|---|---|---|---|---|---|---|---|
| | | 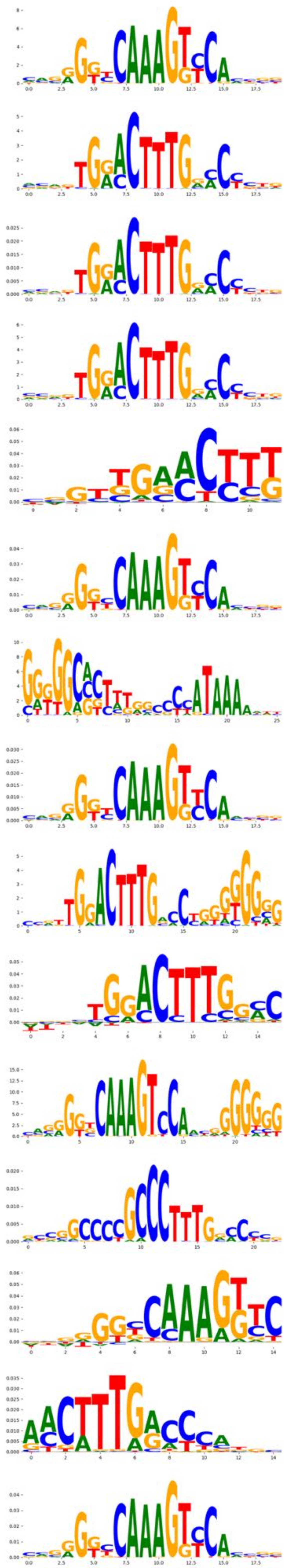 | 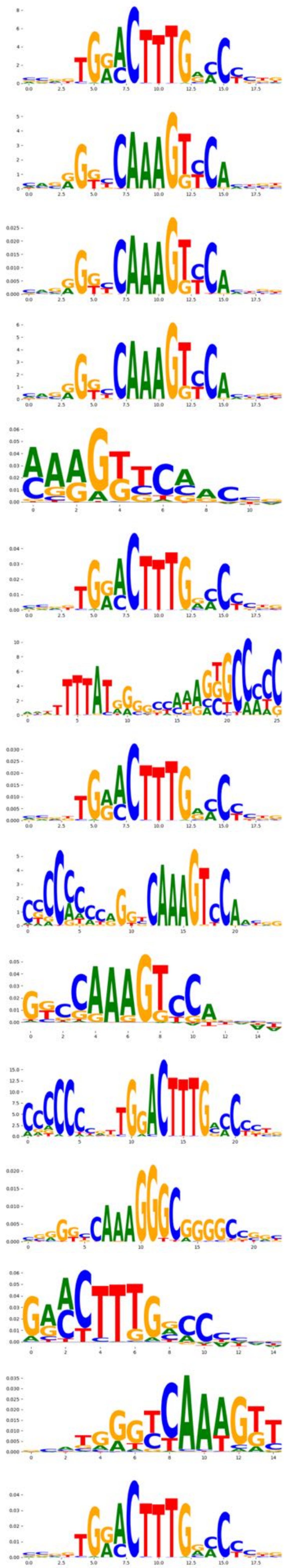 | 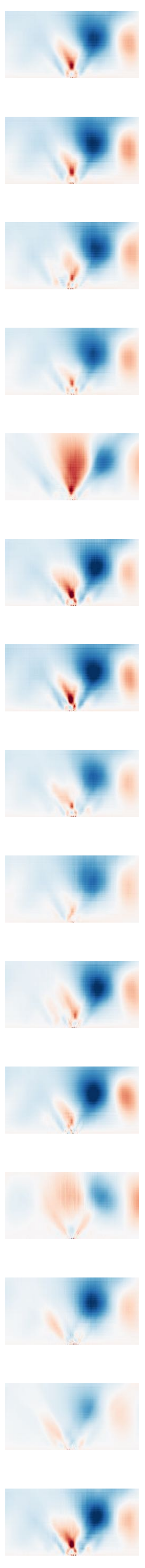 | | | 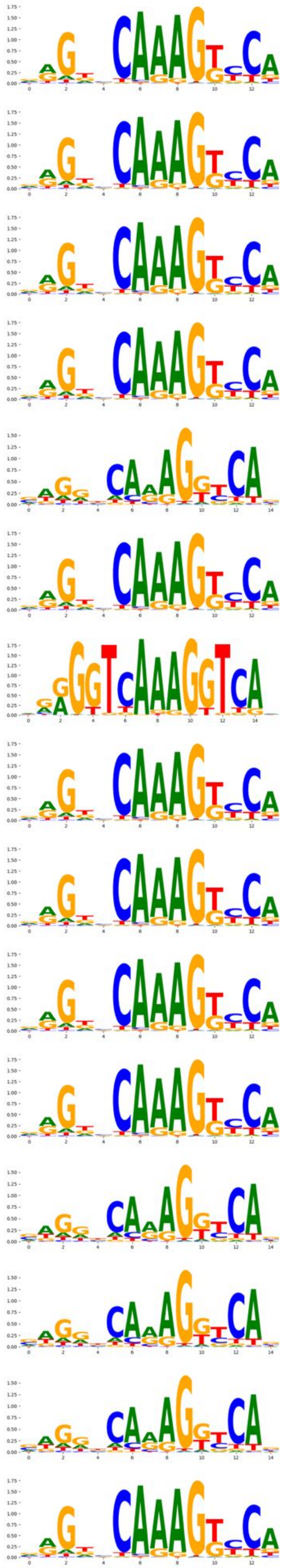 | | | 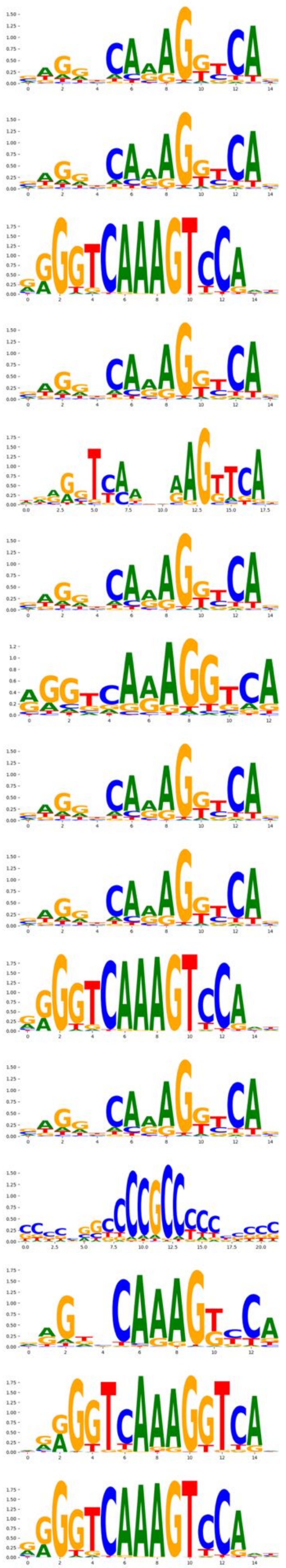 | | | 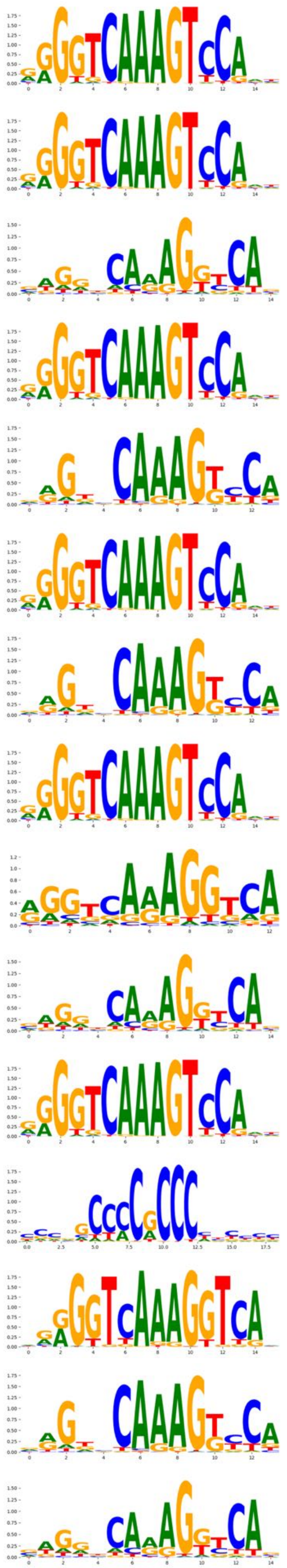 |
| de novo #122 | 7003 | | | | Hnf4g | 2.734790e-08 | | Nr1h3 | 3.019100e-05 | | Hnf4a | 3.128750e-05 | |
| de novo #123 | 18200 | | | | Hnf4g | 1.130930e-06 | | Hnf4a | 1.626550e-05 | | Nr1h3 | 0.000022 | |
| de novo #124 | 7794 | | | | Hnf4g | 3.365370e-08 | | Nr1h3 | 2.324910e-05 | | Hnf4a | 2.324910e-05 | |
| de novo #125 | 59 | | | | Nr1h3 | 1.151940e-03 | | Nr1i2 | 1.151940e-03 | | Hnf4g | 0.007071 | |
| de novo #126 | 24734 | | | | Hnf4g | 7.803890e-07 | | Nr1h3 | 8.006090e-06 | | Hnf4a | 0.000020 | |
| de novo #127 | 61 | | | | Ppard | 4.537550e-04 | | Nr2c2 | 4.537550e-04 | | Hnf4g | 2.768100e-03 | |
| de novo #128 | 20574 | | | | Hnf4g | 1.129920e-06 | | Nr1h3 | 1.801230e-05 | | Hnf4a | 2.456620e-05 | |
| de novo #129 | 30 | | | | Hnf4g | 4.825680e-04 | | Nr1h3 | 3.379230e-03 | | Nr2c2 | 8.362710e-03 | |
| de novo #130 | 53 | | | | Hnf4g | 8.050360e-05 | | Hnf4a | 5.189290e-03 | | Nr1h3 | 6.364160e-03 | |
| de novo #131 | 43 | | | | Hnf4g | 1.657000e-03 | | Nr1h3 | 3.191940e-03 | | Hnf4a | 3.513060e-03 | |
| de novo #132 | 48 | | | | Nr1h3 | 3.355830e-02 | | Sp2 | 3.355830e-02 | | Klf3 | 5.153700e-02 | |
| de novo #133 | 56 | | | | Nr1h3 | 1.610720e-05 | | Hnf4g | 5.836350e-04 | | Ppard | 7.898150e-04 | |
| de novo #134 | 99 | | | | Nr1h3 | 1.303280e-03 | | Ppard | 1.303280e-03 | | Hnf4g | 1.303280e-03 | |
| de novo #135 | 21835 | | | | Hnf4g | 3.216390e-07 | | Hnf4a | 1.410670e-05 | | Nr1h3 | 1.410670e-05 | |
| de novo #136 | 111 | | | | Hnf4a | 4.482770e-06 | | Esrrb | 5.794250e-02 | | Hnf4g | 5.794250e-02 | |

| pattern | num_seqlets | modisco_cwm_fwd | modisco_cwm_rev | delta_effects | match0 | qval0 | match0_logo | match1 | qval1 | match1_logo | match2 | qval2 | match2_logo |
|---|---|---|---|---|---|---|---|---|---|---|---|---|---|
| de novo #137 | 12297 | | | | Hnf4a | 5.667580e-06 | | Hnf4g | 5.667580e-06 | | Nr1h3 | 1.251150e-03 | |
| de novo #138 | 46 | | | | Hnf4g | 1.182750e-04 | | Ppard | 1.244520e-04 | | Nr1h3 | 0.000553 | |
| de novo #139 | 86 | | | | Hnf4a | 4.470080e-06 | | Esrrb | 5.697020e-02 | | Esrra | 5.697020e-02 | |
| de novo #140 | 23241 | | | | Hnf4g | 2.367900e-06 | | Nr1h3 | 6.159820e-06 | | Hnf4a | 2.403120e-05 | |
| de novo #141 | 14406 | | | | Hnf4g | 3.445030e-08 | | Hnf4a | 1.550170e-05 | | Nr1h3 | 1.689000e-05 | |
| de novo #142 | 9412 | | | | Hnf4g | 2.774410e-08 | | Nr1h3 | 1.770000e-05 | | Hnf4a | 1.770000e-05 | |
| de novo #143 | 103 | | | | Hnf4g | 2.330990e-04 | | Pparg | 5.569710e-03 | | Nr2c2 | 5.569710e-03 | |
| de novo #144 | 196 | | | | Nr5a1 | 2.828560e-03 | | Nr2c2 | 2.828560e-03 | | Esrra | 2.828560e-03 | |
| de novo #145 | 7289 | | | | Ppard | 5.247990e-08 | | Nr2c2 | 3.908270e-06 | | Ppara | 3.583600e-05 | |
| de novo #146 | 41 | | | | Ppard | 6.378380e-02 | | Nr2c2 | 9.832770e-02 | | Pparg | 0.098328 | |
| de novo #147 | 46 | | | | Mbtps2 | 4.365700e-02 | | Yy1 | 4.365700e-02 | | Yy2 | 4.365700e-02 | |
| de novo #148 | 20020 | | | | Ctcf | 3.987550e-18 | | Ctcfl | 9.961500e-07 | | Zfp661 | 9.839710e-02 | |
| de novo #149 | 19926 | | | | Ctcf | 5.579200e-18 | | Ctcfl | 8.469540e-07 | | Zfp661 | 1.004840e-01 | |
| de novo #150 | 138 | | | | Ctcf | 4.669660e-04 | | Ctcfl | 1.091950e-02 | | Tcf12 | 3.923700e-02 | |
| de novo #151 | 22787 | | | | Ctcf | 2.172540e-18 | | Ctcfl | 7.500890e-07 | | Zfp661 | 9.690380e-02 | |

| pattern | num_seqlets | modisco_cwm_fwd | modisco_cwm_rev | delta_effects | match0 | qval0 | match0_logo | match1 | qval1 | match1_logo | match2 | qval2 | match2_logo |
|---|---|---|---|---|---|---|---|---|---|---|---|---|---|
| de novo #152 | 507 | | | | Ctcfl | 9.343630e-06 | | Ctcf | 2.663360e-05 | | Zfp46 | 2.595330e-01 | |
| de novo #153 | 23263 | | | | Ctcf | 3.508700e-18 | | Ctcfl | 6.175850e-07 | | Zfp661 | 7.864470e-02 | |
| de novo #154 | 21200 | | | | Ctcf | 3.549170e-18 | | Ctcfl | 9.396300e-07 | | Zfp661 | 8.634770e-02 | |
| de novo #155 | 22073 | | | | Ctcf | 9.767180e-19 | | Ctcfl | 1.053800e-06 | | Zfp661 | 9.368690e-02 | |
| de novo #156 | 21249 | | | | Ctcf | 7.640640e-19 | | Ctcfl | 5.911040e-07 | | Zfp661 | 1.022060e-01 | |
| de novo #157 | 179 | | | | Ctcf | 8.098070e-03 | | Ctcfl | 2.508240e-02 | | Zfp423 | 1.000000e+00 | |
| de novo #158 | 22155 | | | | Ctcf | 2.045610e-18 | | Ctcfl | 9.443880e-07 | | Zfp661 | 8.792090e-02 | |
| de novo #159 | 555 | | | | Ctcfl | 3.827220e-06 | | Ctcf | 3.595430e-05 | | Zfp46 | 2.241990e-01 | |
| de novo #160 | 721 | | | | Ctcf | 3.066420e-05 | | Ctcfl | 5.411670e-05 | | Zfp46 | 1.207990e-01 | |
| de novo #161 | 20735 | | | | Ctcf | 1.227970e-17 | | Ctcfl | 5.000180e-07 | | Zfp661 | 0.092693 | |
| de novo #162 | 1079 | | | | Ctcfl | 1.064980e-06 | | Ctcf | 2.694810e-06 | | Zfp661 | 1.507180e-01 | |
| de novo #163 | 21440 | | | | Ctcf | 2.227560e-17 | | Ctcfl | 6.899160e-07 | | Zfp661 | 0.106021 | |
| de novo #164 | 41 | | | | Ctcf | 1.035510e-02 | | Mbtps2 | 2.563730e-02 | | Yy1 | 2.563730e-02 | |
| de novo #165 | 122 | | | | Ctcfl | 1.546560e-03 | | Ctcf | 1.347320e-02 | | Klf11 | 0.017045 | |
| de novo #166 | 80 | | | | Ctcf | 8.216710e-03 | | Mbtps2 | 1.130980e-02 | | Yy1 | 1.130980e-02 | |

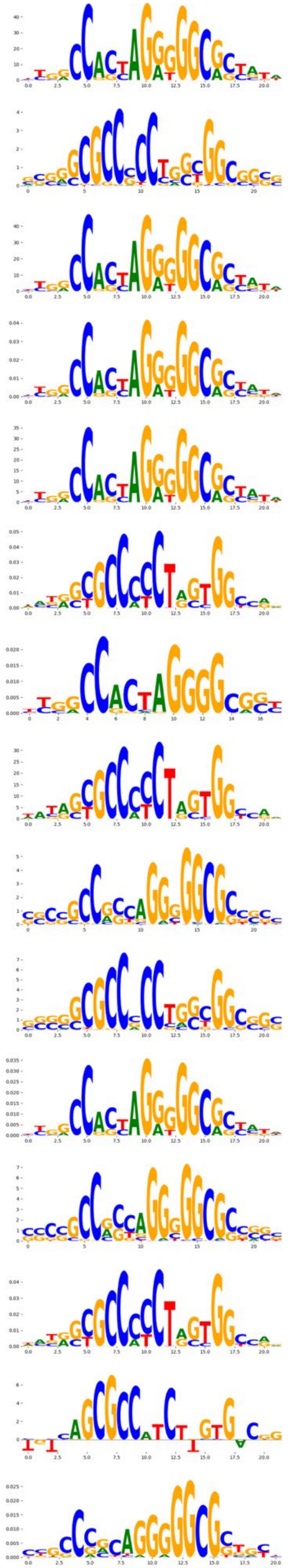
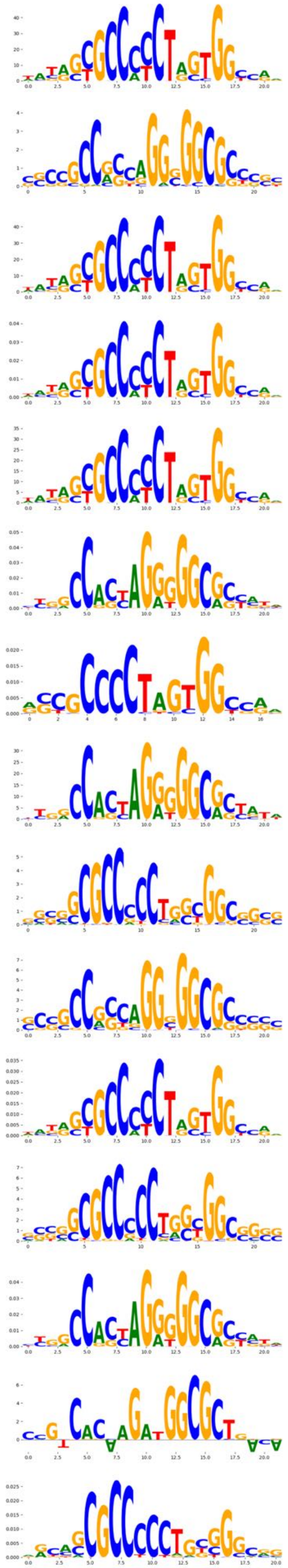
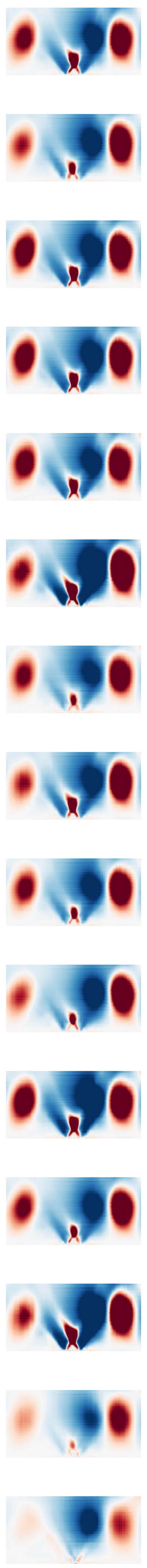
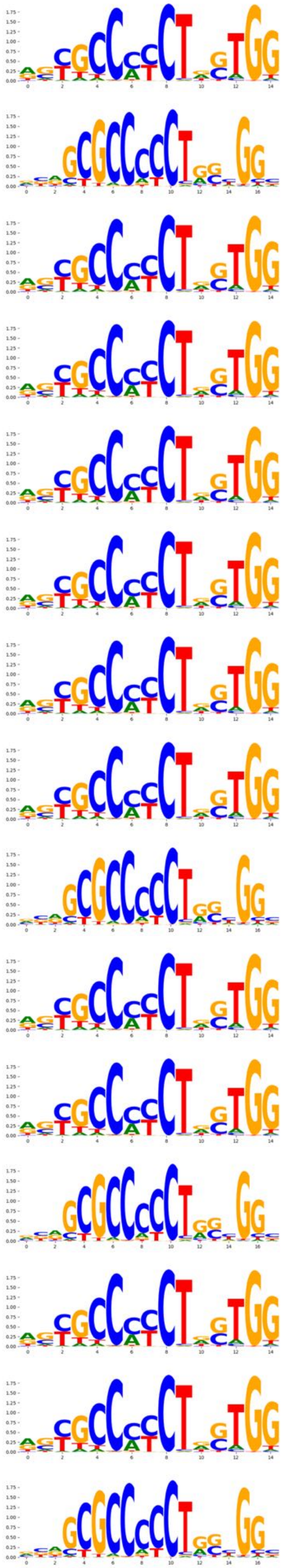
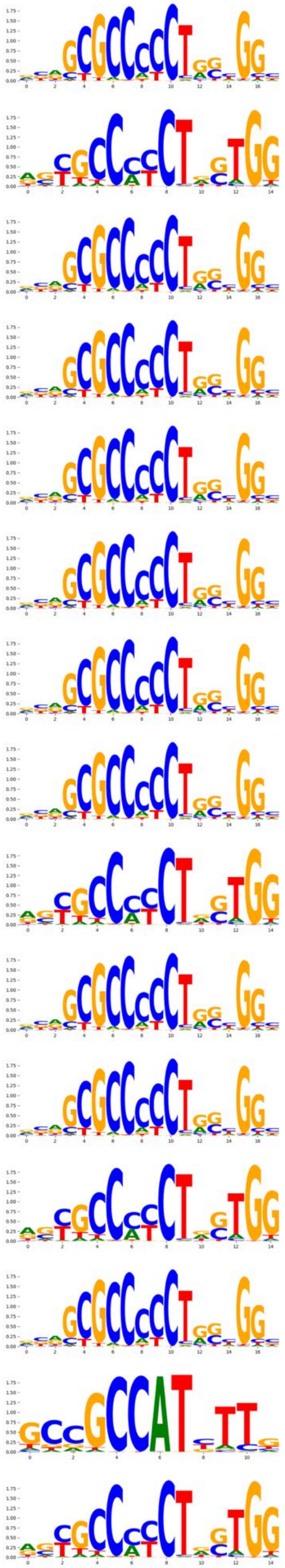
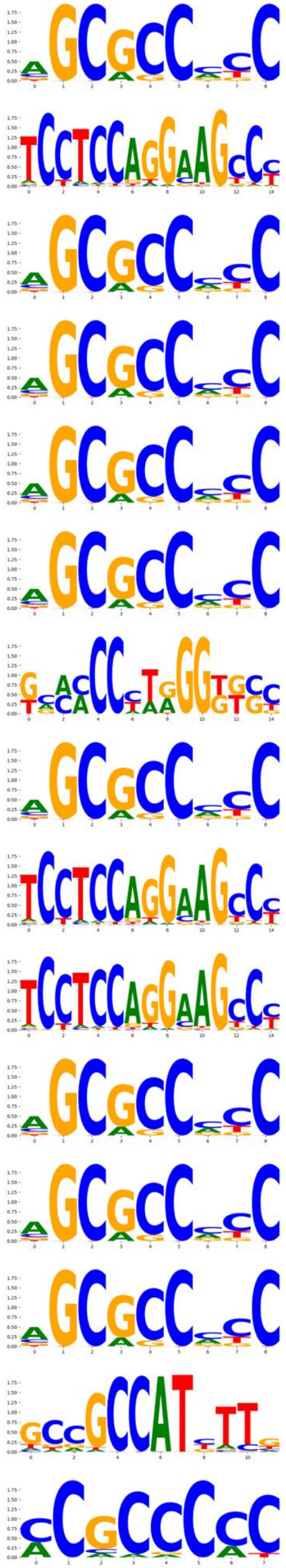

| pattern | num_seqlets | modisco_cwm_fwd | modisco_cwm_rev | delta_effects | match0 | qval0 | match0_logo | match1 | qval1 | match1_logo | match2 | qval2 | match2_logo |
|---|---|---|---|---|---|---|---|---|---|---|---|---|---|
| de novo #167 | 22122 | | | | Ctcf | 4.517090e-18 | | Ctcfl | 6.391610e-07 | | Zfp661 | 8.821040e-02 | |
| de novo #168 | 24 | | | | Ctcf | 1.541540e-01 | | Neurod2 | 2.061180e-01 | | Rest | 0.346957 | |
| de novo #169 | 23303 | | | | Ctcf | 3.663750e-18 | | Ctcfl | 5.266890e-07 | | Zfp661 | 9.921580e-02 | |
| de novo #170 | 4772 | | | | Foxi1 | 1.172530e-01 | | Nfyb | 1.172530e-01 | | Nfya | 1.172530e-01 | |
| de novo #171 | 5682 | | | | Foxi1 | 1.289550e-01 | | Nfyb | 1.289550e-01 | | Nfyc | 1.289550e-01 | |
| de novo #172 | 5606 | | | | Foxi1 | 1.270340e-01 | | Nfyb | 1.270340e-01 | | Nfya | 1.270340e-01 | |
| de novo #173 | 5697 | | | | Foxi1 | 1.115870e-01 | | Nfyb | 1.115870e-01 | | Nfya | 0.111587 | |
| de novo #174 | 4978 | | | | Foxi1 | 1.268460e-01 | | Nfyb | 1.268460e-01 | | Nfyc | 1.268460e-01 | |
| de novo #175 | 5267 | | | | Foxi1 | 1.290340e-01 | | Nfyb | 1.290340e-01 | | Nfyc | 0.129034 | |
| de novo #176 | 5021 | | | | Foxi1 | 1.004580e-01 | | Nfyb | 1.004580e-01 | | Nfyc | 1.004580e-01 | |
| de novo #177 | 4803 | | | | Foxi1 | 1.268220e-01 | | Nfyb | 1.268220e-01 | | Nfyc | 1.268220e-01 | |
| de novo #178 | 4863 | | | | Foxi1 | 1.134790e-01 | | Nfyb | 1.134790e-01 | | Nfyc | 1.134790e-01 | |
| de novo #179 | 5453 | | | | Foxi1 | 1.275380e-01 | | Nfyb | 1.275380e-01 | | Nfyc | 1.275380e-01 | |
| de novo #180 | 4833 | | | | Foxi1 | 5.700760e-02 | | Nfyb | 5.700760e-02 | | Nfyc | 5.700760e-02 | |
| de novo #181 | 4808 | | | | Foxi1 | 1.266910e-01 | | Nfyb | 1.266910e-01 | | Nfyc | 1.266910e-01 | |

| pattern | num_seqlets | modisco_cwm_fwd | modisco_cwm_rev | delta_effects | match0 | qval0 | match0_logo | match1 | qval1 | match1_logo | match2 | qval2 | match2_logo |
|---|---|---|---|---|---|---|---|---|---|---|---|---|---|
| de novo #182 | 59 | | | | Foxl2 | 5.489810e-03 | | Foxa1 | 1.098120e-02 | | Foxc1 | 1.098120e-02 | |
| de novo #183 | 32 | | | | Foxa1 | 1.418390e-03 | | Foxc1 | 6.872920e-03 | | Foxd3 | 0.006873 | |
| de novo #184 | 47 | | | | Foxa2 | 4.273560e-04 | | Foxa3 | 5.917090e-03 | | Foxa1 | 2.244270e-02 | |
| de novo #185 | 151 | | | | Foxa1 | 3.325200e-03 | | Foxb1 | 3.325200e-03 | | Foxb2 | 3.325200e-03 | |
| de novo #186 | 227 | | | | Foxb1 | 4.284640e-02 | | Foxb2 | 4.284640e-02 | | Foxa1 | 0.042846 | |
| de novo #187 | 55 | | | | Foxa1 | 7.942870e-05 | | Foxb1 | 6.630460e-04 | | Foxb2 | 6.630460e-04 | |
| de novo #188 | 60 | | | | Foxa1 | 4.094640e-01 | | Pou4f2 | 4.094640e-01 | | Foxb1 | 4.094640e-01 | |
| de novo #189 | 609 | | | | Foxb1 | 1.197860e-01 | | Foxb2 | 1.197860e-01 | | Foxa1 | 1.197860e-01 | |
| de novo #190 | 318 | | | | Batf3 | 2.248070e-02 | | Foxb1 | 2.248070e-02 | | Foxb2 | 2.248070e-02 | |
| de novo #191 | 1200 | | | | Foxa1 | 3.555390e-04 | | Foxb1 | 3.555390e-04 | | Foxb2 | 3.555390e-04 | |
| de novo #192 | 12015 | | | | Foxa1 | 2.896070e-05 | | Foxa3 | 2.896070e-05 | | Foxd1 | 4.545350e-05 | |
| de novo #193 | 980 | | | | Foxa1 | 4.254500e-06 | | Foxb1 | 1.284130e-05 | | Foxb2 | 1.284130e-05 | |
| de novo #194 | 716 | | | | Foxa1 | 8.324730e-04 | | Foxa2 | 1.310010e-03 | | Foxc1 | 1.310010e-03 | |
| de novo #195 | 568 | | | | Foxa1 | 4.283670e-02 | | Foxb1 | 4.283670e-02 | | Foxb2 | 4.283670e-02 | |
| de novo #196 | 198 | | | | Foxe1 | 2.070390e-03 | | Foxe3 | 2.070390e-03 | | Foxj1 | 0.007988 | |

| pattern | num_seqlets | modisco_cwm_fwd | modisco_cwm_rev | delta_effects | match0 | qval0 | match0_logo | match1 | qval1 | match1_logo | match2 | qval2 | match2_logo |
|---|---|---|---|---|---|---|---|---|---|---|---|---|---|
| de novo #197 | 89 | | | | Foxa1 | 1.875180e-03 | | Foxd3 | 2.039810e-03 | | Foxa3 | 4.191990e-03 | |
| de novo #198 | 2287 | | | | Foxa1 | 4.268350e-07 | | Foxb1 | 1.093420e-05 | | Foxb2 | 1.093420e-05 | |
| de novo #199 | 2365 | | | | Foxa1 | 1.297570e-06 | | Foxb1 | 1.779460e-05 | | Foxb2 | 1.779460e-05 | |
| de novo #200 | 5096 | | | | Foxa1 | 7.145400e-07 | | Foxa2 | 3.871600e-06 | | Foxa3 | 4.078610e-06 | |
| de novo #201 | 283 | | | | Foxb1 | 3.966450e-02 | | Foxb2 | 3.966450e-02 | | Foxa1 | 3.966450e-02 | |
| de novo #202 | 67 | | | | Foxe1 | 2.983470e-03 | | Foxe3 | 2.983470e-03 | | Foxa3 | 9.905380e-03 | |
| de novo #203 | 12389 | | | | Foxa1 | 5.018210e-06 | | Foxa3 | 5.018210e-06 | | Foxd1 | 0.000005 | |
| de novo #204 | 8789 | | | | Foxa1 | 1.157550e-06 | | Foxa3 | 2.315100e-06 | | Foxd1 | 2.700950e-06 | |
| de novo #205 | 357 | | | | Foxe1 | 1.516140e-03 | | Foxe3 | 1.516140e-03 | | Foxj1 | 1.516140e-03 | |
| de novo #206 | 5994 | | | | Foxa1 | 4.301150e-06 | | Foxa3 | 8.602300e-06 | | Foxd1 | 0.000010 | |
| de novo #207 | 179 | | | | Foxa1 | 1.130600e-03 | | Foxb1 | 1.465900e-03 | | Foxb2 | 0.001466 | |
| de novo #208 | 5420 | | | | Foxa1 | 8.615990e-06 | | Foxa3 | 8.615990e-06 | | Foxd1 | 8.615990e-06 | |
| de novo #209 | 6463 | | | | Foxa1 | 2.479240e-07 | | Foxa3 | 1.738970e-06 | | Foxa2 | 1.738970e-06 | |
| de novo #210 | 10097 | | | | Foxa1 | 1.354670e-05 | | Foxa3 | 1.354670e-05 | | Foxd1 | 1.354670e-05 | |
| de novo #211 | 247 | | | | Foxe1 | 1.007770e-03 | | Foxe3 | 1.007770e-03 | | Foxj1 | 2.876580e-03 | |

| pattern | num_seqlets | modisco_cwm_fwd | modisco_cwm_rev | delta_effects | match0 | qval0 | match0_logo | match1 | qval1 | match1_logo | match2 | qval2 | match2_logo |
|---|---|---|---|---|---|---|---|---|---|---|---|---|---|
| de novo #212 | 8419 | | | | Nfia | 4.502530e-05 | | Nfix | 1.257380e-04 | | Nfib | 0.059880 | |
| de novo #213 | 12038 | | | | Nfia | 5.947000e-04 | | Nfix | 1.557050e-03 | | Nfib | 3.631430e-02 | |
| de novo #214 | 10742 | | | | Nfia | 2.474360e-04 | | Nfix | 4.048930e-04 | | Nfib | 5.993450e-02 | |
| de novo #215 | 12936 | | | | Nfia | 4.480720e-05 | | Nfix | 5.994880e-05 | | Nfib | 6.509320e-02 | |
| de novo #216 | 12649 | | | | Nfia | 3.252370e-04 | | Nfix | 5.077820e-04 | | Nfib | 0.057647 | |
| de novo #217 | 12235 | | | | Nfia | 5.496600e-05 | | Nfix | 5.632610e-05 | | Nfib | 7.937560e-02 | |
| de novo #218 | 10021 | | | | Nfia | 3.571630e-06 | | Nfix | 1.253910e-05 | | Nfib | 4.313630e-03 | |
| de novo #219 | 17692 | | | | Nfia | 2.047640e-04 | | Nfix | 3.709800e-04 | | Nfib | 3.466890e-03 | |
| de novo #220 | 46 | | | | Nfia | 1.239430e-04 | | Nfix | 3.531860e-03 | | Nfic | 0.283058 | |
| de novo #221 | 8507 | | | | Nfia | 1.132860e-05 | | Nfix | 3.355680e-05 | | Nfib | 3.203050e-03 | |
| de novo #222 | 10327 | | | | Nfia | 2.323610e-05 | | Nfix | 2.323610e-05 | | Nfib | 1.640720e-01 | |
| de novo #223 | 148 | | | | Nfix | 2.334390e-02 | | Nfia | 2.334390e-02 | | Klf7 | 0.027380 | |
| de novo #224 | 1970 | | | | Nfib | 6.197450e-02 | | Nfia | 6.197450e-02 | | Nfix | 9.233940e-02 | |
| de novo #225 | 10853 | | | | Nfia | 1.765680e-05 | | Nfix | 2.963110e-05 | | Nfib | 1.269940e-01 | |
| de novo #226 | 34 | | | | Nfix | 1.719990e-02 | | Nfia | 5.046160e-02 | | Insm1 | 5.212400e-02 | |

| pattern | num_seqlets | modisco_cwm_fwd | modisco_cwm_rev | delta_effects | match0 | qval0 | match0_logo | match1 | qval1 | match1_logo | match2 | qval2 | match2_logo |
|---|---|---|---|---|---|---|---|---|---|---|---|---|---|
| de novo #227 | 10764 | | | | Nfia | 3.399960e-05 | | Nfix | 7.988520e-05 | | Nfib | 7.969060e-02 | |
| de novo #228 | 535 | | | | Batf3 | 1.201320e-01 | | Jund | 5.808030e-01 | | Fosb | 5.808030e-01 | |
| de novo #229 | 405 | | | | Batf3 | 6.557990e-02 | | Fosb | 1.519830e-01 | | Jund | 2.326260e-01 | |
| de novo #230 | 143 | | | | Elf3 | 1.518150e-01 | | Elf5 | 1.518150e-01 | | Elf1 | 1.518150e-01 | |
| de novo #231 | 270 | | | | Ehf | 1.300130e-04 | | Gabpa | 1.887020e-04 | | Ets1 | 3.497400e-04 | |
| de novo #232 | 45 | | | | Spic | 3.344310e-01 | | Ets2 | 1.000000e+00 | | Ctcf | 1.000000e+00 | |
| de novo #233 | 168 | | | | Elf1 | 4.261680e-04 | | Ehf | 6.256240e-04 | | Elf5 | 1.416600e-03 | |
| de novo #234 | 253 | | | | Ehf | 1.397990e-05 | | Elf5 | 7.486510e-04 | | Elf3 | 7.343490e-03 | |
| de novo #235 | 35 | | | | Zfp296 | 7.098940e-01 | | Zkscan6 | 7.098940e-01 | | Ctcf | 1.000000e+00 | |
| de novo #236 | 180 | | | | Elk1 | 1.507500e-02 | | Elk4 | 1.507500e-02 | | Erg | 1.507500e-02 | |
| de novo #237 | 84 | | | | Elk1 | 3.330640e-02 | | Elk4 | 3.330640e-02 | | Elf1 | 9.536030e-02 | |
| de novo #238 | 31 | | | | Ctcf | 1.000000e+00 | | Hhex | 1.000000e+00 | | Prdm16 | 1.000000e+00 | |
| de novo #239 | 35 | | | | Dmrtc2 | 1.000000e+00 | | Rfx6 | 1.000000e+00 | | Prdm16 | 1.000000e+00 | |
| de novo #240 | 951 | | | | Hoxb9 | 5.755880e-01 | | Cdx4 | 5.755880e-01 | | Hoxd10 | 5.755880e-01 | |
| de novo #241 | 140 | | | | Hoxa13 | 2.975470e-01 | | Hoxd13 | 2.975470e-01 | | Hoxd9 | 3.253570e-01 | |

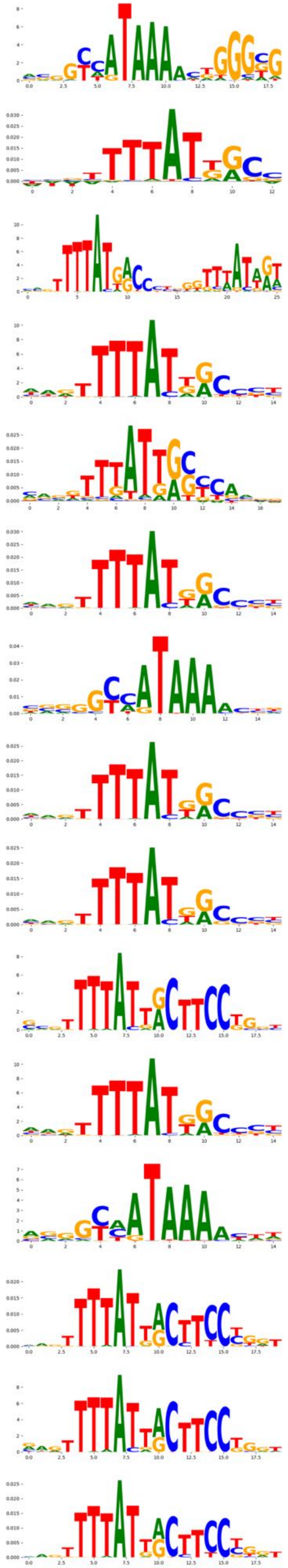
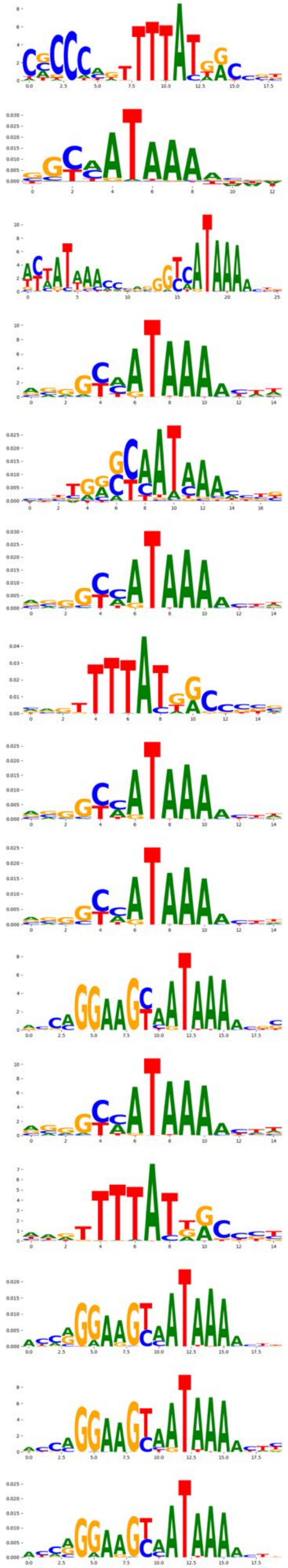
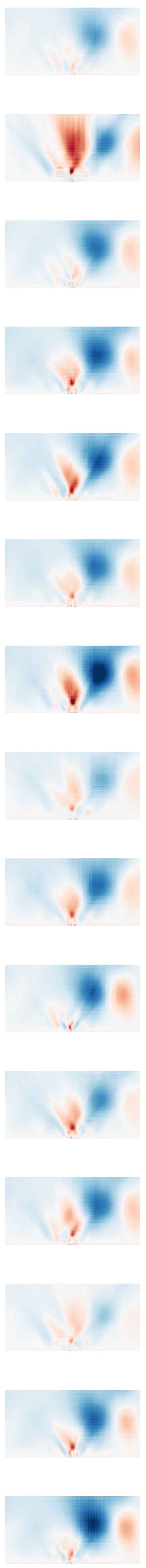
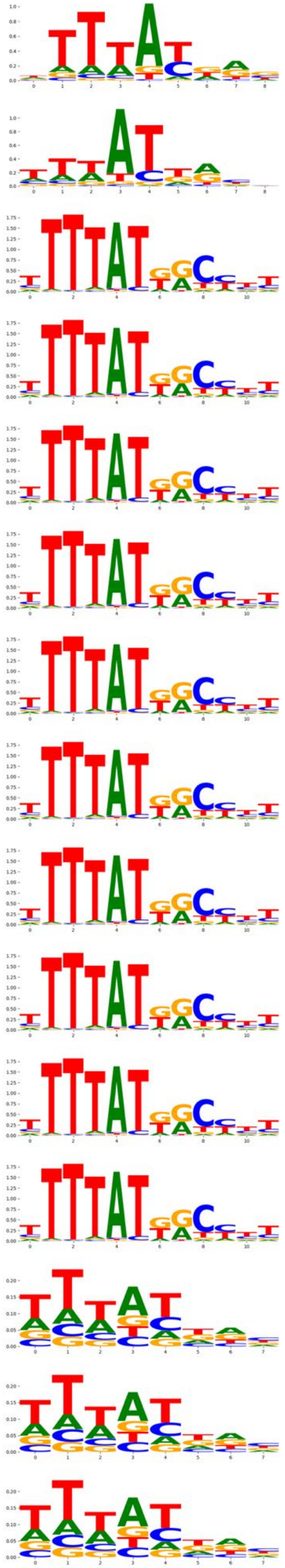
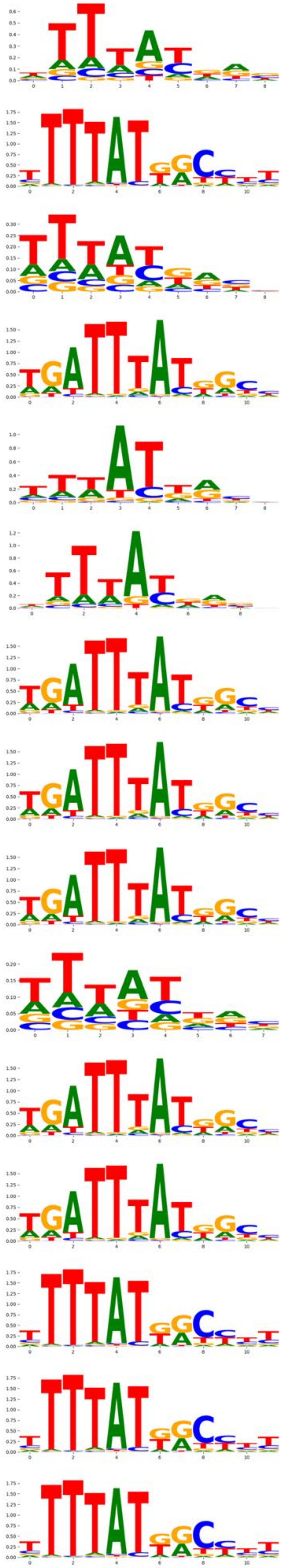
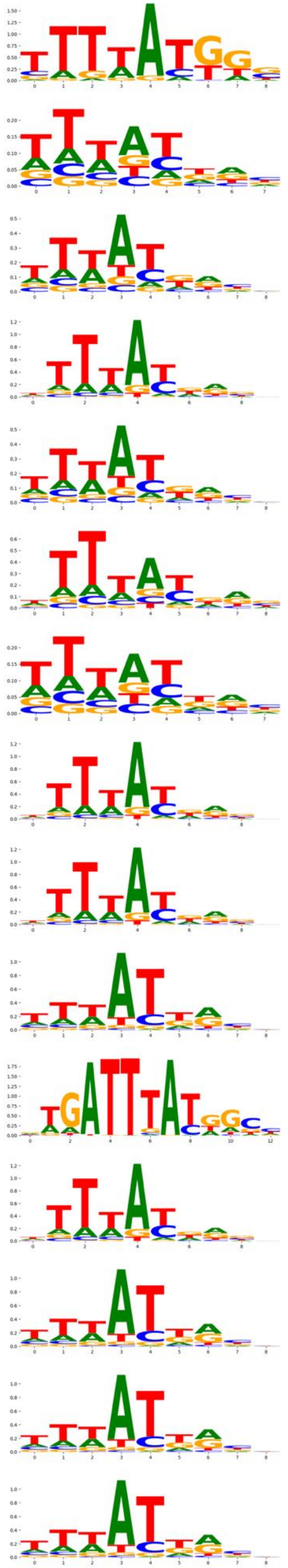

| pattern | num_seqlets | modisco_cwm_fwd | modisco_cwm_rev | delta_effects | match0 | qval0 | match0_logo | match1 | qval1 | match1_logo | match2 | qval2 | match2_logo |
|---|---|---|---|---|---|---|---|---|---|---|---|---|---|
| de novo #242 | 234 | | | | Cdx1 | 5.725080e-05 | | Cdx2 | 5.725080e-05 | | Hoxd10 | 9.505580e-05 | |
| de novo #243 | 439 | | | | Cdx2 | 1.547870e-03 | | Hoxa11 | 1.547870e-03 | | Hoxa10 | 1.600450e-03 | |
| de novo #244 | 21438 | | | | Cdx2 | 1.928070e-04 | | Cdx4 | 7.573610e-04 | | Hoxb13 | 7.573610e-04 | |
| de novo #245 | 275 | | | | Cdx2 | 1.263970e-04 | | Cdx1 | 6.821280e-04 | | Hoxa10 | 4.991530e-03 | |
| de novo #246 | 37351 | | | | Cdx2 | 5.486950e-04 | | Hoxb13 | 5.690130e-04 | | Hoxd13 | 5.690130e-04 | |
| de novo #247 | 35379 | | | | Cdx2 | 3.595990e-05 | | Cdx4 | 6.896940e-05 | | Hoxd10 | 1.585800e-04 | |
| de novo #248 | 33178 | | | | Cdx2 | 2.812740e-04 | | Cdx4 | 9.083030e-04 | | Hoxb13 | 9.083030e-04 | |
| de novo #249 | 31384 | | | | Cdx2 | 6.056690e-04 | | Cdx4 | 6.056690e-04 | | Hoxb13 | 0.000606 | |
| de novo #250 | 36 | | | | Cdx2 | 6.982860e-02 | | Hoxd10 | 6.982860e-02 | | Cdx1 | 8.726180e-02 | |
| de novo #251 | 16803 | | | | Cdx2 | 5.032580e-05 | | Cdx4 | 2.868030e-04 | | Hoxc9 | 2.536080e-03 | |
| de novo #252 | 13651 | | | | Cdx2 | 1.741600e-04 | | Cdx4 | 7.255270e-04 | | Hoxb13 | 7.255270e-04 | |
| de novo #253 | 111 | | | | Hoxd10 | 1.200030e-02 | | Cdx2 | 1.326890e-02 | | Cdx1 | 2.880910e-02 | |
| de novo #254 | 51 | | | | Hoxd10 | 7.643280e-02 | | Cdx2 | 7.643280e-02 | | Cdx1 | 9.589870e-02 | |
| de novo #255 | 110 | | | | Hoxd10 | 1.388310e-02 | | Cdx2 | 1.388310e-02 | | Cdx1 | 3.731710e-02 | |
| de novo #256 | 262 | | | | Cdx1 | 5.384130e-06 | | Cdx2 | 2.597860e-04 | | Hoxa10 | 2.597860e-04 | |

| pattern | num_seqlets | modisco_cwm_fwd | modisco_cwm_rev | delta_effects | match0 | qval0 | match0_logo | match1 | qval1 | match1_logo | match2 | qval2 | match2_logo |
|---|---|---|---|---|---|---|---|---|---|---|---|---|---|
| de novo #257 | 10539 | | | | Cdx2 | 1.347560e-03 | | Hoxb13 | 1.347560e-03 | | Hoxc13 | 1.347560e-03 | |
| de novo #258 | 112 | | | | Dbx1 | 4.137620e-01 | | Hoxd10 | 4.137620e-01 | | Cdc5l | 4.137620e-01 | |
| de novo #259 | 89 | | | | Hoxd10 | 4.451880e-02 | | Cdx1 | 6.468280e-02 | | Hoxb8 | 0.064683 | |
| de novo #260 | 8795 | | | | Cdx2 | 2.697980e-04 | | Cdx4 | 1.667080e-03 | | Hoxd13 | 1.993960e-03 | |
| de novo #261 | 51 | | | | Cdx1 | 3.876740e-05 | | Hoxa11 | 1.214610e-04 | | Hoxa10 | 0.000148 | |
| de novo #262 | 103 | | | | Dbx1 | 4.668750e-01 | | Zfp523 | 4.668750e-01 | | Cdc5l | 4.668750e-01 | |
| de novo #263 | 18363 | | | | Cdx2 | 3.721710e-04 | | Cdx4 | 1.599210e-03 | | Hoxb13 | 1.599210e-03 | |
| de novo #264 | 310 | | | | Hoxa10 | 8.733650e-04 | | Hoxa11 | 8.733650e-04 | | Cdx2 | 8.733650e-04 | |
| de novo #265 | 78 | | | | Dbx1 | 6.472410e-01 | | Onecut2 | 6.472410e-01 | | Cdc5l | 0.647241 | |
| de novo #266 | 124 | | | | Hoxd10 | 1.565380e-02 | | Cdx2 | 1.565380e-02 | | Cdx1 | 0.037209 | |
| de novo #267 | 1352 | | | | Cdx4 | 5.856030e-01 | | Hoxc9 | 5.856030e-01 | | Hoxb9 | 5.856030e-01 | |
| de novo #268 | 37925 | | | | Cdx2 | 4.964230e-05 | | Cdx4 | 1.478590e-04 | | Hoxd10 | 0.000161 | |
| de novo #269 | 184 | | | | Hmg20a | 9.583340e-02 | | Hmg20b | 9.583340e-02 | | Hoxd10 | 1.796610e-01 | |
| de novo #270 | 3467 | | | | Zbtb32 | 1.376540e-01 | | Cdx2 | 2.311090e-01 | | Hoxd10 | 5.991560e-01 | |
| de novo #271 | 68 | | | | Cdx1 | 1.313970e-04 | | Cdx2 | 3.512360e-04 | | Hoxa10 | 3.512360e-04 | |

| pattern | num_seqlets | modisco_cwm_fwd | modisco_cwm_rev | delta_effects | match0 | qval0 | match0_logo | match1 | qval1 | match1_logo | match2 | qval2 | match2_logo |
|---|---|---|---|---|---|---|---|---|---|---|---|---|---|
| de novo #272 | 2895 | | | | Zbtb32 | 1.469910e-01 | | Cdx2 | 1.469910e-01 | | Hoxa13 | 6.600880e-01 | |
| de novo #273 | 157 | | | | Zfp653 | 1.000000e+00 | | Bsx | 1.000000e+00 | | Hoxd9 | 1.000000 | |
| de novo #274 | 299 | | | | Cdx1 | 2.974110e-05 | | Cdx2 | 3.717640e-05 | | Hoxd10 | 0.000098 | |
| de novo #275 | 39407 | | | | Cdx2 | 6.096990e-04 | | Hoxb13 | 7.638510e-04 | | Hoxd13 | 7.638510e-04 | |
| de novo #276 | 585 | | | | Sp2 | 6.319320e-05 | | Klf6 | 6.319320e-05 | | Sp3 | 0.000551 | |
| de novo #277 | 84 | | | | Klf1 | 4.406910e-03 | | Klf4 | 6.206030e-03 | | Klf8 | 0.020599 | |
| de novo #278 | 96 | | | | Klf6 | 7.676270e-04 | | Klf3 | 5.640010e-03 | | Patz1 | 5.640010e-03 | |
| de novo #279 | 213 | | | | Sp3 | 1.252720e-03 | | Klf3 | 1.252720e-03 | | Sp5 | 1.743180e-03 | |
| de novo #280 | 242 | | | | Klf3 | 5.943200e-08 | | Sp3 | 1.201580e-04 | | Klf7 | 1.418650e-04 | |
| de novo #281 | 40567 | | | | Klf3 | 3.027440e-07 | | Klf12 | 5.907840e-05 | | Klf7 | 5.907840e-05 | |
| de novo #282 | 387 | | | | Patz1 | 1.242280e-02 | | Klf2 | 4.117340e-02 | | Wt1 | 4.117340e-02 | |
| de novo #283 | 71 | | | | Zfx | 4.992440e-04 | | Klf6 | 4.992440e-04 | | Sp2 | 0.000499 | |
| de novo #284 | 48189 | | | | Klf3 | 2.593820e-06 | | Klf4 | 3.570540e-05 | | Klf12 | 3.570540e-05 | |
| de novo #285 | 182 | | | | Klf3 | 3.898810e-06 | | Klf12 | 3.898810e-06 | | Klf8 | 3.898810e-06 | |
| de novo #286 | 32 | | | | Klf1 | 4.585310e-03 | | Klf4 | 4.585310e-03 | | Klf12 | 4.719290e-03 | |

| pattern | num_seqlets | modisco_cwm_fwd | modisco_cwm_rev | delta_effects | match0 | qval0 | match0_logo | match1 | qval1 | match1_logo | match2 | qval2 | match2_logo |
|---|---|---|---|---|---|---|---|---|---|---|---|---|---|
| de novo #287 | 46808 | | | | Klf3 | 6.687110e-06 | | Klf4 | 6.687110e-06 | | Klf8 | 0.000009 | |
| de novo #288 | 390 | | | | Klf3 | 7.962920e-07 | | Sp2 | 1.008760e-06 | | Sp3 | 2.495260e-05 | |
| de novo #289 | 420 | | | | Patz1 | 1.168870e-02 | | Klf2 | 4.150580e-02 | | Wt1 | 4.150580e-02 | |
| de novo #290 | 186 | | | | Patz1 | 6.237970e-03 | | Sp5 | 1.123540e-02 | | Wt1 | 5.220390e-02 | |
| de novo #291 | 701 | | | | Klf3 | 1.796740e-04 | | Sp3 | 2.016780e-04 | | Sp2 | 5.998170e-04 | |
| de novo #292 | 338 | | | | Klf12 | 5.998370e-04 | | Klf8 | 5.998370e-04 | | Klf7 | 5.998370e-04 | |
| de novo #293 | 356 | | | | Klf12 | 7.138190e-04 | | Klf8 | 7.138190e-04 | | Klf7 | 0.000788 | |
| de novo #294 | 342 | | | | Klf12 | 4.664650e-04 | | Klf8 | 4.664650e-04 | | Klf7 | 6.496850e-04 | |
| de novo #295 | 47 | | | | Klf12 | 4.779050e-03 | | Klf8 | 4.779050e-03 | | Patz1 | 0.004779 | |
| de novo #296 | 36 | | | | Klf6 | 3.000010e-03 | | Klf1 | 3.000010e-03 | | Klf10 | 3.000010e-03 | |
| de novo #297 | 244 | | | | Sp2 | 9.364210e-04 | | Wt1 | 9.364210e-04 | | Klf15 | 9.364210e-04 | |
| de novo #298 | 450 | | | | Sp2 | 3.083040e-04 | | Klf6 | 3.332050e-04 | | Klf15 | 0.000348 | |
| de novo #299 | 1417 | | | | Klf3 | 1.506870e-06 | | Sp3 | 4.048320e-05 | | Klf6 | 3.412510e-04 | |
| de novo #300 | 121 | | | | Klf6 | 5.740760e-04 | | Patz1 | 2.316560e-03 | | Klf3 | 6.776090e-03 | |
| de novo #301 | 33 | | | | Klf2 | 2.087280e-02 | | Klf3 | 2.236360e-02 | | Klf12 | 2.469430e-02 | |

| pattern | num_seqlets | modisco_cwm_fwd | modisco_cwm_rev | delta_effects | match0 | qval0 | match0_logo | match1 | qval1 | match1_logo | match2 | qval2 | match2_logo |
|---|---|---|---|---|---|---|---|---|---|---|---|---|---|
| de novo #302 | 296 | | | | Sp9 | 3.101230e-04 | | Sp2 | 6.149840e-04 | | Sp4 | 6.149840e-04 | |
| de novo #303 | 399 | | | | Patz1 | 1.000620e-02 | | Klf2 | 3.195810e-02 | | Sp9 | 3.195810e-02 | |
| de novo #304 | 87 | | | | Ehf | 2.261150e-04 | | Sp2 | 1.514020e-03 | | Wt1 | 1.677400e-03 | |
| de novo #305 | 145 | | | | Zfp692 | 8.529210e-03 | | Patz1 | 2.718220e-02 | | Sp2 | 2.718220e-02 | |
| de novo #306 | 54788 | | | | Klf3 | 8.408220e-06 | | Klf4 | 1.161630e-05 | | Klf8 | 1.712320e-05 | |
| de novo #307 | 104 | | | | Klf5 | 1.531390e-02 | | Klf4 | 2.329130e-02 | | Patz1 | 2.329130e-02 | |
| de novo #308 | 59 | | | | Klf6 | 1.556330e-03 | | Klf3 | 4.837700e-03 | | Sp2 | 4.837700e-03 | |
| de novo #309 | 95 | | | | Klf5 | 5.842840e-06 | | Klf4 | 5.842840e-06 | | Klf6 | 5.842840e-06 | |
| de novo #310 | 130 | | | | Klf6 | 5.330380e-04 | | Patz1 | 2.173710e-03 | | Klf3 | 0.006323 | |
| de novo #311 | 398 | | | | Patz1 | 1.449140e-02 | | Klf2 | 4.028890e-02 | | Sp9 | 4.028890e-02 | |
| de novo #312 | 30 | | | | Sp2 | 5.337010e-05 | | Zfx | 5.591200e-05 | | Sp3 | 9.218770e-04 | |
| de novo #313 | 33142 | | | | Klf3 | 9.663900e-08 | | Klf7 | 1.717660e-04 | | Klf12 | 1.717660e-04 | |
| de novo #314 | 30 | | | | Patz1 | 9.942720e-03 | | Elk4 | 2.835490e-02 | | Klf5 | 4.011260e-02 | |
| de novo #315 | 142 | | | | Klf6 | 6.357090e-04 | | Patz1 | 3.308910e-03 | | Klf3 | 8.007750e-03 | |
| de novo #316 | 156 | | | | Sp3 | 3.258880e-06 | | Klf3 | 2.031210e-05 | | Klf15 | 5.775400e-05 | |

| pattern | num_seqlets | modisco_cwm_fwd | modisco_cwm_rev | delta_effects | match0 | qval0 | match0_logo | match1 | qval1 | match1_logo | match2 | qval2 | match2_logo |
|---|---|---|---|---|---|---|---|---|---|---|---|---|---|
| de novo #317 | 58030 | | | | Klf4 | 3.589340e-06 | | Klf3 | 8.746670e-06 | | Klf8 | 0.000009 | |
| de novo #318 | 63 | | | | Zfp692 | 7.848610e-03 | | Klf3 | 1.426330e-02 | | Patz1 | 1.426330e-02 | |
| de novo #319 | 100 | | | | Zfx | 2.638370e-05 | | Sp2 | 3.084470e-03 | | Sp3 | 4.978180e-03 | |
| de novo #320 | 243 | | | | Sp6 | 3.693430e-05 | | Klf14 | 6.686650e-04 | | Klf7 | 0.000669 | |
| de novo #321 | 148 | | | | Sp3 | 1.946130e-03 | | Klf3 | 1.946130e-03 | | Patz1 | 1.946130e-03 | |
| de novo #322 | 21 | | | | Klf1 | 3.362820e-03 | | Klf4 | 3.362820e-03 | | Klf12 | 4.805540e-03 | |
| de novo #323 | 532 | | | | Patz1 | 1.137080e-02 | | Wt1 | 4.860800e-02 | | Klf2 | 4.860800e-02 | |
| de novo #324 | 449 | | | | Klf3 | 6.303230e-05 | | Sp3 | 6.303230e-05 | | Sp2 | 1.742880e-04 | |
| de novo #325 | 469 | | | | Patz1 | 8.489200e-03 | | Klf2 | 4.165850e-02 | | Wt1 | 0.041659 | |
| de novo #326 | 218 | | | | Sp3 | 1.300020e-04 | | Zbtb17 | 1.300020e-04 | | Sp2 | 1.513900e-04 | |
| de novo #327 | 410 | | | | Klf15 | 3.236060e-05 | | Wt1 | 3.236060e-05 | | Sp3 | 3.432940e-04 | |
| de novo #328 | 100 | | | | Klf6 | 8.743660e-04 | | Klf3 | 5.966710e-03 | | Patz1 | 0.005967 | |
| de novo #329 | 594 | | | | Klf3 | 3.733180e-05 | | Sp3 | 6.046500e-05 | | Sp2 | 2.037850e-04 | |
| de novo #330 | 55021 | | | | Klf4 | 6.384850e-06 | | Klf3 | 9.681040e-06 | | Klf8 | 9.681040e-06 | |
| de novo #331 | 47 | | | | Klf5 | 5.734230e-04 | | Klf8 | 5.734230e-04 | | Klf1 | 5.734230e-04 | |

| pattern | num_seqlets | modisco_cwm_fwd | modisco_cwm_rev | delta_effects | match0 | qval0 | match0_logo | match1 | qval1 | match1_logo | match2 | qval2 | match2_logo |
|---|---|---|---|---|---|---|---|---|---|---|---|---|---|
| de novo #332 | 24 | | | | Nrf1 | 6.955650e-04 | | Klf12 | 2.876500e-02 | | Klf7 | 3.209170e-02 | |
| de novo #333 | 235 | | | | Patz1 | 5.922350e-04 | | Sp3 | 5.922350e-04 | | Klf3 | 0.000852 | |
| de novo #334 | 44793 | | | | Klf3 | 4.896940e-07 | | Klf6 | 2.441620e-05 | | Klf4 | 2.441620e-05 | |
| de novo #335 | 540 | | | | Wt1 | 7.636360e-05 | | Klf15 | 7.636360e-05 | | Sp3 | 7.636360e-05 | |
| de novo #336 | 449 | | | | Klf15 | 3.947330e-04 | | Sp3 | 3.947330e-04 | | Klf6 | 3.947330e-04 | |
| de novo #337 | 36734 | | | | Klf3 | 4.489610e-07 | | Klf7 | 1.232920e-04 | | Klf12 | 1.232920e-04 | |
| de novo #338 | 66 | | | | Klf6 | 1.244180e-03 | | Klf3 | 4.912110e-03 | | Sp2 | 5.818060e-03 | |
| de novo #339 | 166 | | | | Patz1 | 6.176580e-04 | | Sp3 | 6.176580e-04 | | Klf3 | 7.126440e-04 | |
| de novo #340 | 413 | | | | Patz1 | 1.403570e-02 | | Klf2 | 4.052360e-02 | | Sp9 | 4.052360e-02 | |
| de novo #341 | 94 | | | | Klf6 | 9.827730e-04 | | Klf3 | 4.989960e-03 | | Patz1 | 4.989960e-03 | |
| de novo #342 | 435 | | | | Sp3 | 1.130670e-04 | | Sp2 | 1.130670e-04 | | Klf10 | 1.078310e-03 | |
| de novo #343 | 45935 | | | | Klf3 | 3.418860e-06 | | Klf4 | 1.837130e-05 | | Klf8 | 2.825550e-05 | |
| de novo #344 | 371 | | | | Sp2 | 1.219880e-04 | | Sp3 | 1.219880e-04 | | Klf6 | 1.050030e-03 | |
| de novo #345 | 245 | | | | Sp9 | 4.630510e-04 | | Sp4 | 6.761930e-04 | | Sp2 | 6.761930e-04 | |
| de novo #346 | 169 | | | | Klf6 | 7.421430e-04 | | Wt1 | 7.421430e-04 | | Sp2 | 7.421430e-04 | |

| pattern | num_seqlets | modisco_cwm_fwd | modisco_cwm_rev | delta_effects | match0 | qval0 | match0_logo | match1 | qval1 | match1_logo | match2 | qval2 | match2_logo |
|---|---|---|---|---|---|---|---|---|---|---|---|---|---|
| de novo #347 | 764 | | | | Sp3 | 8.996250e-05 | | Sp2 | 2.349200e-04 | | Zbtb17 | 0.000969 | |
| de novo #348 | 428 | | | | Klf10 | 1.443460e-05 | | Sp3 | 1.443460e-05 | | Sp2 | 2.102810e-05 | |
| de novo #349 | 52525 | | | | Klf3 | 5.610830e-06 | | Klf4 | 5.610830e-06 | | Klf8 | 7.988790e-06 | |
| de novo #350 | 245 | | | | Sp3 | 1.996150e-03 | | Klf3 | 1.996150e-03 | | Sp2 | 1.996150e-03 | |
| de novo #351 | 23 | | | | Klf4 | 9.555230e-03 | | Patz1 | 9.555230e-03 | | Klf7 | 1.289980e-02 | |
| de novo #352 | 251 | | | | Patz1 | 1.469830e-04 | | Sp3 | 4.571180e-04 | | Klf3 | 5.614240e-04 | |
| de novo #353 | 215 | | | | Klf4 | 3.044940e-06 | | Klf8 | 5.301790e-06 | | Klf5 | 0.000008 | |
| de novo #354 | 479 | | | | Patz1 | 1.039740e-02 | | Wt1 | 4.570020e-02 | | Klf2 | 0.045700 | |
| de novo #355 | 175 | | | | Sp3 | 2.013640e-03 | | Klf3 | 2.013640e-03 | | Patz1 | 2.013640e-03 | |
| de novo #356 | 419 | | | | Klf10 | 1.660380e-05 | | Sp3 | 1.660380e-05 | | Sp2 | 1.898230e-05 | |
| de novo #357 | 39601 | | | | Klf3 | 3.935730e-07 | | Klf7 | 7.105330e-05 | | Klf8 | 7.105330e-05 | |
| de novo #358 | 631 | | | | Sp2 | 3.185080e-04 | | Sp3 | 4.855340e-04 | | Zbtb17 | 9.342300e-04 | |
| de novo #359 | 437 | | | | Patz1 | 1.274330e-02 | | Klf2 | 3.531620e-02 | | Sp9 | 3.531620e-02 | |
| de novo #360 | 1636 | | | | Sp3 | 3.243660e-07 | | Sp2 | 2.988540e-06 | | Klf3 | 1.354700e-04 | |
| de novo #361 | 361 | | | | Sp3 | 3.123660e-04 | | Sp2 | 3.123660e-04 | | Klf15 | 3.123660e-04 | |

| pattern | num_seqlets | modisco_cwm_fwd | modisco_cwm_rev | delta_effects | match0 | qval0 | match0_logo | match1 | qval1 | match1_logo | match2 | qval2 | match2_logo |
|---|---|---|---|---|---|---|---|---|---|---|---|---|---|
| de novo #362 | 69 | | | | Fosb | 2.847850e-01 | | Hoxc5 | 2.847850e-01 | | Bsx | 2.847850e-01 | |
| de novo #363 | 7704 | | | | Pdx1 | 5.690690e-01 | | Arid3b | 5.690690e-01 | | Otx2 | 5.690690e-01 | |
| de novo #364 | 5932 | | | | Pdx1 | 5.586630e-01 | | Nanog | 5.586630e-01 | | Hoxa4 | 0.558663 | |
| de novo #365 | 4732 | | | | Pdx1 | 4.854910e-01 | | Arid3b | 4.854910e-01 | | Nanog | 4.854910e-01 | |
| de novo #366 | 3483 | | | | Pdx1 | 5.476390e-01 | | Nanog | 5.476390e-01 | | Hoxa4 | 5.476390e-01 | |
| de novo #367 | 7278 | | | | Pdx1 | 5.526290e-01 | | Nanog | 5.526290e-01 | | Arid3b | 5.526290e-01 | |
| de novo #368 | 7917 | | | | Pdx1 | 5.599420e-01 | | Hoxa4 | 5.599420e-01 | | Nanog | 0.559942 | |
| de novo #369 | 210 | | | | Hoxc10 | 1.695080e-01 | | Fosb | 1.695080e-01 | | Hoxa4 | 1.695080e-01 | |
| de novo #370 | 6445 | | | | Pdx1 | 5.397540e-01 | | Arid3b | 5.397540e-01 | | Pou4f1 | 5.397540e-01 | |
| de novo #371 | 7409 | | | | Pdx1 | 5.722250e-01 | | Nanog | 5.722250e-01 | | Hoxa4 | 5.722250e-01 | |
| de novo #372 | 4356 | | | | Pdx1 | 5.616770e-01 | | Nanog | 5.616770e-01 | | Arid3b | 5.616770e-01 | |
| de novo #373 | 3683 | | | | Pdx1 | 5.483540e-01 | | Nanog | 5.483540e-01 | | Arid3b | 5.483540e-01 | |
| de novo #374 | 227 | | | | Arid5a | 8.572820e-01 | | Foxl2 | 8.572820e-01 | | Hmg20a | 8.572820e-01 | |
| de novo #375 | 6540 | | | | Pdx1 | 5.558850e-01 | | Nanog | 5.558850e-01 | | Hoxa4 | 5.558850e-01 | |
| de novo #376 | 5979 | | | | Pdx1 | 5.317380e-01 | | Nanog | 5.317380e-01 | | Arid3b | 5.317380e-01 | |

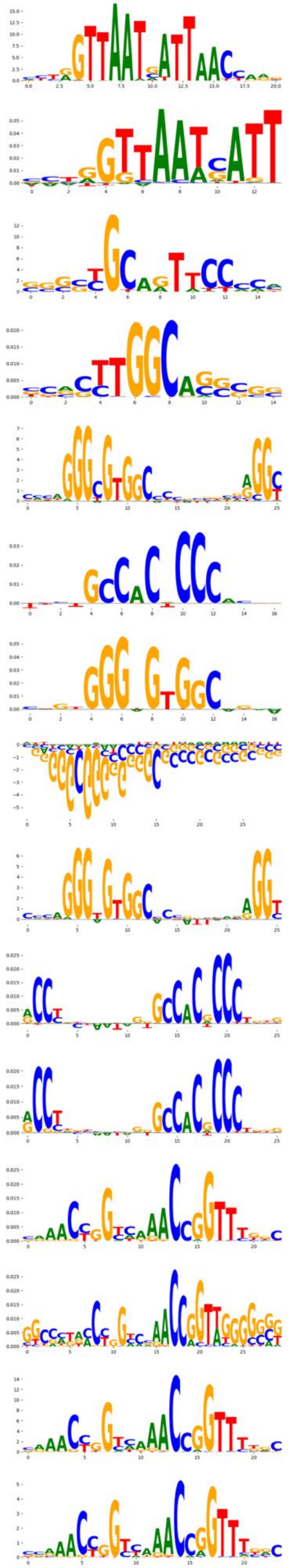
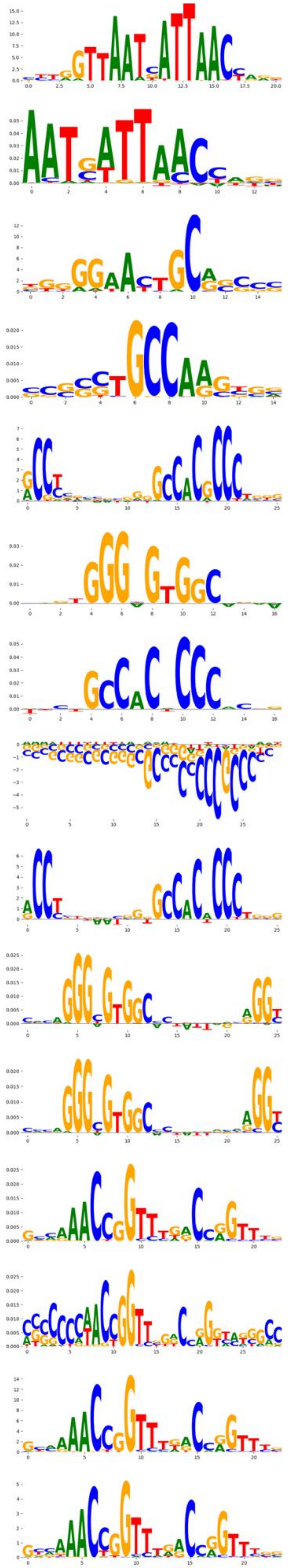
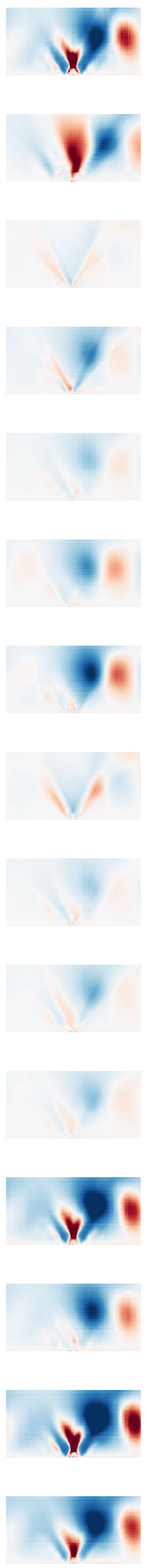
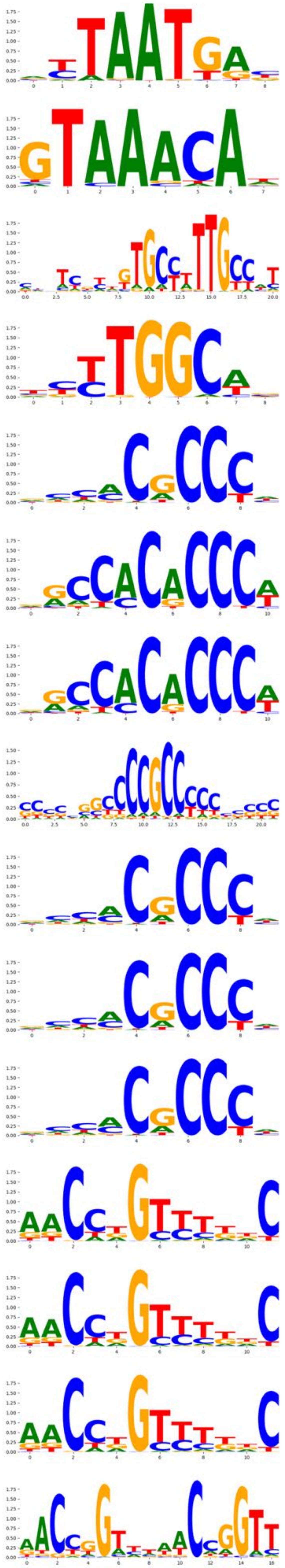
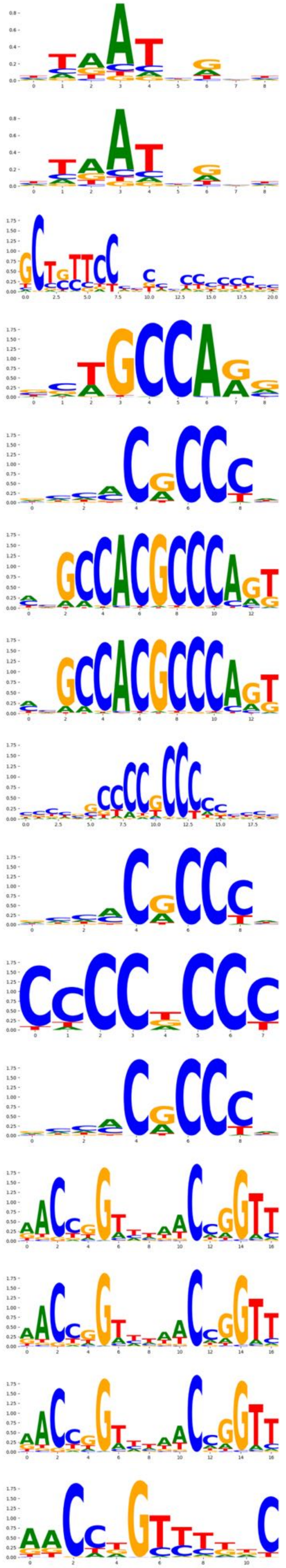
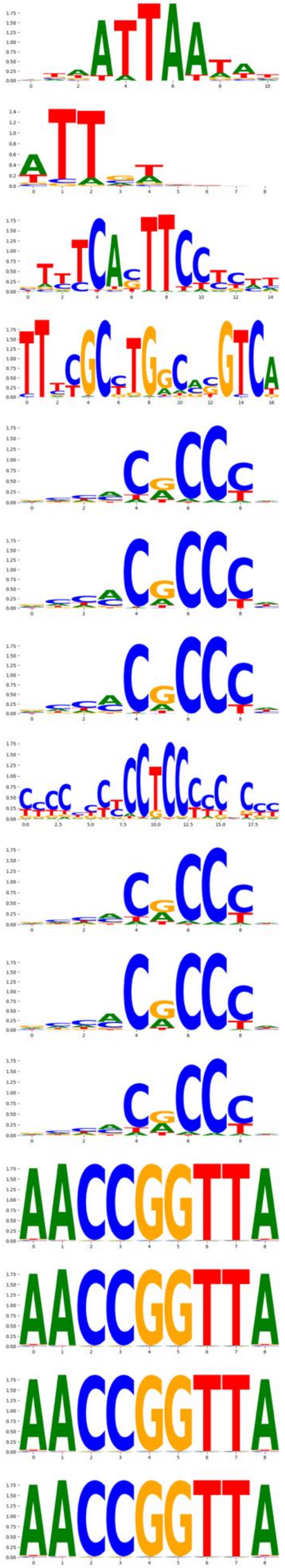

| pattern | num_seqlets | modisco_cwm_fwd | modisco_cwm_rev | delta_effects | match0 | qval0 | match0_logo | match1 | qval1 | match1_logo | match2 | qval2 | match2_logo |
|---|---|---|---|---|---|---|---|---|---|---|---|---|---|
| de novo #377 | 87 | | | | Foxd2 | 8.650090e-01 | | Nanog | 8.650090e-01 | | Sox15 | 0.865009 | |
| de novo #378 | 138 | | | | Zik1 | 1.000000e+00 | | Zfp341 | 1.000000e+00 | | Bcl11a | 1.000000e+00 | |
| de novo #379 | 3110 | | | | Nfic | 7.198460e-03 | | Nfib | 4.216370e-02 | | Zbtb49 | 0.531475 | |
| de novo #380 | 481 | | | | Klf7 | 3.325390e-01 | | Klf12 | 3.325390e-01 | | Klf8 | 3.325390e-01 | |
| de novo #381 | 57 | | | | Klf1 | 4.075540e-02 | | Sp6 | 4.075540e-02 | | Klf7 | 4.075540e-02 | |
| de novo #382 | 56 | | | | Klf1 | 4.190720e-02 | | Sp6 | 4.190720e-02 | | Klf7 | 4.190720e-02 | |
| de novo #383 | 105 | | | | Sp2 | 4.702370e-05 | | Sp3 | 4.702370e-05 | | Wt1 | 7.348190e-04 | |
| de novo #384 | 354 | | | | Klf7 | 5.726030e-01 | | Klf12 | 5.726030e-01 | | Klf8 | 5.726030e-01 | |
| de novo #385 | 828 | | | | Klf7 | 5.209680e-01 | | Patz1 | 5.209680e-01 | | Klf12 | 5.209680e-01 | |
| de novo #386 | 665 | | | | Klf7 | 4.182910e-01 | | Klf12 | 4.182910e-01 | | Klf8 | 4.182910e-01 | |
| de novo #387 | 2297 | | | | Grhl2 | 1.251430e-06 | | Grhl1 | 1.543450e-05 | | Tfcp2 | 1.148610e-02 | |
| de novo #388 | 26 | | | | Grhl2 | 9.361970e-03 | | Grhl1 | 2.466000e-02 | | Tfcp2 | 6.139840e-02 | |
| de novo #389 | 1870 | | | | Grhl2 | 5.657740e-07 | | Grhl1 | 1.390640e-05 | | Tfcp2 | 1.129270e-02 | |
| de novo #390 | 444 | | | | Grhl1 | 2.144290e-06 | | Grhl2 | 8.318640e-05 | | Tfcp2 | 1.182690e-02 | |
| de novo #391 | 904 | | | | Grhl1 | 1.305210e-05 | | Grhl2 | 1.945310e-05 | | Tfcp2 | 1.154860e-02 | |

| pattern | num_seqlets | modisco_cwm_fwd | modisco_cwm_rev | delta_effects | match0 | qval0 | match0_logo | match1 | qval1 | match1_logo | match2 | qval2 | match2_logo |
|---|---|---|---|---|---|---|---|---|---|---|---|---|---|
| de novo #392 | 897 | | | | Grhl2 | 8.416310e-06 | | Grhl1 | 8.416310e-06 | | Tfcp2 | 1.339060e-02 | |
| de novo #393 | 3357 | | | | Grhl2 | 1.497430e-07 | | Grhl1 | 1.635860e-05 | | Tfcp2 | 0.012409 | |
| de novo #394 | 943 | | | | Grhl2 | 1.854410e-05 | | Grhl1 | 1.854410e-05 | | Tfcp2 | 1.179520e-02 | |
| de novo #395 | 1890 | | | | Grhl2 | 3.427030e-07 | | Grhl1 | 1.618990e-05 | | Tfcp2 | 1.157100e-02 | |
| de novo #396 | 34 | | | | Grhl1 | 7.836940e-04 | | Fli1 | 1.602090e-01 | | Ets1 | 1.602090e-01 | |
| de novo #397 | 2072 | | | | Grhl2 | 1.250760e-06 | | Grhl1 | 2.157480e-05 | | Tfcp2 | 1.403330e-02 | |
| de novo #398 | 3335 | | | | Grhl2 | 8.281140e-07 | | Grhl1 | 1.900960e-05 | | Tfcp2 | 1.006200e-02 | |
| de novo #399 | 298 | | | | Grhl1 | 1.026640e-03 | | Grhl2 | 1.026640e-03 | | Tfcp2 | 2.847210e-02 | |
| de novo #400 | 1203 | | | | Grhl2 | 6.621740e-06 | | Grhl1 | 1.181850e-05 | | Tfcp2 | 0.012632 | |
| de novo #401 | 93 | | | | Sox9 | 2.166990e-09 | | Sox3 | 9.277680e-03 | | Sox6 | 9.277680e-03 | |
| de novo #402 | 161 | | | | Zkscan17 | 1.000000e+00 | | NaN | NaN | | NaN | NaN | |
| de novo #403 | 310 | | | | Zbtb3 | 1.000000e+00 | | Ddit3 | 1.000000e+00 | | Atf4 | 1.000000e+00 | |
| de novo #404 | 3745 | | | | Nrf1 | 3.821710e-07 | | Egr1 | 1.000000e+00 | | Egr4 | 1.000000e+00 | |
| de novo #405 | 3776 | | | | Nrf1 | 4.693570e-06 | | Egr1 | 9.253110e-01 | | Mtf1 | 1.000000e+00 | |
| de novo #406 | 4107 | | | | Nrf1 | 1.837240e-05 | | Egr1 | 8.938490e-01 | | Mtf1 | 8.938490e-01 | |

| pattern | num_seqlets | modisco_cwm_fwd | modisco_cwm_rev | delta_effects | match0 | qval0 | match0_logo | match1 | qval1 | match1_logo | match2 | qval2 | match2_logo |
|---|---|---|---|---|---|---|---|---|---|---|---|---|---|
| de novo #407 | 3031 | | | | Nrf1 | 5.586510e-05 | | Hes7 | 9.702470e-01 | | Zfx | 9.702470e-01 | |
| de novo #408 | 70 | | | | Nrf1 | 2.351130e-02 | | Zfx | 4.076190e-02 | | Sp2 | 0.232095 | |
| de novo #409 | 27 | | | | Nrf1 | 9.136930e-03 | | Mnt | 8.305800e-01 | | Myc | 8.489310e-01 | |
| de novo #410 | 2942 | | | | Nrf1 | 9.457550e-05 | | Hes7 | 9.323810e-01 | | Zfx | 9.323810e-01 | |
| de novo #411 | 3892 | | | | Nrf1 | 2.580790e-07 | | Egr1 | 1.000000e+00 | | Egr2 | 1.000000e+00 | |
| de novo #412 | 2598 | | | | Nrf1 | 9.453140e-05 | | Zfx | 8.468230e-01 | | Hes7 | 0.846823 | |
| de novo #413 | 3216 | | | | Nrf1 | 5.593980e-05 | | Hes7 | 9.727930e-01 | | Zfx | 0.972793 | |
| de novo #414 | 3043 | | | | Nrf1 | 1.331300e-05 | | Egr1 | 8.532380e-01 | | Mtf1 | 1.000000e+00 | |
| de novo #415 | 2686 | | | | Nrf1 | 1.642270e-05 | | Egr1 | 8.151840e-01 | | Mtf1 | 1.000000e+00 | |
| de novo #416 | 4233 | | | | Nrf1 | 9.196980e-06 | | Mtf1 | 8.002040e-01 | | Egr1 | 8.002040e-01 | |
| de novo #417 | 51 | | | | Zfp566 | 6.847360e-03 | | Nrf1 | 7.865330e-03 | | Zfp467 | 7.865330e-03 | |
| de novo #418 | 46 | | | | Nrf1 | 1.539760e-03 | | Zfp467 | 1.539760e-03 | | Wt1 | 1.539760e-03 | |
| de novo #419 | 55 | | | | Plagl1 | 7.459520e-03 | | Zfx | 1.100360e-02 | | Nrf1 | 2.097070e-02 | |
| de novo #420 | 3709 | | | | Nrf1 | 7.422720e-06 | | Egr1 | 9.324690e-01 | | Mtf1 | 1.000000e+00 | |
| de novo #421 | 128 | | | | Nrf1 | 1.399850e-02 | | Mxd1 | 1.399850e-02 | | Mxi1 | 0.013999 | |

| pattern | num_seqlets | modisco_cwm_fwd | modisco_cwm_rev | delta_effects | match0 | qval0 | match0_logo | match1 | qval1 | match1_logo | match2 | qval2 | match2_logo |
|---|---|---|---|---|---|---|---|---|---|---|---|---|---|
| de novo #422 | 1007 | | | | Foxd2 | 7.389820e-01 | | Foxg1 | 7.389820e-01 | | Foxi2 | 0.738982 | |
| de novo #423 | 633 | | | | Foxd2 | 7.053940e-01 | | Foxg1 | 7.053940e-01 | | Foxi2 | 7.053940e-01 | |
| de novo #424 | 640 | | | | Foxd2 | 7.752200e-01 | | Foxg1 | 7.752200e-01 | | Nkx61 | 7.752200e-01 | |
| de novo #425 | 1069 | | | | Foxd2 | 7.763290e-01 | | Foxg1 | 7.763290e-01 | | Foxi2 | 7.763290e-01 | |
| de novo #426 | 129 | | | | Foxd2 | 6.637840e-01 | | Arid5a | 6.637840e-01 | | Nkx61 | 6.637840e-01 | |
| de novo #427 | 10619 | | | | Ctcfl | 5.226180e-02 | | Zfx | 5.226180e-02 | | Sp2 | 5.226180e-02 | |
| de novo #428 | 407 | | | | Sp2 | 1.156330e-08 | | Sp3 | 1.286370e-07 | | Klf3 | 1.518830e-04 | |
| de novo #429 | 1684 | | | | Sp2 | 2.919030e-02 | | Sp3 | 2.919030e-02 | | Zfx | 7.906170e-02 | |
| de novo #430 | 192 | | | | Sp2 | 3.486450e-05 | | Sp3 | 3.486450e-05 | | Wt1 | 6.440840e-04 | |
| de novo #431 | 1301 | | | | Zfx | 2.648650e-04 | | Sp2 | 5.672660e-04 | | Zfp64 | 5.672660e-04 | |
| de novo #432 | 1487 | | | | Sp2 | 6.418890e-08 | | Sp3 | 1.040180e-05 | | Zfx | 1.251400e-04 | |
| de novo #433 | 621 | | | | Sp2 | 2.650250e-03 | | Sp3 | 2.905590e-02 | | Zfx | 0.029056 | |
| de novo #434 | 331 | | | | Sp2 | 8.026460e-06 | | Sp3 | 1.509010e-05 | | Zfx | 1.187160e-04 | |
| de novo #435 | 955 | | | | Zbtb6 | 1.877220e-01 | | Mxd1 | 1.877220e-01 | | Mxi1 | 1.877220e-01 | |
| de novo #436 | 259 | | | | Sp2 | 1.003340e-05 | | Sp3 | 3.522170e-05 | | Zfx | 3.977780e-03 | |

| pattern | num_seqlets | modisco_cwm_fwd | modisco_cwm_rev | delta_effects | match0 | qval0 | match0_logo | match1 | qval1 | match1_logo | match2 | qval2 | match2_logo |
|---|---|---|---|---|---|---|---|---|---|---|---|---|---|
| de novo #437 | 189 | | | | Sp2 | 1.837740e-06 | | Sp3 | 4.224710e-05 | | Zfx | 3.714830e-04 | |
| de novo #438 | 689 | | | | Sp2 | 7.069750e-02 | | Klf3 | 7.910440e-02 | | Sp3 | 0.079104 | |
| de novo #439 | 33 | | | | Zfx | 5.831020e-03 | | Sp2 | 5.831020e-03 | | Zfp64 | 0.005831 | |
| de novo #440 | 516 | | | | Sp2 | 4.775550e-06 | | Sp3 | 3.418300e-05 | | Wt1 | 2.580020e-04 | |
| de novo #441 | 104 | | | | Sp2 | 3.007390e-04 | | Zfx | 1.882490e-03 | | Sp3 | 1.078100e-02 | |
| de novo #442 | 67 | | | | Sp2 | 1.844670e-06 | | Sp3 | 6.892770e-06 | | Zfx | 6.738980e-04 | |
| de novo #443 | 115 | | | | Sp2 | 3.585010e-07 | | Sp3 | 7.071260e-06 | | Zfp566 | 3.171660e-04 | |
| de novo #444 | 269 | | | | Sp2 | 7.158690e-07 | | Sp3 | 9.802470e-06 | | Zfx | 1.055490e-03 | |
| de novo #445 | 160 | | | | Sp2 | 6.546130e-05 | | Sp3 | 5.925930e-04 | | Klf3 | 2.847250e-03 | |
| de novo #446 | 457 | | | | Sp2 | 2.003380e-06 | | Sp3 | 2.817990e-05 | | Wt1 | 1.773620e-03 | |
| de novo #447 | 203 | | | | Sp2 | 7.898380e-07 | | Sp3 | 1.555440e-05 | | Klf3 | 1.182870e-03 | |
| de novo #448 | 139 | | | | Zfp692 | 2.199180e-01 | | Zfp335 | 6.808590e-01 | | Zfp64 | 6.808590e-01 | |
| de novo #449 | 51 | | | | Taf1 | 3.335580e-05 | | Mbtps2 | 3.777380e-05 | | Yy1 | 3.777380e-05 | |
| de novo #450 | 472 | | | | Smad1 | 4.991070e-02 | | Smad5 | 4.991070e-02 | | Mbd2 | 6.723600e-02 | |
| de novo #451 | 74 | | | | Sp2 | 2.378080e-04 | | Sp3 | 4.315520e-03 | | Zfx | 7.060010e-03 | |

| pattern | num_seqlets | modisco_cwm_fwd | modisco_cwm_rev | delta_effects | match0 | qval0 | match0_logo | match1 | qval1 | match1_logo | match2 | qval2 | match2_logo |
|---|---|---|---|---|---|---|---|---|---|---|---|---|---|
| de novo #452 | 29 | | | | Sp3 | 3.044650e-03 | | Zfp64 | 3.044650e-03 | | Zfx | 3.561430e-03 | |
| de novo #453 | 384 | | | | Sp2 | 1.537770e-07 | | Sp3 | 1.673940e-05 | | Zbtb17 | 3.229580e-04 | |
| de novo #454 | 35 | | | | Sp2 | 8.278650e-04 | | Sp3 | 3.292050e-03 | | Zfx | 3.292050e-03 | |
| de novo #455 | 1624 | | | | Sp2 | 2.080270e-04 | | Zfx | 3.827530e-04 | | Sp3 | 1.265280e-03 | |
| de novo #456 | 240 | | | | Zfx | 8.374740e-05 | | Sp2 | 2.329670e-04 | | Sp3 | 7.958590e-04 | |
| de novo #457 | 1602 | | | | Sp2 | 3.238360e-08 | | Sp3 | 4.511210e-06 | | Zfp566 | 7.587850e-04 | |
| de novo #458 | 1086 | | | | Sp2 | 5.507110e-05 | | Sp3 | 1.269890e-03 | | Zfx | 1.675190e-03 | |
| de novo #459 | 24 | | | | Sp2 | 1.902450e-05 | | Sp3 | 4.164660e-04 | | Zfx | 1.019300e-03 | |
| de novo #460 | 540 | | | | Sp2 | 1.205750e-05 | | Sp3 | 3.276170e-05 | | Zfx | 1.133950e-04 | |
| de novo #461 | 311 | | | | Sp2 | 2.510890e-05 | | Sp3 | 4.483830e-05 | | Wt1 | 2.742480e-04 | |
| de novo #462 | 4912 | | | | Zfx | 2.282850e-03 | | Sp2 | 2.282850e-03 | | Sp3 | 8.065960e-03 | |
| de novo #463 | 351 | | | | Sp2 | 6.917690e-07 | | Sp3 | 5.542800e-06 | | Zbtb17 | 1.008880e-04 | |
| de novo #464 | 807 | | | | Sp2 | 1.617090e-04 | | Zfx | 1.617090e-04 | | Sp3 | 5.658190e-04 | |
| de novo #465 | 160 | | | | Sp2 | 1.130950e-04 | | Zfx | 7.205940e-04 | | Sp3 | 7.205940e-04 | |
| de novo #466 | 2256 | | | | Sp3 | 1.864920e-04 | | Sp2 | 1.864920e-04 | | Zfx | 4.974510e-03 | |

| pattern | num_seqlets | modisco_cwm_fwd | modisco_cwm_rev | delta_effects | match0 | qval0 | match0_logo | match1 | qval1 | match1_logo | match2 | qval2 | match2_logo |
|---|---|---|---|---|---|---|---|---|---|---|---|---|---|
| de novo #467 | 2909 | | | | Sp2 | 1.211360e-03 | | Ctcfl | 1.211360e-03 | | Zfx | 1.211360e-03 | |
| de novo #468 | 490 | | | | Sp2 | 2.815930e-03 | | Zfx | 2.815930e-03 | | Sp3 | 6.204260e-03 | |
| de novo #469 | 7659 | | | | Sp2 | 4.800970e-07 | | Sp3 | 1.110760e-05 | | Zfx | 1.639090e-04 | |
| de novo #470 | 519 | | | | Sp2 | 9.048350e-05 | | Sp3 | 1.029990e-04 | | Zfp64 | 2.791350e-04 | |
| de novo #471 | 401 | | | | Sp2 | 4.525810e-06 | | Sp3 | 9.712550e-05 | | Klf3 | 3.579060e-03 | |
| de novo #472 | 119 | | | | Creb3 | 4.107630e-02 | | Gm28047 | 6.015100e-02 | | Sp3 | 6.015100e-02 | |
| de novo #473 | 309 | | | | Myc | 2.893450e-02 | | Sp3 | 2.893450e-02 | | Sp2 | 2.893450e-02 | |
| de novo #474 | 888 | | | | Sp2 | 6.290030e-02 | | Mbd2 | 6.290030e-02 | | Klf3 | 6.290030e-02 | |
| de novo #475 | 1839 | | | | Mbd2 | 1.447300e-01 | | Sp2 | 1.447300e-01 | | Smad1 | 0.144730 | |
| de novo #476 | 35 | | | | Sp2 | 5.155330e-05 | | Sp3 | 4.811090e-04 | | Zfx | 4.811090e-04 | |
| de novo #477 | 552 | | | | Nrf1 | 4.047090e-01 | | Tfdp1 | 4.047090e-01 | | Tfdp2 | 4.047090e-01 | |
| de novo #478 | 308 | | | | Sp2 | 1.078270e-04 | | Sp3 | 4.135760e-04 | | Wt1 | 4.751010e-04 | |
| de novo #479 | 1770 | | | | Sp2 | 2.207820e-02 | | Zbtb6 | 8.800450e-02 | | Nrf1 | 0.088004 | |
| de novo #480 | 2569 | | | | Sp2 | 1.149380e-04 | | Zfp566 | 1.336010e-03 | | Sp3 | 1.336010e-03 | |
| de novo #481 | 656 | | | | Sp2 | 1.037480e-06 | | Sp3 | 1.157920e-05 | | Wt1 | 8.850050e-05 | |

| pattern | num_seqlets | modisco_cwm_fwd | modisco_cwm_rev | delta_effects | match0 | qval0 | match0_logo | match1 | qval1 | match1_logo | match2 | qval2 | match2_logo |
|---|---|---|---|---|---|---|---|---|---|---|---|---|---|
| de novo #482 | 1210 | | | | Zbtb14 | 3.371190e-01 | | Nrf1 | 3.371190e-01 | | Hes1 | 3.371190e-01 | |
| de novo #483 | 548 | | | | Sp2 | 8.628960e-04 | | Zfx | 8.628960e-04 | | Sp3 | 1.330830e-03 | |
| de novo #484 | 416 | | | | Sp3 | 7.836470e-07 | | Sp2 | 1.142720e-06 | | Zfp341 | 2.238730e-05 | |
| de novo #485 | 587 | | | | Sp2 | 1.660750e-07 | | Sp3 | 3.810260e-07 | | Wt1 | 1.227350e-04 | |
| de novo #486 | 118 | | | | Sp2 | 1.500640e-05 | | Sp3 | 1.418300e-04 | | Zfx | 1.906300e-03 | |
| de novo #487 | 17465 | | | | Sp2 | 1.048780e-03 | | Zfx | 1.048780e-03 | | Sp3 | 4.453680e-03 | |
| de novo #488 | 4867 | | | | Zfx | 6.824050e-03 | | Sp2 | 6.824050e-03 | | Mxd1 | 6.824050e-03 | |
| de novo #489 | 28 | | | | Sp2 | 2.940820e-04 | | Sp3 | 3.188680e-04 | | Zfx | 1.501660e-03 | |
| de novo #490 | 3741 | | | | Sp2 | 9.483770e-06 | | Sp3 | 4.639050e-04 | | Zfx | 1.465260e-03 | |
| de novo #491 | 2276 | | | | Sp2 | 3.660480e-08 | | Sp3 | 3.569200e-06 | | Zfx | 2.023900e-04 | |
| de novo #492 | 72 | | | | Zfx | 9.778580e-02 | | Zbtb14 | 9.778580e-02 | | Hes1 | 9.778580e-02 | |
| de novo #493 | 3741 | | | | Mbd2 | 1.043410e-01 | | Sp2 | 1.043410e-01 | | Dnmt1 | 1.043410e-01 | |
| de novo #494 | 81 | | | | Taf1 | 2.038650e-04 | | Mbtps2 | 2.760060e-04 | | Yy1 | 2.760060e-04 | |
| de novo #495 | 2226 | | | | Sp2 | 2.199360e-01 | | Sp3 | 2.199360e-01 | | Klf3 | 2.199360e-01 | |
| de novo #496 | 1774 | | | | Mbd2 | 1.265820e-01 | | Smad1 | 1.265820e-01 | | Smad5 | 1.265820e-01 | |

| pattern | num_seqlets | modisco_cwm_fwd | modisco_cwm_rev | delta_effects | match0 | qval0 | match0_logo | match1 | qval1 | match1_logo | match2 | qval2 | match2_logo |
|---|---|---|---|---|---|---|---|---|---|---|---|---|---|
| de novo #497 | 838 | | | | Sp2 | 1.889550e-03 | | Sp3 | 2.815070e-03 | | Zfx | 2.815070e-03 | |
| de novo #498 | 1263 | | | | Zbtb14 | 3.540240e-01 | | Nrf1 | 3.540240e-01 | | Myc | 3.540240e-01 | |
| de novo #499 | 654 | | | | Sp2 | 1.023700e-04 | | Zfx | 1.861820e-04 | | Sp3 | 1.241750e-03 | |
| de novo #500 | 288 | | | | Taf1 | 3.593070e-04 | | Mbtps2 | 1.224330e-03 | | Yy1 | 1.224330e-03 | |
| de novo #501 | 1434 | | | | Sp2 | 4.411280e-04 | | Zfx | 1.968590e-03 | | Sp3 | 2.261570e-03 | |
| de novo #502 | 323 | | | | Sp2 | 7.996990e-07 | | Sp3 | 1.824870e-05 | | Zfx | 2.032780e-04 | |
| de novo #503 | 1154 | | | | Sp2 | 2.478820e-07 | | Sp3 | 1.610370e-05 | | Zfp566 | 8.610550e-04 | |
| de novo #504 | 243 | | | | Sp2 | 1.890070e-04 | | Sp3 | 6.242990e-04 | | Zfp467 | 1.196090e-03 | |
| de novo #505 | 19510 | | | | Sp2 | 2.215670e-03 | | Zfx | 3.720200e-03 | | Zfp64 | 3.720200e-03 | |
| de novo #506 | 2237 | | | | Zfx | 7.074940e-05 | | Sp2 | 5.815480e-04 | | Sp3 | 3.022840e-03 | |
| de novo #507 | 508 | | | | Taf1 | 2.155860e-02 | | Smad1 | 2.155860e-02 | | Smad5 | 2.155860e-02 | |
| de novo #508 | 112 | | | | Sp2 | 1.981210e-06 | | Zfx | 1.947790e-04 | | Sp3 | 1.947790e-04 | |
| de novo #509 | 1543 | | | | Sp2 | 3.472400e-05 | | Sp3 | 4.015850e-04 | | Zfx | 4.256450e-04 | |
| de novo #510 | 274 | | | | Sp2 | 6.912760e-07 | | Sp3 | 6.912760e-07 | | Zfp341 | 4.454340e-05 | |
| de novo #511 | 1889 | | | | Sp2 | 1.682040e-05 | | Zfx | 1.101120e-04 | | Sp3 | 1.664410e-04 | |

| pattern | num_seqlets | modisco_cwm_fwd | modisco_cwm_rev | delta_effects | match0 | qval0 | match0_logo | match1 | qval1 | match1_logo | match2 | qval2 | match2_logo |
|---|---|---|---|---|---|---|---|---|---|---|---|---|---|
| de novo #512 | 3988 | | | | Sp2 | 7.910310e-02 | | Dnmt1 | 7.910310e-02 | | Mxd1 | 7.910310e-02 | |
| de novo #513 | 33 | | | | Sp2 | 8.390720e-05 | | Sp3 | 3.997870e-03 | | Wt1 | 1.344560e-02 | |
| de novo #514 | 26 | | | | Sp2 | 5.556310e-05 | | Sp3 | 2.540460e-03 | | Elk4 | 3.379820e-03 | |
| de novo #515 | 17777 | | | | Zfx | 3.105580e-04 | | Sp2 | 2.345680e-03 | | Zfp64 | 7.655260e-03 | |
| de novo #516 | 266 | | | | Sp2 | 2.935220e-05 | | Sp3 | 5.782690e-05 | | Maz | 2.626850e-04 | |
| de novo #517 | 55 | | | | Sp2 | 2.800000e-03 | | Zfx | 2.800000e-03 | | Sp3 | 2.800000e-03 | |
| de novo #518 | 3694 | | | | Zfx | 1.633070e-04 | | Sp2 | 8.041980e-04 | | Mxd1 | 3.713290e-03 | |
| de novo #519 | 53 | | | | Sp2 | 7.344460e-07 | | Sp3 | 5.514750e-06 | | Zfp566 | 9.358760e-04 | |
| de novo #520 | 502 | | | | Sp2 | 3.895490e-07 | | Sp3 | 1.178000e-05 | | Zbtb17 | 4.681950e-04 | |
| de novo #521 | 430 | | | | Sp2 | 1.427090e-06 | | Sp3 | 1.438200e-06 | | Zfp341 | 3.641010e-05 | |
| de novo #522 | 7052 | | | | Sp2 | 1.787360e-10 | | Sp3 | 1.693940e-07 | | Zfx | 2.540330e-05 | |
| de novo #523 | 1505 | | | | Tfdp1 | 2.140370e-01 | | Tfdp2 | 2.140370e-01 | | Ctcfl | 3.052060e-01 | |
| de novo #524 | 137 | | | | Sp3 | 7.528860e-05 | | Sp2 | 9.980750e-05 | | Zfx | 1.830400e-04 | |
| de novo #525 | 128 | | | | Sp2 | 6.717960e-04 | | Sp3 | 3.291530e-03 | | Zfp64 | 6.098970e-03 | |
| de novo #526 | 230 | | | | Taf1 | 5.340220e-02 | | Zfx | 5.340220e-02 | | Sp3 | 5.340220e-02 | |

| pattern | num_seqlets | modisco_cwm_fwd | modisco_cwm_rev | delta_effects | match0 | qval0 | match0_logo | match1 | qval1 | match1_logo | match2 | qval2 | match2_logo |
|---|---|---|---|---|---|---|---|---|---|---|---|---|---|
| de novo #527 | 29 | | | | Sp2 | 1.134230e-04 | | Sp3 | 1.680310e-04 | | Klf6 | 2.881910e-03 | |
| de novo #528 | 303 | | | | Sp2 | 2.523590e-06 | | Sp3 | 3.201020e-06 | | Zfx | 1.651990e-04 | |
| de novo #529 | 685 | | | | Sp2 | 1.592250e-03 | | Sp3 | 2.901650e-03 | | Ctcfl | 5.621050e-03 | |
| de novo #530 | 1223 | | | | Zfx | 2.123270e-02 | | Sp2 | 2.708080e-02 | | Sp3 | 8.156360e-02 | |
| de novo #531 | 1579 | | | | Dlx5 | 1.864160e-02 | | Dlx2 | 1.864160e-02 | | Dlx4 | 1.864160e-02 | |
| de novo #532 | 2316 | | | | Dlx5 | 2.127220e-02 | | Dlx2 | 2.127220e-02 | | Dlx4 | 2.127220e-02 | |
| de novo #533 | 580 | | | | Nkx62 | 7.312600e-03 | | Dlx2 | 7.312600e-03 | | Hoxb7 | 0.007313 | |
| de novo #534 | 2401 | | | | Dlx2 | 1.187790e-02 | | Dlx5 | 1.187790e-02 | | Dlx4 | 0.012772 | |
| de novo #535 | 1496 | | | | Dlx5 | 2.614600e-02 | | Dlx2 | 2.614600e-02 | | Dlx4 | 2.614600e-02 | |
| de novo #536 | 1307 | | | | Dlx5 | 2.105110e-02 | | Dlx2 | 2.105110e-02 | | Dlx4 | 2.105110e-02 | |
| de novo #537 | 879 | | | | Zeb2 | 1.214930e-01 | | Figla | 1.214930e-01 | | Tcf12 | 1.214930e-01 | |
| de novo #538 | 65 | | | | Pparg | 9.389520e-03 | | Nr2c2 | 4.297410e-02 | | Nr1h3 | 1.713410e-01 | |
| de novo #539 | 16606 | | | | Erg | 1.825100e-04 | | Elf5 | 2.429310e-03 | | Ehf | 2.658150e-03 | |
| de novo #540 | 108 | | | | Etv2 | 7.552730e-03 | | Fli1 | 7.552730e-03 | | Etv3 | 7.552730e-03 | |
| de novo #541 | 9111 | | | | Elf2 | 1.807040e-05 | | Elf4 | 1.807040e-05 | | Erg | 1.807040e-05 | |

| pattern | num_seqlets | modisco_cwm_fwd | modisco_cwm_rev | delta_effects | match0 | qval0 | match0_logo | match1 | qval1 | match1_logo | match2 | qval2 | match2_logo |
|---|---|---|---|---|---|---|---|---|---|---|---|---|---|
| de novo #542 | 20716 | | | | Erg | 3.718530e-03 | | Elf5 | 1.149950e-02 | | Ehf | 0.023087 | |
| de novo #543 | 50 | | | | Elk1 | 3.637570e-04 | | Erg | 7.149790e-04 | | Erf | 8.550150e-04 | |
| de novo #544 | 13932 | | | | Erg | 3.819040e-06 | | Ehf | 1.678830e-04 | | Elf2 | 0.000168 | |
| de novo #545 | 22 | | | | Elf5 | 4.744610e-05 | | Ehf | 2.369810e-04 | | Elf3 | 5.471440e-04 | |
| de novo #546 | 40 | | | | Elk1 | 4.606100e-03 | | Erf | 5.121080e-03 | | ENSMUSG00000098546 | 5.121080e-03 | |
| de novo #547 | 12191 | | | | Erg | 7.482000e-06 | | Elf4 | 1.705560e-04 | | Ehf | 1.705560e-04 | |
| de novo #548 | 12601 | | | | Erg | 3.080990e-05 | | Elf4 | 1.011380e-04 | | Ehf | 1.430920e-04 | |
| de novo #549 | 14174 | | | | Erg | 7.557410e-06 | | Ehf | 2.110140e-04 | | Elf4 | 2.110140e-04 | |
| de novo #550 | 162 | | | | Spib | 5.994760e-03 | | Bcl11a | 6.983680e-03 | | Elk1 | 6.983680e-03 | |
| de novo #551 | 188 | | | | Elk1 | 9.170160e-03 | | Spib | 9.170160e-03 | | Erg | 9.170160e-03 | |
| de novo #552 | 18907 | | | | Erg | 4.224970e-04 | | Elf5 | 2.142280e-03 | | Ehf | 2.675360e-03 | |
| de novo #553 | 12059 | | | | Erg | 4.396570e-06 | | Elf2 | 2.105120e-04 | | Elf4 | 2.105120e-04 | |
| de novo #554 | 73 | | | | Ehf | 4.165400e-04 | | Elf5 | 4.348800e-04 | | Elf3 | 4.505640e-03 | |
| de novo #555 | 10146 | | | | Erg | 1.579100e-05 | | Elf2 | 1.579100e-05 | | Elf4 | 1.579100e-05 | |
| de novo #556 | 23 | | | | Etv5 | 1.569820e-02 | | Ehf | 1.569820e-02 | | Elk1 | 1.569820e-02 | |

| pattern | num_seqlets | modisco_cwm_fwd | modisco_cwm_rev | delta_effects | match0 | qval0 | match0_logo | match1 | qval1 | match1_logo | match2 | qval2 | match2_logo |
|---|---|---|---|---|---|---|---|---|---|---|---|---|---|
| de novo #557 | 21219 | | | | Erg | 4.005710e-03 | | Elf5 | 1.026660e-02 | | Ehf | 1.796820e-02 | |
| de novo #558 | 14248 | | | | Erg | 2.698990e-06 | | Elf4 | 2.607810e-04 | | Ehf | 2.607810e-04 | |
| de novo #559 | 1372 | | | | Klf11 | 2.545080e-03 | | Sp1 | 2.545080e-03 | | Sp4 | 0.002545 | |
| de novo #560 | 215 | | | | Zfp46 | 1.177460e-01 | | Klf1 | 2.472850e-01 | | Mga | 5.662640e-01 | |
| de novo #561 | 502 | | | | Ovol1 | 8.461450e-01 | | Zfp523 | 8.461450e-01 | | Nfe2l2 | 8.461450e-01 | |
| de novo #562 | 610 | | | | Ovol1 | 8.743530e-01 | | Zfp523 | 8.743530e-01 | | Nfe2l2 | 8.743530e-01 | |
| de novo #563 | 673 | | | | Ovol1 | 1.000000e+00 | | Nfe2l2 | 1.000000e+00 | | Fosl2 | 1.000000e+00 | |
| de novo #564 | 254 | | | | Zfp46 | 9.215100e-02 | | Klf1 | 2.408690e-01 | | Mga | 5.244020e-01 | |
| de novo #565 | 529 | | | | Ovol1 | 9.841460e-01 | | Nfe2l2 | 9.841460e-01 | | Fosl2 | 9.841460e-01 | |
| de novo #566 | 546 | | | | Ovol1 | 9.541000e-01 | | Nfe2l2 | 9.541000e-01 | | Fosl2 | 9.541000e-01 | |
| de novo #567 | 591 | | | | Ovol1 | 1.000000e+00 | | Nfe2l2 | 1.000000e+00 | | Fosl2 | 1.000000e+00 | |
| de novo #568 | 513 | | | | Ovol1 | 9.996910e-01 | | Nfe2l2 | 9.996910e-01 | | Fosl2 | 9.996910e-01 | |
| de novo #569 | 499 | | | | Ovol1 | 7.336430e-01 | | Zfp523 | 7.336430e-01 | | Fosl2 | 0.733643 | |
| de novo #570 | 578 | | | | Ovol1 | 1.000000e+00 | | Fosl2 | 1.000000e+00 | | Nfe2l2 | 1.000000e+00 | |
| de novo #571 | 742 | | | | Ovol1 | 1.000000e+00 | | Fosl2 | 1.000000e+00 | | Nfe2l2 | 1.000000 | |

| pattern | num_seqlets | modisco_cwm_fwd | modisco_cwm_rev | delta_effects | match0 | qval0 | match0_logo | match1 | qval1 | match1_logo | match2 | qval2 | match2_logo |
|---|---|---|---|---|---|---|---|---|---|---|---|---|---|
| de novo #572 | 556 | | | | Ovol1 | 9.473760e-01 | | Zfp523 | 9.473760e-01 | | Fosl2 | 9.473760e-01 | |
| de novo #573 | 27 | | | | Twist2 | 1.000000e+00 | | Zbtb18 | 1.000000e+00 | | Elk1 | 1.000000e+00 | |
| de novo #574 | 43 | | | | Dmrtc2 | 9.606400e-01 | | Hoxb9 | 9.606400e-01 | | Lcorl | 9.606400e-01 | |
| de novo #575 | 350 | | | | Zfp335 | 1.000000e+00 | | Rela | 1.000000e+00 | | Ovol1 | 1.000000e+00 | |
| de novo #576 | 641 | | | | Hoxc6 | 1.315880e-02 | | Hoxb1 | 1.962610e-02 | | Nkx62 | 1.962610e-02 | |
| de novo #577 | 1978 | | | | Zbtb7a | 1.206780e-03 | | Glis2 | 6.915580e-02 | | Hnf4a | 6.915580e-02 | |
| de novo #578 | 2007 | | | | Zbtb7a | 8.009020e-02 | | Glis2 | 8.009020e-02 | | Hnf4a | 0.080090 | |
| de novo #579 | 121 | | | | Glis3 | 2.471620e-02 | | Zfp641 | 2.471620e-02 | | Glis1 | 2.471620e-02 | |
| de novo #580 | 49 | | | | Plagl1 | 2.807300e-02 | | Zfp64 | 1.275470e-01 | | Zbtb7a | 1.883630e-01 | |
| de novo #581 | 3174 | | | | Glis2 | 2.232460e-02 | | Zbtb7a | 2.232460e-02 | | Zbtb7b | 8.065880e-02 | |
| de novo #582 | 3915 | | | | Hnf4a | 7.196640e-02 | | Zbtb7a | 7.196640e-02 | | Glis2 | 7.196640e-02 | |
| de novo #583 | 66 | | | | Maz | 1.438880e-04 | | Sp2 | 1.452870e-04 | | Sp3 | 0.000199 | |
| de novo #584 | 168 | | | | Zfp64 | 2.200920e-02 | | Zbtb7a | 2.918490e-02 | | Prdm4 | 1.231570e-01 | |
| de novo #585 | 1613 | | | | Hnf4a | 7.736430e-02 | | Glis2 | 7.736430e-02 | | Zbtb7a | 7.736430e-02 | |
| de novo #586 | 2077 | | | | Zbtb7a | 7.627620e-04 | | Hnf4a | 5.237320e-02 | | Glis1 | 0.068319 | |

| pattern | num_seqlets | modisco_cwm_fwd | modisco_cwm_rev | delta_effects | match0 | qval0 | match0_logo | match1 | qval1 | match1_logo | match2 | qval2 | match2_logo |
|---|---|---|---|---|---|---|---|---|---|---|---|---|---|
| de novo #587 | 4283 | | | | Zbtb7a | 2.929870e-02 | | Glis2 | 5.245800e-02 | | Glis1 | 5.245800e-02 | |
| de novo #588 | 2585 | | | | Zbtb7a | 2.692340e-03 | | Zbtb7b | 4.919140e-02 | | Glis1 | 5.804950e-02 | |
| de novo #589 | 2688 | | | | Zbtb7a | 8.071270e-04 | | Zbtb7b | 2.359110e-02 | | Glis2 | 6.916950e-02 | |
| de novo #590 | 102 | | | | Zfp641 | 4.606480e-02 | | Glis3 | 4.606480e-02 | | Glis1 | 4.606480e-02 | |
| de novo #591 | 5606 | | | | Zbtb7a | 3.106770e-02 | | Glis2 | 5.588290e-02 | | Glis1 | 5.588290e-02 | |
| de novo #592 | 4280 | | | | Glis2 | 1.631740e-02 | | Zbtb7a | 1.631740e-02 | | Zbtb7b | 9.336160e-02 | |
| de novo #593 | 3593 | | | | Zbtb7a | 7.458370e-04 | | Glis2 | 6.840810e-02 | | Zbtb7b | 6.840810e-02 | |
| de novo #594 | 1933 | | | | Glis2 | 2.620250e-02 | | Zbtb7a | 3.881970e-02 | | Glis1 | 1.541390e-01 | |
| de novo #595 | 3498 | | | | Zbtb7a | 4.505360e-02 | | Glis2 | 4.505360e-02 | | Glis1 | 4.505360e-02 | |
| de novo #596 | 2296 | | | | Maz | 6.902630e-08 | | Zfp467 | 6.902630e-08 | | Sp5 | 6.902630e-08 | |
| de novo #597 | 28 | | | | Zfp467 | 5.417810e-07 | | Maz | 5.417810e-07 | | Sp3 | 1.006690e-06 | |
| de novo #598 | 48 | | | | Maz | 3.952810e-03 | | Zfp467 | 3.952810e-03 | | Sp2 | 4.572040e-03 | |
| de novo #599 | 46 | | | | Sp5 | 4.976320e-07 | | Maz | 4.976320e-07 | | Sp2 | 4.976320e-07 | |
| de novo #600 | 350 | | | | Sp2 | 2.551980e-08 | | Zfp467 | 4.939670e-08 | | Maz | 4.939670e-08 | |
| de novo #601 | 1468 | | | | Sp5 | 7.040880e-06 | | Maz | 1.059470e-05 | | Zfp467 | 1.120480e-05 | |

| pattern | num_seqlets | modisco_cwm_fwd | modisco_cwm_rev | delta_effects | match0 | qval0 | match0_logo | match1 | qval1 | match1_logo | match2 | qval2 | match2_logo |
|---|---|---|---|---|---|---|---|---|---|---|---|---|---|
| de novo #602 | 1866 | | | | Sp5 | 3.394130e-08 | | Maz | 5.456360e-08 | | Zfp467 | 5.456360e-08 | |
| de novo #603 | 1757 | | | | Maz | 8.503750e-08 | | Zfp467 | 8.503750e-08 | | Sp5 | 1.052670e-07 | |
| de novo #604 | 843 | | | | Zfp341 | 2.290070e-06 | | Maz | 6.308190e-06 | | Zfp467 | 6.308190e-06 | |
| de novo #605 | 1347 | | | | Maz | 4.966840e-08 | | Zfp467 | 4.966840e-08 | | Sp5 | 2.744210e-07 | |
| de novo #606 | 115 | | | | Maz | 9.437750e-05 | | Sp2 | 2.892090e-04 | | Klf15 | 2.892090e-04 | |
| de novo #607 | 97 | | | | Klf15 | 3.291430e-04 | | Sp2 | 3.291430e-04 | | Maz | 3.291430e-04 | |
| de novo #608 | 80 | | | | Sp2 | 6.545750e-03 | | Maz | 6.545750e-03 | | Sp3 | 0.006546 | |
| de novo #609 | 250 | | | | Sp5 | 6.668320e-07 | | Maz | 6.668320e-07 | | Zfp467 | 6.668320e-07 | |
| de novo #610 | 1361 | | | | Sp3 | 8.639400e-07 | | Maz | 8.639400e-07 | | Zfp467 | 9.226630e-07 | |
| de novo #611 | 70 | | | | Sp5 | 8.193750e-05 | | Zfp281 | 8.193750e-05 | | Maz | 0.000082 | |
| de novo #612 | 33 | | | | Patz1 | 2.342000e-03 | | Wt1 | 3.477450e-03 | | Zbtb17 | 3.477450e-03 | |
| de novo #613 | 31 | | | | Sp2 | 1.176730e-04 | | Sp3 | 1.176730e-04 | | Wt1 | 2.576530e-03 | |
| de novo #614 | 213 | | | | Zfp64 | 5.893790e-03 | | Sp3 | 5.893790e-03 | | Sp2 | 0.005894 | |
| de novo #615 | 557 | | | | Nfkb1 | 1.019130e-05 | | Rela | 8.694490e-05 | | Relb | 2.314570e-04 | |
| de novo #616 | 342 | | | | Nfkb1 | 6.066220e-06 | | Rela | 4.763520e-05 | | Relb | 1.919460e-04 | |

| pattern | num_seqlets | modisco_cwm_fwd | modisco_cwm_rev | delta_effects | match0 | qval0 | match0_logo | match1 | qval1 | match1_logo | match2 | qval2 | match2_logo |
|---|---|---|---|---|---|---|---|---|---|---|---|---|---|
| de novo #617 | 232 | | | | Nfkb1 | 2.616990e-06 | | Rela | 2.982920e-05 | | Relb | 1.199840e-04 | |
| de novo #618 | 1213 | | | | Nfkb1 | 2.825430e-07 | | Rela | 2.496670e-05 | | Nfkb2 | 4.563870e-04 | |
| de novo #619 | 1683 | | | | Nfkb1 | 2.971170e-08 | | Rela | 1.518700e-05 | | Nfkb2 | 0.000159 | |
| de novo #620 | 517 | | | | Nfkb1 | 1.228650e-06 | | Rela | 1.427660e-04 | | Nfkb2 | 6.350990e-04 | |
| de novo #621 | 34 | | | | Nfkb1 | 1.815330e-05 | | Rela | 1.965980e-04 | | Nfkb2 | 1.228590e-03 | |
| de novo #622 | 669 | | | | Nfkb1 | 2.725840e-09 | | Rela | 1.716040e-05 | | Relb | 2.145970e-04 | |
| de novo #623 | 62 | | | | Nfkb1 | 1.513140e-06 | | Rela | 2.027780e-05 | | Relb | 2.215010e-04 | |
| de novo #624 | 241 | | | | Nfkb1 | 4.418140e-06 | | Rela | 3.402780e-05 | | Relb | 0.000325 | |
| de novo #625 | 1162 | | | | Elf5 | 2.568690e-01 | | Elf3 | 2.568690e-01 | | Spi1 | 0.256869 | |
| de novo #626 | 1019 | | | | Elf5 | 6.028360e-01 | | Elf3 | 6.028360e-01 | | Spi1 | 6.539720e-01 | |
| de novo #627 | 1431 | | | | Elf5 | 7.716080e-01 | | Elf3 | 7.716080e-01 | | Spi1 | 0.771608 | |
| de novo #628 | 2400 | | | | Elf5 | 1.566210e-01 | | Elf3 | 1.566210e-01 | | Spi1 | 1.939480e-01 | |
| de novo #629 | 1275 | | | | Elf5 | 2.220800e-01 | | Elf3 | 2.220800e-01 | | Spi1 | 2.527070e-01 | |
| de novo #630 | 23 | | | | Zfp212 | 8.411510e-01 | | Rreb1 | 9.075840e-01 | | Egr2 | 1.000000e+00 | |
| de novo #631 | 2943 | | | | Rreb1 | 1.000000e+00 | | T | 1.000000e+00 | | Egr2 | 1.000000e+00 | |

| pattern | num_seqlets | modisco_cwm_fwd | modisco_cwm_rev | delta_effects | match0 | qval0 | match0_logo | match1 | qval1 | match1_logo | match2 | qval2 | match2_logo |
|---|---|---|---|---|---|---|---|---|---|---|---|---|---|
| de novo #632 | 1357 | | | | Rreb1 | 1.000000e+00 | | T | 1.000000e+00 | | Egr2 | 1.000000e+00 | |
| de novo #633 | 184 | | | | Klf1 | 2.327640e-01 | | Egr2 | 2.327640e-01 | | Rreb1 | 2.327640e-01 | |
| de novo #634 | 814 | | | | Rreb1 | 1.000000e+00 | | T | 1.000000e+00 | | Egr2 | 1.000000e+00 | |
| de novo #635 | 3408 | | | | Rreb1 | 1.000000e+00 | | T | 1.000000e+00 | | Egr2 | 1.000000e+00 | |
| de novo #636 | 176 | | | | Egr2 | 1.229960e-02 | | Rreb1 | 1.229960e-02 | | Klf1 | 9.907460e-02 | |
| de novo #637 | 1236 | | | | Rreb1 | 1.000000e+00 | | T | 1.000000e+00 | | Egr2 | 1.000000e+00 | |
| de novo #638 | 2995 | | | | Rreb1 | 1.000000e+00 | | T | 1.000000e+00 | | Egr2 | 1.000000e+00 | |
| de novo #639 | 126 | | | | Egr2 | 1.779170e-01 | | Mxd1 | 1.779170e-01 | | Mxi1 | 1.779170e-01 | |
| de novo #640 | 65 | | | | Zfp661 | 2.019660e-03 | | Ctcfl | 2.134500e-02 | | Ctcf | 4.519710e-02 | |
| de novo #641 | 63 | | | | Zfp661 | 2.115260e-03 | | Ctcfl | 1.811420e-02 | | Ctcf | 3.817170e-02 | |
| de novo #642 | 51 | | | | Zfp661 | 1.630130e-03 | | Ctcfl | 1.187340e-02 | | Ctcf | 2.978670e-02 | |
| de novo #643 | 263 | | | | Ctcf | 1.000000e+00 | | Ctcfl | 1.000000e+00 | | Plag1 | 1.000000e+00 | |
| de novo #644 | 64 | | | | Zfp661 | 1.617330e-03 | | Ctcfl | 1.681910e-02 | | Ctcf | 3.777740e-02 | |
| de novo #645 | 357 | | | | Ctcf | 6.307190e-01 | | Ctcfl | 9.299740e-01 | | Klf13 | 1.000000e+00 | |
| de novo #646 | 183 | | | | Ctcf | 7.451990e-01 | | Ctcfl | 7.451990e-01 | | Plag1 | 1.000000e+00 | |

| pattern | num_seqlets | modisco_cwm_fwd | modisco_cwm_rev | delta_effects | match0 | qval0 | match0_logo | match1 | qval1 | match1_logo | match2 | qval2 | match2_logo |
|---|---|---|---|---|---|---|---|---|---|---|---|---|---|
| de novo #647 | 100 | | | | Zfp661 | 2.248880e-03 | | Ctcfl | 2.348340e-02 | | Ctcf | 4.838120e-02 | |
| de novo #648 | 58 | | | | Zfp661 | 1.693020e-03 | | Ctcfl | 7.864050e-03 | | Ctcf | 2.163480e-02 | |
| de novo #649 | 304 | | | | Sp2 | 2.730540e-03 | | Klf6 | 9.844340e-03 | | Sp3 | 1.028190e-02 | |
| de novo #650 | 39 | | | | Nfya | 1.711190e-01 | | Nfyc | 1.711190e-01 | | Nfyb | 2.019400e-01 | |
| de novo #651 | 55 | | | | Nfia | 7.061070e-01 | | Insm1 | 7.061070e-01 | | Nfix | 9.903440e-01 | |
| de novo #652 | 107 | | | | Rarg | 2.864390e-02 | | Hnf4g | 2.864390e-02 | | Nr2c2 | 2.864390e-02 | |
| de novo #653 | 44 | | | | Gata3 | 9.471880e-01 | | Hmg20a | 9.471880e-01 | | Hmg20b | 0.947188 | |
| de novo #654 | 100 | | | | Crebl2 | 1.630660e-01 | | Atf7 | 1.630660e-01 | | Creb5 | 1.630660e-01 | |
| de novo #655 | 139 | | | | Crebl2 | 2.215410e-01 | | Atf7 | 2.215410e-01 | | Creb5 | 2.215410e-01 | |
| de novo #656 | 61 | | | | Gmeb2 | 8.074260e-02 | | Gmeb1 | 8.074260e-02 | | Jdp2 | 2.559810e-01 | |
| de novo #657 | 2687 | | | | Creb3 | 3.008610e-04 | | Creb5 | 3.008610e-04 | | Gm28047 | 3.834930e-04 | |
| de novo #658 | 2465 | | | | Creb3 | 1.738010e-04 | | Creb5 | 1.738010e-04 | | Atf7 | 1.738010e-04 | |
| de novo #659 | 3329 | | | | Jdp2 | 1.183660e-04 | | Crebl2 | 3.466950e-04 | | Atf7 | 0.000347 | |
| de novo #660 | 126 | | | | Atf7 | 1.409430e-01 | | Crebl2 | 1.409430e-01 | | Creb5 | 1.759050e-01 | |
| de novo #661 | 3300 | | | | Atf7 | 1.740280e-04 | | Creb5 | 1.740280e-04 | | Creb3 | 1.740280e-04 | |

| pattern | num_seqlets | modisco_cwm_fwd | modisco_cwm_rev | delta_effects | match0 | qval0 | match0_logo | match1 | qval1 | match1_logo | match2 | qval2 | match2_logo |
|---|---|---|---|---|---|---|---|---|---|---|---|---|---|
| de novo #662 | 3645 | | | | Atf7 | 1.986230e-04 | | Creb5 | 1.986230e-04 | | Creb3 | 1.986230e-04 | |
| de novo #663 | 3645 | | | | Atf7 | 1.495860e-04 | | Creb3 | 1.495860e-04 | | Creb5 | 1.495860e-04 | |
| de novo #664 | 3224 | | | | Jdp2 | 1.253070e-04 | | Crem | 3.461900e-04 | | Atf7 | 0.000369 | |
| de novo #665 | 48 | | | | Atf7 | 4.217430e-03 | | Jdp2 | 4.217430e-03 | | Crem | 4.217430e-03 | |
| de novo #666 | 26 | | | | Atf7 | 6.279400e-04 | | Jdp2 | 6.279400e-04 | | Gm28047 | 2.274260e-03 | |
| de novo #667 | 114 | | | | Atf7 | 1.354580e-01 | | Crebl2 | 1.354580e-01 | | Creb5 | 0.152789 | |
| de novo #668 | 4347 | | | | Atf7 | 1.852520e-04 | | Creb3 | 1.852520e-04 | | Creb5 | 1.852520e-04 | |
| de novo #669 | 2414 | | | | Creb3 | 1.544650e-04 | | Creb5 | 1.544650e-04 | | Gm28047 | 1.989920e-04 | |
| de novo #670 | 2481 | | | | Jdp2 | 2.351460e-04 | | Creb5 | 2.351460e-04 | | Atf7 | 2.351460e-04 | |
| de novo #671 | 3103 | | | | Gm28047 | 1.276000e-04 | | Atf7 | 1.276000e-04 | | Creb3 | 1.276000e-04 | |
| de novo #672 | 2430 | | | | Creb5 | 2.752930e-04 | | Atf7 | 2.752930e-04 | | Creb3 | 2.752930e-04 | |
| de novo #673 | 128 | | | | Crebl2 | 1.560330e-01 | | Atf7 | 1.560330e-01 | | Creb5 | 1.699980e-01 | |
| de novo #674 | 53 | | | | Gm28047 | 8.561760e-04 | | Creb1 | 8.561760e-04 | | Creb3 | 8.561760e-04 | |
| de novo #675 | 44 | | | | Gm28047 | 6.356840e-04 | | Creb3 | 1.161210e-03 | | Atf2 | 6.908190e-03 | |
| de novo #676 | 1220 | | | | Gata6 | 2.187460e-02 | | Gata1 | 2.187460e-02 | | Tal1 | 0.036457 | |

| pattern | num_seqlets | modisco_cwm_fwd | modisco_cwm_rev | delta_effects | match0 | qval0 | match0_logo | match1 | qval1 | match1_logo | match2 | qval2 | match2_logo |
|---|---|---|---|---|---|---|---|---|---|---|---|---|---|
| de novo #677 | 21 | | | | Gata1 | 2.720950e-01 | | Gata2 | 2.720950e-01 | | Gata3 | 0.441625 | |
| de novo #678 | 135 | | | | Gata6 | 4.968210e-02 | | Gata1 | 4.968210e-02 | | Tal1 | 7.756850e-02 | |
| de novo #679 | 1443 | | | | Gata6 | 2.153830e-02 | | Gata1 | 2.153830e-02 | | Tal1 | 3.589650e-02 | |
| de novo #680 | 33 | | | | Nr4a2 | 2.985280e-03 | | Nr4a3 | 2.985280e-03 | | Rarg | 0.224949 | |
| de novo #681 | 29 | | | | Nr4a2 | 2.389060e-01 | | Nr4a3 | 2.389060e-01 | | Nr1d1 | 8.028310e-01 | |
| de novo #682 | 141 | | | | Hoxc10 | 4.271370e-02 | | Hoxd11 | 4.271370e-02 | | Hoxd10 | 0.042714 | |
| de novo #683 | 208 | | | | Sall4 | 7.024600e-02 | | Gli3 | 5.846140e-01 | | Gli1 | 0.584614 | |
| de novo #684 | 140 | | | | Sall4 | 1.127000e-01 | | Gli3 | 5.747630e-01 | | Gli1 | 5.747630e-01 | |
| de novo #685 | 91 | | | | Zfp287 | 3.388250e-01 | | Irf1 | 3.388250e-01 | | Stat1 | 3.388250e-01 | |
| de novo #686 | 403 | | | | Zfp287 | 8.450120e-10 | | Prdm6 | 1.102250e-07 | | Zfp422 | 9.605520e-05 | |
| de novo #687 | 378 | | | | Zfp287 | 3.920800e-04 | | Foxj1 | 4.457560e-02 | | Stat1 | 7.197330e-02 | |
| de novo #688 | 138 | | | | Zfp287 | 3.839660e-01 | | Irf4 | 3.839660e-01 | | Irf5 | 0.383966 | |
| de novo #689 | 115 | | | | Zfp287 | 3.861030e-02 | | Stat1 | 6.001900e-02 | | Prdm6 | 6.001900e-02 | |
| de novo #690 | 94 | | | | Zfp287 | 3.139150e-01 | | Stat1 | 3.139150e-01 | | Irf1 | 3.139150e-01 | |
| de novo #691 | 88 | | | | Zfp287 | 2.753550e-02 | | Zfp879 | 3.066570e-01 | | Hmg20a | 4.312980e-01 | |

| pattern | num_seqlets | modisco_cwm_fwd | modisco_cwm_rev | delta_effects | match0 | qval0 | match0_logo | match1 | qval1 | match1_logo | match2 | qval2 | match2_logo |
|---|---|---|---|---|---|---|---|---|---|---|---|---|---|
| de novo #692 | 646 | | | | Zfp287 | 1.125750e-07 | | Zfp422 | 2.257130e-06 | | Prdm6 | 2.638270e-06 | |
| de novo #693 | 547 | | | | Zfp422 | 2.188180e-05 | | Zfp287 | 1.587070e-04 | | Prdm6 | 1.289440e-03 | |
| de novo #694 | 101 | | | | Stat1 | 3.652300e-02 | | Zfp287 | 9.309640e-02 | | Irf1 | 9.309640e-02 | |
| de novo #695 | 3652 | | | | Zfp248 | 9.154430e-01 | | Zbtb32 | 9.154430e-01 | | Rhox11 | 1.000000e+00 | |
| de novo #696 | 4560 | | | | Rhox11 | 7.095720e-01 | | Zbtb32 | 7.095720e-01 | | Zfp248 | 7.095720e-01 | |
| de novo #697 | 212 | | | | Zfp146 | 1.000000e+00 | | Zbtb22 | 1.000000e+00 | | Zfp296 | 1.000000e+00 | |
| de novo #698 | 4805 | | | | Zbtb32 | 4.248070e-01 | | Zfp248 | 7.213790e-01 | | Zfp146 | 9.851350e-01 | |
| de novo #699 | 1567 | | | | Foxp2 | 1.000000e+00 | | Zfp146 | 1.000000e+00 | | Zbtb22 | 1.000000e+00 | |
| de novo #700 | 1313 | | | | Zbtb22 | 1.000000e+00 | | Zfp146 | 1.000000e+00 | | NaN | NaN | |
| de novo #701 | 1469 | | | | Zbtb22 | 1.000000e+00 | | Zfp879 | 1.000000e+00 | | Zscan4b | 1.000000e+00 | |
| de novo #702 | 2447 | | | | Zfp248 | 3.221570e-01 | | Rhox11 | 1.000000e+00 | | NaN | NaN | |
| de novo #703 | 3090 | | | | Zfp146 | 1.000000e+00 | | Zfp296 | 1.000000e+00 | | Zfp322a | 1.000000e+00 | |
| de novo #704 | 729 | | | | Mef2c | 6.549350e-01 | | Zfp422 | 6.549350e-01 | | Onecut3 | 6.549350e-01 | |
| de novo #705 | 481 | | | | Irf1 | 1.187770e-04 | | Stat2 | 3.373750e-03 | | Stat1 | 3.373750e-03 | |
| de novo #706 | 2512 | | | | Irf1 | 1.075050e-10 | | Stat2 | 2.192860e-05 | | Stat1 | 8.868950e-05 | |

| pattern | num_seqlets | modisco_cwm_fwd | modisco_cwm_rev | delta_effects | match0 | qval0 | match0_logo | match1 | qval1 | match1_logo | match2 | qval2 | match2_logo |
|---|---|---|---|---|---|---|---|---|---|---|---|---|---|
| de novo #707 | 1756 | | | | Irf1 | 1.631840e-09 | | Stat2 | 8.885310e-05 | | Prdm1 | 1.465280e-04 | |
| de novo #708 | 1401 | | | | Irf1 | 6.291360e-11 | | Prdm1 | 6.664970e-05 | | Irf8 | 2.958010e-04 | |
| de novo #709 | 2251 | | | | Irf1 | 3.349870e-11 | | Stat2 | 2.578790e-05 | | Stat1 | 5.687980e-05 | |
| de novo #710 | 424 | | | | Irf1 | 4.706910e-08 | | Prdm1 | 7.865050e-06 | | Stat2 | 2.997220e-05 | |
| de novo #711 | 1766 | | | | Irf1 | 7.599320e-11 | | Stat2 | 2.415010e-05 | | Stat1 | 8.169800e-05 | |
| de novo #712 | 412 | | | | Irf1 | 7.060990e-07 | | Prdm1 | 9.388650e-06 | | Stat2 | 2.203840e-04 | |
| de novo #713 | 3072 | | | | Irf1 | 9.290650e-11 | | Stat2 | 1.804560e-05 | | Stat1 | 0.000033 | |
| de novo #714 | 1928 | | | | Irf1 | 1.947380e-11 | | Prdm1 | 1.112800e-04 | | Irf8 | 2.801400e-04 | |
| de novo #715 | 67 | | | | Irf1 | 1.782060e-01 | | Irf4 | 1.782060e-01 | | Stat1 | 1.832100e-01 | |
| de novo #716 | 1580 | | | | Irf1 | 7.411170e-10 | | Prdm1 | 6.406950e-05 | | Irf8 | 2.547900e-04 | |
| de novo #717 | 502 | | | | Irf1 | 3.298400e-07 | | Prdm1 | 5.266450e-06 | | Stat2 | 0.000134 | |
| de novo #718 | 1335 | | | | Atf2 | 5.579790e-03 | | Gm28047 | 5.041290e-02 | | Creb5 | 0.050413 | |
| de novo #719 | 871 | | | | Atf2 | 9.764710e-04 | | Gm28047 | 2.563250e-02 | | Atf7 | 2.563250e-02 | |
| de novo #720 | 453 | | | | Atf2 | 6.145910e-03 | | Gm28047 | 5.211620e-02 | | Atf7 | 0.052116 | |
| de novo #721 | 1487 | | | | Zfp143 | 1.699630e-17 | | Thap11 | 3.119170e-17 | | Six5 | 0.000027 | |

| pattern | num_seqlets | modisco_cwm_fwd | modisco_cwm_rev | delta_effects | match0 | qval0 | match0_logo | match1 | qval1 | match1_logo | match2 | qval2 | match2_logo |
|---|---|---|---|---|---|---|---|---|---|---|---|---|---|
| de novo #722 | 1472 | | | | Zfp143 | 1.595050e-19 | | Thap11 | 3.325840e-19 | | Six5 | 3.872540e-05 | |
| de novo #723 | 22 | | | | Thap11 | 2.717160e-08 | | Zfp143 | 1.265700e-07 | | Six5 | 2.017030e-03 | |
| de novo #724 | 1414 | | | | Zfp143 | 1.164780e-17 | | Thap11 | 7.870210e-17 | | Six5 | 3.853210e-05 | |
| de novo #725 | 1439 | | | | Zfp143 | 1.246610e-16 | | Thap11 | 1.833010e-16 | | Six5 | 2.435710e-05 | |
| de novo #726 | 47 | | | | Thap11 | 1.958490e-10 | | Zfp143 | 3.923920e-10 | | Six5 | 2.736100e-03 | |
| de novo #727 | 1425 | | | | Zfp143 | 2.561840e-17 | | Thap11 | 1.376230e-16 | | Six5 | 4.421530e-05 | |
| de novo #728 | 1407 | | | | Zfp143 | 1.869390e-17 | | Thap11 | 1.070110e-16 | | Six5 | 3.799470e-05 | |
| de novo #729 | 1402 | | | | Zfp143 | 2.021880e-18 | | Thap11 | 1.902910e-17 | | Six5 | 1.311270e-05 | |
| de novo #730 | 1460 | | | | Zfp143 | 3.671010e-17 | | Thap11 | 6.185520e-17 | | Six5 | 4.120720e-05 | |
| de novo #731 | 1429 | | | | Thap11 | 1.011100e-17 | | Zfp143 | 1.994630e-17 | | Six5 | 5.003290e-05 | |
| de novo #732 | 1475 | | | | Thap11 | 8.565070e-20 | | Zfp143 | 1.831300e-19 | | Six5 | 3.711980e-05 | |
| de novo #733 | 47 | | | | Zfp143 | 2.714530e-03 | | Stat3 | 2.714530e-03 | | Stat4 | 2.714530e-03 | |
| de novo #734 | 1475 | | | | Zfp143 | 2.394360e-17 | | Thap11 | 2.394360e-17 | | Six5 | 3.536280e-05 | |
| de novo #735 | 1427 | | | | Zfp143 | 4.522780e-17 | | Thap11 | 4.522780e-17 | | Six5 | 0.000018 | |
| de novo #736 | 35 | | | | Nfix | 1.000000e+00 | | Nfia | 1.000000e+00 | | Rfx4 | 1.000000e+00 | |

| pattern | num_seqlets | modisco_cwm_fwd | modisco_cwm_rev | delta_effects | match0 | qval0 | match0_logo | match1 | qval1 | match1_logo | match2 | qval2 | match2_logo |
|---|---|---|---|---|---|---|---|---|---|---|---|---|---|
| de novo #737 | 101 | | | | Sp2 | 8.580790e-03 | | Mxd1 | 1.358930e-02 | | Mxi1 | 1.358930e-02 | |
| de novo #738 | 50 | | | | Nrf1 | 9.726960e-03 | | Zfp566 | 1.313810e-01 | | Elf1 | 0.131381 | |
| de novo #739 | 70 | | | | Sp2 | 2.083390e-06 | | Sp3 | 2.162430e-05 | | Wt1 | 1.138100e-03 | |
| de novo #740 | 622 | | | | Zbtb14 | 3.254080e-01 | | Nrf1 | 3.254080e-01 | | Zfp449 | 8.052930e-01 | |
| de novo #741 | 35 | | | | Sp2 | 1.499730e-03 | | Sp3 | 2.680200e-03 | | Zfx | 8.533670e-03 | |
| de novo #742 | 64 | | | | Sp2 | 1.122600e-03 | | Zfp64 | 3.250040e-03 | | Zfx | 3.250040e-03 | |
| de novo #743 | 19564 | | | | Sp2 | 1.078940e-02 | | Sp3 | 1.902520e-02 | | Zfx | 1.919390e-02 | |
| de novo #744 | 73 | | | | Nrf1 | 1.686340e-02 | | Sp2 | 1.007180e-01 | | Sp3 | 1.007180e-01 | |
| de novo #745 | 940 | | | | Zbtb14 | 3.075790e-01 | | Nrf1 | 3.075790e-01 | | Egr2 | 6.810890e-01 | |
| de novo #746 | 68 | | | | Zfp692 | 1.024550e-01 | | Hic2 | 1.119330e-01 | | Sp2 | 1.119330e-01 | |
| de novo #747 | 4844 | | | | Zfx | 5.233010e-04 | | Sp2 | 4.231530e-03 | | Mxd1 | 4.835150e-03 | |
| de novo #748 | 26 | | | | Sp2 | 7.886600e-05 | | Sp3 | 1.213960e-03 | | Mxd1 | 1.032660e-02 | |
| de novo #749 | 930 | | | | Nrf1 | 2.379610e-01 | | Zbtb14 | 2.379610e-01 | | Egr2 | 7.494450e-01 | |
| de novo #750 | 824 | | | | Nrf1 | 2.521350e-01 | | Zbtb14 | 2.521350e-01 | | Egr2 | 7.748480e-01 | |
| de novo #751 | 662 | | | | Nrf1 | 1.954190e-01 | | Zbtb14 | 1.954190e-01 | | Egr2 | 0.785539 | |

| pattern | num_seqlets | modisco_cwm_fwd | modisco_cwm_rev | delta_effects | match0 | qval0 | match0_logo | match1 | qval1 | match1_logo | match2 | qval2 | match2_logo |
|---|---|---|---|---|---|---|---|---|---|---|---|---|---|
| de novo #752 | 1932 | | | | Mlxipl | 4.420390e-04 | | Tfeb | 1.318980e-03 | | Usf1 | 1.545940e-03 | |
| de novo #753 | 22 | | | | Tfe3 | 4.317240e-04 | | Tfeb | 7.492260e-04 | | Tfec | 7.492260e-04 | |
| de novo #754 | 1746 | | | | Mlxipl | 4.850630e-04 | | Tfeb | 1.428790e-03 | | Tfe3 | 1.624080e-03 | |
| de novo #755 | 2245 | | | | Usf1 | 3.547330e-04 | | Tfeb | 7.772760e-04 | | Tfe3 | 1.098500e-03 | |
| de novo #756 | 2060 | | | | Tfeb | 2.633380e-05 | | Usf1 | 3.950070e-05 | | Arntl | 1.586110e-04 | |
| de novo #757 | 2589 | | | | Usf1 | 1.337120e-05 | | Tfeb | 2.627250e-05 | | Arntl | 3.011600e-04 | |
| de novo #758 | 1708 | | | | Usf1 | 1.272600e-04 | | Tfeb | 1.272600e-04 | | Arntl | 0.000194 | |
| de novo #759 | 1880 | | | | Tfeb | 5.485810e-05 | | Usf1 | 8.228710e-05 | | Arntl | 2.200540e-04 | |
| de novo #760 | 48 | | | | Tfeb | 1.783560e-03 | | Mxd1 | 1.783560e-03 | | Mxi1 | 1.783560e-03 | |
| de novo #761 | 1969 | | | | Tfeb | 1.100490e-03 | | Usf1 | 1.283900e-03 | | Mlxipl | 1.447250e-03 | |
| de novo #762 | 58 | | | | Tfeb | 7.865160e-04 | | Mitf | 4.018400e-03 | | Tfe3 | 4.018400e-03 | |
| de novo #763 | 2626 | | | | Tfeb | 1.241240e-03 | | Usf1 | 1.448110e-03 | | Mlxipl | 1.499320e-03 | |
| de novo #764 | 2229 | | | | Usf1 | 2.529390e-05 | | Tfeb | 4.544370e-05 | | Arntl | 1.402440e-04 | |
| de novo #765 | 2764 | | | | Usf1 | 7.164380e-04 | | Tfe3 | 7.164380e-04 | | Tfeb | 7.164380e-04 | |
| de novo #766 | 2879 | | | | Usf1 | 6.661910e-05 | | Tfeb | 9.534550e-05 | | Arntl | 0.000288 | |

| pattern | num_seqlets | modisco_cwm_fwd | modisco_cwm_rev | delta_effects | match0 | qval0 | match0_logo | match1 | qval1 | match1_logo | match2 | qval2 | match2_logo |
|---|---|---|---|---|---|---|---|---|---|---|---|---|---|
| de novo #767 | 49 | | | | Maf | 9.446240e-01 | | Hnf4g | 9.446240e-01 | | Hnf4a | 9.446240e-01 | |
| de novo #768 | 37 | | | | Tfap4 | 8.656480e-01 | | Figla | 8.656480e-01 | | Maf | 8.656480e-01 | |
| de novo #769 | 51 | | | | Tfap4 | 8.494110e-01 | | Maf | 8.494110e-01 | | Hnf4g | 8.494110e-01 | |
| de novo #770 | 30 | | | | Tfap4 | 9.407930e-01 | | Maf | 1.000000e+00 | | Figla | 1.000000 | |
| de novo #771 | 29 | | | | Tfap4 | 8.134290e-01 | | Maf | 1.000000e+00 | | Hnf4a | 1.000000e+00 | |
| de novo #772 | 39 | | | | Tfap4 | 8.337870e-01 | | Figla | 1.000000e+00 | | Maf | 1.000000 | |
| de novo #773 | 144 | | | | Zfp513 | 1.000000e+00 | | Lmx1a | 1.000000e+00 | | Dmrtc2 | 1.000000 | |
| de novo #774 | 217 | | | | Hoxc10 | 2.034690e-01 | | Hoxa6 | 2.034690e-01 | | Hoxb8 | 2.034690e-01 | |
| de novo #775 | 1256 | | | | Zfp692 | 1.402370e-01 | | Glis3 | 1.402370e-01 | | Glis1 | 1.402370e-01 | |
| de novo #776 | 29 | | | | ENSMUSG00000097126 | 3.948810e-01 | | Rhox8 | 3.948810e-01 | | Hsf1 | 4.193210e-01 | |
| de novo #777 | 2011 | | | | Hoxd11 | 6.137180e-01 | | ENSMUSG00000097126 | 6.137180e-01 | | Rhox8 | 6.137180e-01 | |
| de novo #778 | 25 | | | | ENSMUSG00000097126 | 6.929790e-01 | | Rhox8 | 6.929790e-01 | | Six5 | 1.000000e+00 | |
| de novo #779 | 95 | | | | Rfx1 | 2.730600e-08 | | Rfx3 | 4.100160e-07 | | Rfx5 | 4.100160e-07 | |
| de novo #780 | 593 | | | | Rfx1 | 3.755350e-08 | | Rfx2 | 4.273450e-07 | | Rfx3 | 4.273450e-07 | |
| de novo #781 | 496 | | | | Rfx1 | 6.805510e-09 | | Rfx3 | 2.102160e-08 | | Rfx5 | 2.102160e-08 | |

| pattern | num_seqlets | modisco_cwm_fwd | modisco_cwm_rev | delta_effects | match0 | qval0 | match0_logo | match1 | qval1 | match1_logo | match2 | qval2 | match2_logo |
|---|---|---|---|---|---|---|---|---|---|---|---|---|---|
| de novo #782 | 517 | | | | Rfx1 | 6.270490e-09 | | Rfx3 | 5.771450e-08 | | Rfx5 | 5.771450e-08 | |
| de novo #783 | 87 | | | | Rfx1 | 2.712800e-08 | | Rfx3 | 5.312380e-08 | | Rfx5 | 5.539100e-08 | |
| de novo #784 | 74 | | | | Rfx1 | 1.599270e-08 | | Rfx3 | 1.599270e-08 | | Rfx5 | 3.704350e-08 | |
| de novo #785 | 530 | | | | Rfx1 | 1.547070e-08 | | Rfx3 | 6.614360e-08 | | Rfx5 | 6.614360e-08 | |
| de novo #786 | 112 | | | | Gli1 | 1.000000e+00 | | Zfp558 | 1.000000e+00 | | Zfp212 | 1.000000e+00 | |
| de novo #787 | 400 | | | | Lcorl | 1.125590e-01 | | Lcor | 1.125590e-01 | | Mef2a | 4.576510e-01 | |
| de novo #788 | 319 | | | | Zfx | 3.500380e-02 | | Klf3 | 6.164530e-01 | | Sp2 | 6.164530e-01 | |
| de novo #789 | 208 | | | | Zfx | 4.681980e-01 | | Klf3 | 4.681980e-01 | | Tead4 | 4.681980e-01 | |
| de novo #790 | 487 | | | | Klf4 | 5.646680e-01 | | Klf7 | 5.646680e-01 | | Klf12 | 5.646680e-01 | |
| de novo #791 | 316 | | | | Klf3 | 2.266030e-01 | | Klf5 | 2.266030e-01 | | Zfx | 3.282680e-01 | |
| de novo #792 | 1299 | | | | Zfx | 2.051630e-04 | | Sp2 | 5.288080e-03 | | Klf3 | 5.288080e-03 | |
| de novo #793 | 278 | | | | Zfx | 1.000000e+00 | | Trp63 | 1.000000e+00 | | Klf5 | 1.000000e+00 | |
| de novo #794 | 1231 | | | | Ddit3 | 1.000000e+00 | | Jdp2 | 1.000000e+00 | | Atf4 | 1.000000e+00 | |
| de novo #795 | 93 | | | | Msantd3 | 1.000000e+00 | | Zfp110 | 1.000000e+00 | | Zfp691 | 1.000000e+00 | |
| de novo #796 | 37 | | | | Hoxc12 | 1.000000e+00 | | Hoxd12 | 1.000000e+00 | | Hoxc11 | 1.000000 | |

| pattern | num_seqlets | modisco_cwm_fwd | modisco_cwm_rev | delta_effects | match0 | qval0 | match0_logo | match1 | qval1 | match1_logo | match2 | qval2 | match2_logo |
|---|---|---|---|---|---|---|---|---|---|---|---|---|---|
| de novo #797 | 100 | | | | Zbtb43 | 1.000000e+00 | | Aire | 1.000000e+00 | | Ovol1 | 1.000000e+00 | |
| de novo #798 | 13335 | | | | Zfp213 | 5.572320e-01 | | Smad1 | 5.572320e-01 | | Smad5 | 5.572320e-01 | |
| de novo #799 | 33 | | | | Foxi2 | 5.109770e-03 | | Foxi3 | 5.109770e-03 | | Foxg1 | 8.589870e-03 | |
| de novo #800 | 28 | | | | Foxi2 | 2.626300e-03 | | Foxi3 | 2.626300e-03 | | Foxg1 | 0.009708 | |
| de novo #801 | 40 | | | | Foxi2 | 4.287320e-03 | | Foxi3 | 4.287320e-03 | | Foxg1 | 0.014429 | |
| de novo #802 | 35 | | | | Foxi2 | 3.550630e-02 | | Foxi3 | 3.550630e-02 | | Foxg1 | 5.049670e-02 | |
| de novo #803 | 37 | | | | Foxi2 | 2.673450e-03 | | Foxi3 | 2.673450e-03 | | Foxg1 | 4.339960e-03 | |
| de novo #804 | 1409 | | | | Mbtps2 | 2.020200e-06 | | Yy1 | 2.020200e-06 | | Yy2 | 2.020200e-06 | |
| de novo #805 | 983 | | | | Mbtps2 | 3.128790e-06 | | Yy1 | 3.128790e-06 | | Yy2 | 3.128790e-06 | |
| de novo #806 | 1307 | | | | Mbtps2 | 2.219290e-06 | | Yy1 | 2.219290e-06 | | Yy2 | 2.219290e-06 | |
| de novo #807 | 340 | | | | Taf1 | 2.600770e-04 | | Mbtps2 | 1.378030e-03 | | Yy1 | 1.378030e-03 | |
| de novo #808 | 1207 | | | | Mbtps2 | 2.934530e-06 | | Yy1 | 2.934530e-06 | | Yy2 | 2.934530e-06 | |
| de novo #809 | 1371 | | | | Mbtps2 | 2.753180e-06 | | Yy1 | 2.753180e-06 | | Yy2 | 2.753180e-06 | |
| de novo #810 | 1359 | | | | Mbtps2 | 2.061760e-06 | | Yy1 | 2.061760e-06 | | Yy2 | 0.000002 | |
| de novo #811 | 1560 | | | | Mbtps2 | 1.278130e-06 | | Yy1 | 1.278130e-06 | | Yy2 | 1.278130e-06 | |

| pattern | num_seqlets | modisco_cwm_fwd | modisco_cwm_rev | delta_effects | match0 | qval0 | match0_logo | match1 | qval1 | match1_logo | match2 | qval2 | match2_logo |
|---|---|---|---|---|---|---|---|---|---|---|---|---|---|
| de novo #812 | 1447 | | | | Mbtps2 | 1.598220e-06 | | Yy1 | 1.598220e-06 | | Yy2 | 1.598220e-06 | |
| de novo #813 | 348 | | | | Taf1 | 3.463260e-05 | | Mbtps2 | 3.463260e-05 | | Yy1 | 3.463260e-05 | |
| de novo #814 | 444 | | | | Taf1 | 1.481250e-04 | | Mbtps2 | 1.481250e-04 | | Yy1 | 1.481250e-04 | |
| de novo #815 | 1322 | | | | Mbtps2 | 2.790440e-06 | | Yy1 | 2.790440e-06 | | Yy2 | 2.790440e-06 | |
| de novo #816 | 1812 | | | | Mbtps2 | 1.044780e-06 | | Yy1 | 1.044780e-06 | | Yy2 | 1.044780e-06 | |
| de novo #817 | 1203 | | | | Mbtps2 | 4.837490e-06 | | Yy1 | 4.837490e-06 | | Yy2 | 4.837490e-06 | |
| de novo #818 | 1129 | | | | Mbtps2 | 3.719150e-06 | | Yy1 | 3.719150e-06 | | Yy2 | 0.000004 | |
| de novo #819 | 112 | | | | Mbtps2 | 3.609060e-05 | | Yy1 | 3.609060e-05 | | Yy2 | 3.609060e-05 | |
| de novo #820 | 1359 | | | | Zfp692 | 1.271090e-01 | | Glis3 | 1.271090e-01 | | Glis1 | 1.271090e-01 | |
| de novo #821 | 926 | | | | Dnmt1 | 3.771670e-01 | | Sp1 | 3.771670e-01 | | Klf11 | 3.771670e-01 | |
| de novo #822 | 829 | | | | Zbtb32 | 3.367220e-01 | | Lin28a | 3.367220e-01 | | Lin28b | 3.367220e-01 | |
| de novo #823 | 103 | | | | Klf15 | 5.209350e-02 | | Patz1 | 2.607040e-01 | | Wt1 | 0.308157 | |
| de novo #824 | 745 | | | | Ddit3 | 1.000000e+00 | | Jdp2 | 1.000000e+00 | | Atf4 | 1.000000e+00 | |
| de novo #825 | 623 | | | | Cebpb | 7.806750e-05 | | Cebpg | 1.851200e-04 | | Dbp | 6.527810e-04 | |
| de novo #826 | 839 | | | | Cebpg | 5.629310e-05 | | Cebpb | 1.151730e-04 | | Cebpa | 3.713930e-04 | |

| pattern | num_seqlets | modisco_cwm_fwd | modisco_cwm_rev | delta_effects | match0 | qval0 | match0_logo | match1 | qval1 | match1_logo | match2 | qval2 | match2_logo |
|---|---|---|---|---|---|---|---|---|---|---|---|---|---|
| de novo #827 | 963 | | | | Cebpg | 2.508040e-06 | | Cebpb | 5.029810e-06 | | Cebpa | 1.928110e-04 | |
| de novo #828 | 1615 | | | | Cebpb | 4.272830e-06 | | Cebpg | 3.076200e-05 | | Cebpa | 0.000372 | |
| de novo #829 | 344 | | | | Cebpb | 1.596370e-04 | | Dbp | 1.596370e-04 | | Hlf | 3.172690e-04 | |
| de novo #830 | 840 | | | | Cebpb | 8.140910e-07 | | Cebpg | 9.533170e-06 | | Cebpa | 9.032300e-05 | |
| de novo #831 | 188 | | | | Tef | 2.803680e-03 | | Dbp | 2.803680e-03 | | Hlf | 2.803680e-03 | |
| de novo #832 | 220 | | | | Zbtb7c | 2.415940e-02 | | Ctcfl | 9.940760e-02 | | Zfp661 | 9.940760e-02 | |
| de novo #833 | 827 | | | | Ddit3 | 1.000000e+00 | | Jdp2 | 1.000000e+00 | | Fosl1 | 1.000000e+00 | |
| de novo #834 | 92 | | | | Rfx7 | 6.920220e-02 | | Rfx6 | 6.920220e-02 | | Rfx4 | 7.845180e-02 | |
| de novo #835 | 189 | | | | Rfx7 | 8.145620e-02 | | Rfx4 | 8.145620e-02 | | Rfx8 | 8.145620e-02 | |
| de novo #836 | 72 | | | | Rfx6 | 1.250620e-01 | | Rfx4 | 1.250620e-01 | | Rfx7 | 1.250620e-01 | |
| de novo #837 | 52 | | | | Rfx6 | 2.616870e-03 | | Ovol1 | 3.241280e-01 | | Zfp740 | 5.138190e-01 | |
| de novo #838 | 318 | | | | Rfx7 | 8.290820e-02 | | Rfx6 | 8.290820e-02 | | Rfx4 | 1.302900e-01 | |
| de novo #839 | 391 | | | | Zbtb33 | 6.472020e-05 | | Srebf2 | 4.317450e-01 | | E2f8 | 1.000000e+00 | |
| de novo #840 | 397 | | | | Zbtb33 | 3.640190e-05 | | Srebf2 | 3.826190e-01 | | E2f8 | 1.000000 | |
| de novo #841 | 387 | | | | Zbtb33 | 2.480140e-05 | | Srebf2 | 3.355080e-01 | | E2f8 | 1.000000e+00 | |

| pattern | num_seqlets | modisco_cwm_fwd | modisco_cwm_rev | delta_effects | match0 | qval0 | match0_logo | match1 | qval1 | match1_logo | match2 | qval2 | match2_logo |
|---|---|---|---|---|---|---|---|---|---|---|---|---|---|
| de novo #842 | 375 | | | | Zbtb33 | 1.125320e-04 | | Srebf2 | 4.009510e-01 | | E2f8 | 1.000000e+00 | |
| de novo #843 | 373 | | | | Zbtb33 | 5.158180e-05 | | Srebf2 | 3.630690e-01 | | E2f8 | 1.000000e+00 | |
| de novo #844 | 353 | | | | Zbtb33 | 3.706960e-05 | | Srebf2 | 3.678380e-01 | | E2f8 | 1.000000e+00 | |
| de novo #845 | 350 | | | | Zbtb33 | 3.892130e-05 | | Srebf2 | 3.999330e-01 | | E2f8 | 1.000000e+00 | |
| de novo #846 | 364 | | | | Zbtb33 | 1.763280e-04 | | Srebf2 | 4.136620e-01 | | E2f8 | 1.000000e+00 | |
| de novo #847 | 415 | | | | Zbtb33 | 2.661440e-05 | | Srebf2 | 4.093920e-01 | | E2f8 | 1.000000e+00 | |
| de novo #848 | 397 | | | | Zbtb33 | 1.012100e-04 | | Srebf2 | 4.161420e-01 | | E2f8 | 1.000000 | |
| de novo #849 | 392 | | | | Zbtb33 | 7.258180e-05 | | Srebf2 | 3.786330e-01 | | E2f8 | 1.000000e+00 | |
| de novo #850 | 405 | | | | Zbtb33 | 1.666840e-04 | | Srebf2 | 4.274000e-01 | | E2f8 | 1.000000e+00 | |
| de novo #851 | 48 | | | | Elk1 | 1.477420e-01 | | Elf1 | 3.441610e-01 | | Ascl1 | 3.441610e-01 | |
| de novo #852 | 1112 | | | | Ddit3 | 1.000000e+00 | | Jdp2 | 1.000000e+00 | | Atf4 | 1.000000 | |
| de novo #853 | 498 | | | | Elk1 | 3.041220e-01 | | Erg | 3.041220e-01 | | Erf | 3.041220e-01 | |
| de novo #854 | 360 | | | | Elk1 | 2.776610e-01 | | Erg | 2.776610e-01 | | Erf | 0.277661 | |
| de novo #855 | 27 | | | | Sox3 | 5.060310e-01 | | Sox6 | 5.060310e-01 | | Tcf7l1 | 5.060310e-01 | |
| de novo #856 | 35 | | | | Sox3 | 5.494680e-01 | | Sox6 | 6.614350e-01 | | Cdx2 | 0.721729 | |

| pattern | num_seqlets | modisco_cwm_fwd | modisco_cwm_rev | delta_effects | match0 | qval0 | match0_logo | match1 | qval1 | match1_logo | match2 | qval2 | match2_logo |
|---|---|---|---|---|---|---|---|---|---|---|---|---|---|
| de novo #857 | 69 | | | | Zfp513 | 2.839400e-01 | | Foxm1 | 1.000000e+00 | | Atf3 | 1.000000 | |
| de novo #858 | 142 | | | | Zfp513 | 1.348040e-02 | | Atf3 | 4.195860e-01 | | Atf4 | 4.195860e-01 | |
| de novo #859 | 67 | | | | Zfp513 | 3.640060e-03 | | Atf3 | 1.000000e+00 | | Spic | 1.000000e+00 | |
| de novo #860 | 40 | | | | Zfp422 | 1.243490e-01 | | Zfp182 | 6.001590e-01 | | Bcl6b | 6.001590e-01 | |
| de novo #861 | 198 | | | | Nhlh1 | 3.853760e-01 | | Dnmt1 | 7.342690e-01 | | Sp3 | 7.342690e-01 | |
| de novo #862 | 96 | | | | Nfyc | 3.438180e-01 | | Nr4a2 | 3.438180e-01 | | Nr4a3 | 3.438180e-01 | |
| de novo #863 | 22 | | | | Sp2 | 4.915650e-01 | | Nfix | 4.915650e-01 | | Nfia | 4.915650e-01 | |
| de novo #864 | 438 | | | | Egr4 | 1.172050e-01 | | Zfp692 | 1.172050e-01 | | Egr3 | 0.117205 | |
| de novo #865 | 23 | | | | Klf2 | 6.158450e-01 | | Sp6 | 6.158450e-01 | | Hic2 | 0.615845 | |
| de novo #866 | 52 | | | | Gm28047 | 5.163340e-02 | | Atf2 | 5.163340e-02 | | Creb3 | 5.163340e-02 | |
| de novo #867 | 100 | | | | Zfp281 | 9.087940e-04 | | Zfp341 | 1.770330e-03 | | Wt1 | 1.876350e-03 | |
| de novo #868 | 52 | | | | Sp9 | 1.307970e-01 | | Hic2 | 1.307970e-01 | | Sp3 | 0.130797 | |
| de novo #869 | 320 | | | | Egr4 | 8.566680e-02 | | Zfp692 | 8.566680e-02 | | Egr2 | 8.566680e-02 | |
| de novo #870 | 61 | | | | Hoxb9 | 2.965560e-01 | | Cdx2 | 2.965560e-01 | | Hoxd9 | 2.965560e-01 | |
| de novo #871 | 39 | | | | Zfp1 | 7.326430e-01 | | Gm5294 | 7.326430e-01 | | Tbp | 0.732643 | |

| pattern | num_seqlets | modisco_cwm_fwd | modisco_cwm_rev | delta_effects | match0 | qval0 | match0_logo | match1 | qval1 | match1_logo | match2 | qval2 | match2_logo |
|---|---|---|---|---|---|---|---|---|---|---|---|---|---|
| de novo #872 | 257 | | | | Hoxb9 | 3.495340e-01 | | Cdx2 | 3.495340e-01 | | Tbp | 3.495340e-01 | |
| de novo #873 | 388 | | | | Cdx2 | 2.183960e-02 | | Hoxb9 | 4.119070e-02 | | Hoxc11 | 7.499810e-02 | |
| de novo #874 | 292 | | | | Cdx2 | 2.017840e-01 | | Hoxb9 | 2.017840e-01 | | Hoxd9 | 2.017840e-01 | |
| de novo #875 | 189 | | | | Sp2 | 3.820920e-03 | | Sp3 | 3.622310e-02 | | Klf5 | 3.622310e-02 | |
| de novo #876 | 29 | | | | Klf15 | 3.172150e-03 | | Maz | 7.417370e-03 | | Zfp566 | 7.417370e-03 | |
| de novo #877 | 240 | | | | Zfp64 | 2.970940e-04 | | Sp2 | 2.970940e-04 | | Sp3 | 0.000627 | |
| de novo #878 | 25 | | | | Sp2 | 1.334870e-02 | | Zfx | 1.334870e-02 | | Sp3 | 0.014580 | |
| de novo #879 | 195 | | | | Plagl1 | 1.381220e-01 | | Zfp692 | 5.487670e-01 | | Rxrb | 5.487670e-01 | |
| de novo #880 | 162 | | | | Zfp64 | 5.185630e-02 | | Zfp692 | 1.626580e-01 | | Zbtb7a | 1.801650e-01 | |
| de novo #881 | 35 | | | | Plagl1 | 2.327240e-01 | | Tfdp1 | 2.327240e-01 | | Tfdp2 | 2.327240e-01 | |
| de novo #882 | 163 | | | | Zfp513 | 4.260840e-03 | | Tfap2c | 1.000000e+00 | | Kmt2a | 1.000000e+00 | |
| de novo #883 | 238 | | | | Zfp513 | 2.433830e-03 | | Mbd2 | 1.000000e+00 | | Zbtb14 | 1.000000e+00 | |
| de novo #884 | 92 | | | | Nfia | 5.405600e-03 | | Nfix | 4.785680e-02 | | Rfx2 | 0.070284 | |
| de novo #885 | 126 | | | | Egr1 | 9.393120e-02 | | Egr3 | 9.393120e-02 | | Egr4 | 2.073930e-01 | |
| de novo #886 | 84 | | | | Zfp641 | 2.379920e-02 | | Glis1 | 2.379920e-02 | | Glis3 | 2.379920e-02 | |

| pattern | num_seqlets | modisco_cwm_fwd | modisco_cwm_rev | delta_effects | match0 | qval0 | match0_logo | match1 | qval1 | match1_logo | match2 | qval2 | match2_logo |
|---|---|---|---|---|---|---|---|---|---|---|---|---|---|
| de novo #887 | 69 | | | | Zfp641 | 5.822350e-02 | | Glis3 | 5.822350e-02 | | Glis1 | 0.058223 | |
| de novo #888 | 42 | | | | Sp1 | 1.245120e-02 | | Klf16 | 1.245120e-02 | | Sp2 | 1.245120e-02 | |
| de novo #889 | 31 | | | | Ctcf | 9.469410e-02 | | Mbtps2 | 9.469410e-02 | | Yy1 | 9.469410e-02 | |
| de novo #890 | 34 | | | | Ctcf | 4.547620e-02 | | Mbtps2 | 4.547620e-02 | | Yy1 | 4.547620e-02 | |
| de novo #891 | 60 | | | | Ctcf | 6.624900e-02 | | Zfp383 | 6.003150e-01 | | Taf1 | 6.003150e-01 | |
| de novo #892 | 69 | | | | Hic2 | 5.532840e-01 | | Klf16 | 1.000000e+00 | | Hic1 | 1.000000e+00 | |
| de novo #893 | 346 | | | | Zbtb7b | 1.089080e-02 | | Zbtb7a | 2.129220e-02 | | Zfp641 | 2.129220e-02 | |
| de novo #894 | 41 | | | | Foxm1 | 4.934880e-01 | | Atf4 | 1.000000e+00 | | Snai2 | 1.000000e+00 | |
| de novo #895 | 36 | | | | Ctcfl | 2.206550e-01 | | Patz1 | 2.206550e-01 | | Sp2 | 2.206550e-01 | |
| de novo #896 | 145 | | | | Ctcfl | 1.982930e-02 | | Tfap2c | 1.712290e-01 | | Tfap2a | 1.712290e-01 | |
| de novo #897 | 205 | | | | Sp2 | 8.940130e-03 | | Ctcfl | 8.940130e-03 | | Sp3 | 1.655340e-02 | |
| de novo #898 | 23 | | | | Irf9 | 1.000000e+00 | | Zfp213 | 1.000000e+00 | | Zkscan17 | 1.000000e+00 | |
| de novo #899 | 83 | | | | Ubp1 | 1.000000e+00 | | Hes6 | 1.000000e+00 | | Zbtb1 | 1.000000e+00 | |
| de novo #900 | 52 | | | | Sp2 | 1.000000e+00 | | Irf9 | 1.000000e+00 | | Hes6 | 1.000000e+00 | |
| de novo #901 | 681 | | | | Sox18 | 5.434120e-01 | | Sox7 | 5.434120e-01 | | Sox14 | 5.434120e-01 | |

| pattern | num_seqlets | modisco_cwm_fwd | modisco_cwm_rev | delta_effects | match0 | qval0 | match0_logo | match1 | qval1 | match1_logo | match2 | qval2 | match2_logo |
|---|---|---|---|---|---|---|---|---|---|---|---|---|---|
| de novo #902 | 24 | | | | Dmrt2 | 8.686820e-01 | | Hoxc4 | 8.686820e-01 | | Hoxa4 | 8.686820e-01 | |
| de novo #903 | 37 | | | | Foxg1 | 1.057650e-02 | | Foxi2 | 1.057650e-02 | | Foxi3 | 1.057650e-02 | |
| de novo #904 | 166 | | | | Zfp513 | 1.569830e-01 | | NaN | NaN | | NaN | NaN | |
| de novo #905 | 113 | | | | Zfp513 | 1.000000e+00 | | Zfp341 | 1.000000e+00 | | Klf15 | 1.000000 | |
| de novo #906 | 64 | | | | Elk1 | 5.766790e-02 | | Elk4 | 5.766790e-02 | | ENSMUSG00000098546 | 5.766790e-02 | |
| de novo #907 | 135 | | | | Zkscan17 | 1.000000e+00 | | Zbtb6 | 1.000000e+00 | | Mzf1 | 1.000000e+00 | |
| de novo #908 | 90 | | | | Lcorl | 1.000000e+00 | | Zfp740 | 1.000000e+00 | | Rreb1 | 1.000000 | |
| de novo #909 | 58 | | | | Foxm1 | 1.000000e+00 | | Sox1 | 1.000000e+00 | | Nhlh1 | 1.000000 | |
| de novo #910 | 219 | | | | Sp5 | 4.804700e-02 | | Sp3 | 4.804700e-02 | | Klf11 | 0.048047 | |
| de novo #911 | 203 | | | | Hmg20a | 2.487680e-01 | | Hmg20b | 2.487680e-01 | | Foxp1 | 0.248768 | |
| de novo #912 | 159 | | | | Zfp64 | 7.660710e-03 | | Zbtb6 | 8.819150e-02 | | Zbtb7a | 8.819150e-02 | |
| de novo #913 | 156 | | | | Zfp513 | 8.534520e-02 | | Ptf1a | 3.872740e-01 | | Dnmt1 | 3.872740e-01 | |
| de novo #914 | 90 | | | | Klf2 | 1.449740e-01 | | Klf8 | 1.449740e-01 | | Klf12 | 1.449740e-01 | |
| de novo #915 | 27 | | | | Esrrb | 2.360660e-01 | | Nr5a1 | 2.360660e-01 | | Esrra | 2.360660e-01 | |
| de novo #916 | 46 | | | | Nr5a1 | 4.169360e-01 | | Esrra | 4.169360e-01 | | Zfp524 | 7.531790e-01 | |

| pattern | num_seqlets | modisco_cwm_fwd | modisco_cwm_rev | delta_effects | match0 | qval0 | match0_logo | match1 | qval1 | match1_logo | match2 | qval2 | match2_logo |
|---|---|---|---|---|---|---|---|---|---|---|---|---|---|
| de novo #917 | 20 | | | | Nr5a1 | 3.176500e-01 | | Esrra | 3.176500e-01 | | Zfp524 | 1.000000e+00 | |
| de novo #918 | 61 | | | | Zfp423 | 1.000000e+00 | | Plagl1 | 1.000000e+00 | | Insm1 | 1.000000e+00 | |
| de novo #919 | 38 | | | | Irf9 | 1.000000e+00 | | Onecut1 | 1.000000e+00 | | Zfp524 | 1.000000e+00 | |
| de novo #920 | 98 | | | | Irf9 | 1.000000e+00 | | NaN | NaN | | NaN | NaN | |
| de novo #921 | 25 | | | | Irf5 | 3.350280e-01 | | Irf4 | 3.350280e-01 | | Irf9 | 0.335028 | |
| de novo #922 | 63 | | | | Sp2 | 1.000000e+00 | | Irf9 | 1.000000e+00 | | NaN | NaN | |
| de novo #923 | 24 | | | | Sp2 | 1.000000e+00 | | Irf9 | 1.000000e+00 | | Prdm14 | 1.000000 | |
| de novo #924 | 44 | | | | Irf9 | 1.000000e+00 | | NaN | NaN | | NaN | NaN | |
| de novo #925 | 50 | | | | Rorc | 1.643900e-01 | | Gmeb2 | 1.643900e-01 | | Nr1d2 | 1.643900e-01 | |
| de novo #926 | 53 | | | | Rorc | 1.312700e-01 | | Nr1d2 | 1.312700e-01 | | Gmeb2 | 1.727260e-01 | |
| de novo #927 | 55 | | | | Zfp513 | 5.154070e-01 | | Zbtb17 | 1.000000e+00 | | Rest | 1.000000e+00 | |
| de novo #928 | 34 | | | | Elk1 | 1.663850e-02 | | Elf1 | 1.663850e-02 | | Elf2 | 1.663850e-02 | |
| de novo #929 | 99 | | | | Zfp740 | 9.086260e-01 | | Zic5 | 9.086260e-01 | | Pbx1 | 0.908626 | |
| de novo #930 | 101 | | | | Hoxa7 | 1.000000e+00 | | Barhl2 | 1.000000e+00 | | Lcorl | 1.000000e+00 | |
| de novo #931 | 32 | | | | Zfp64 | 1.284770e-01 | | Usf1 | 3.030700e-01 | | Zfp770 | 3.842820e-01 | |

| pattern | num_seqlets | modisco_cwm_fwd | modisco_cwm_rev | delta_effects | match0 | qval0 | match0_logo | match1 | qval1 | match1_logo | match2 | qval2 | match2_logo |
|---|---|---|---|---|---|---|---|---|---|---|---|---|---|
| de novo #932 | 51 | | | | Tfap2a | 5.044140e-01 | | Tfap2d | 5.044140e-01 | | Nr1i3 | 1.000000e+00 | |
| de novo #933 | 28 | | | | Klf1 | 2.757340e-02 | | Trp63 | 2.757340e-02 | | Klf4 | 2.757340e-02 | |
| de novo #934 | 212 | | | | Zfp182 | 1.000000e+00 | | Foxj1 | 1.000000e+00 | | Foxq1 | 1.000000e+00 | |
| de novo #935 | 53 | | | | Zfp422 | 4.404100e-01 | | Hmga1 | 4.404100e-01 | | Hmga1rs1 | 0.440410 | |
| de novo #936 | 92 | | | | Smad1 | 2.883630e-01 | | Smad5 | 2.883630e-01 | | Taf1 | 5.112820e-01 | |
| de novo #937 | 85 | | | | Klf6 | 9.999260e-01 | | Klf5 | 9.999260e-01 | | Plag1 | 9.999260e-01 | |
| de novo #938 | 213 | | | | Sp4 | 4.071480e-02 | | Klf11 | 5.380940e-02 | | E2f4 | 6.428610e-02 | |
| de novo #939 | 24 | | | | Nhlh2 | 1.384520e-01 | | Runx3 | 7.255690e-01 | | Nhlh1 | 9.289600e-01 | |
| de novo #940 | 36 | | | | Zbtb7b | 1.378390e-01 | | Zfp423 | 1.378390e-01 | | Zbtb7a | 1.378390e-01 | |
| de novo #941 | 134 | | | | Atf4 | 3.530840e-01 | | Hoxa1 | 3.530840e-01 | | Msx3 | 3.530840e-01 | |
| de novo #942 | 137 | | | | Sp6 | 5.934270e-02 | | Klf1 | 2.984300e-01 | | Klf2 | 0.298430 | |
| de novo #943 | 48 | | | | Stat2 | 1.000000e+00 | | Stat1 | 1.000000e+00 | | Rorc | 1.000000e+00 | |
| de novo #944 | 39 | | | | Spic | 1.853470e-01 | | Etv6 | 3.813000e-01 | | Etv3 | 0.494329 | |
| de novo #945 | 37 | | | | Mxd1 | 2.085830e-04 | | Mxi1 | 2.085830e-04 | | Usf1 | 1.593050e-02 | |
| de novo #946 | 50 | | | | Arntl | 4.216840e-02 | | Tfe3 | 4.216840e-02 | | E2f6 | 4.216840e-02 | |

| pattern | num_seqlets | modisco_cwm_fwd | modisco_cwm_rev | delta_effects | match0 | qval0 | match0_logo | match1 | qval1 | match1_logo | match2 | qval2 | match2_logo |
|---|---|---|---|---|---|---|---|---|---|---|---|---|---|
| | | 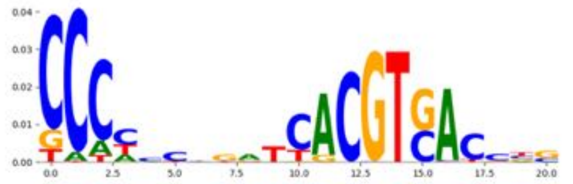 | 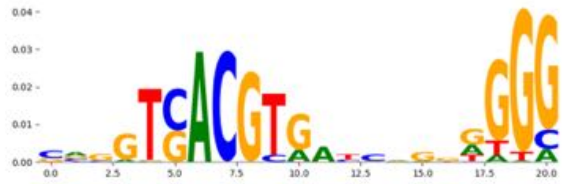 | 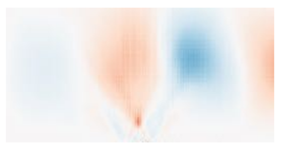 | | | 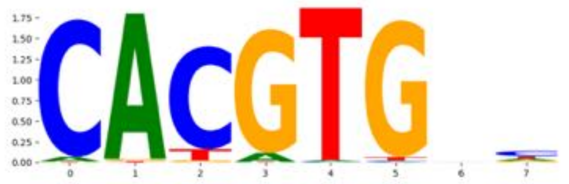 | | | 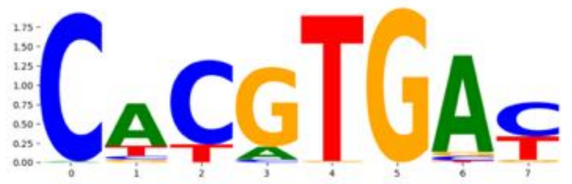 | | | 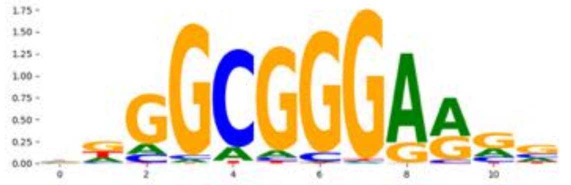 |
| de novo #947 | 85 | 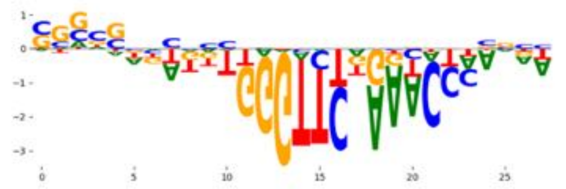 | 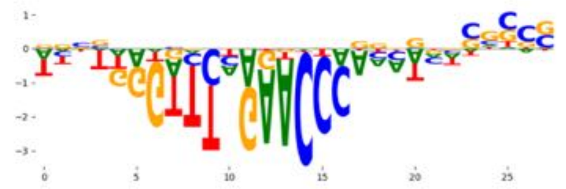 | 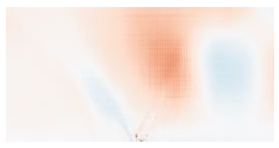 | Zfp69 | 1.000000e+00 | 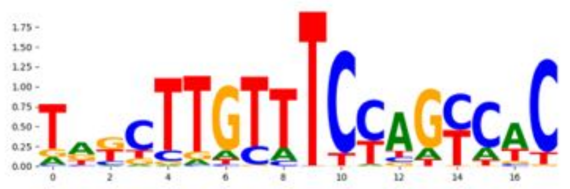 | Zfp1 | 1.000000e+00 | 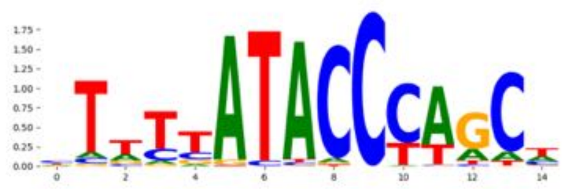 | Tcf12 | 1.000000e+00 | 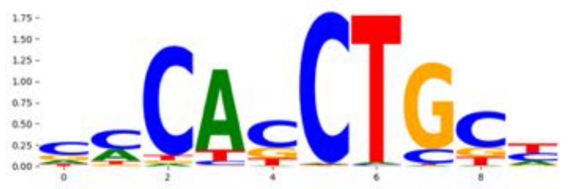 |
| de novo #948 | 64 | 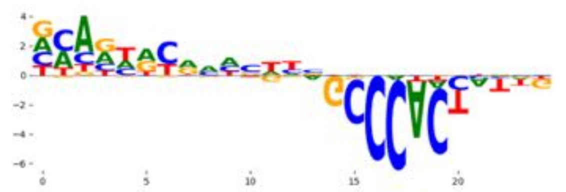 | 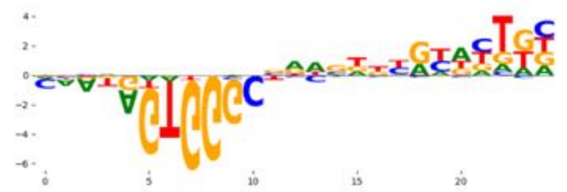 | 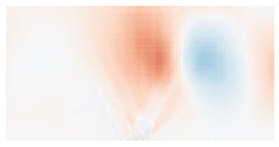 | Hic2 | 2.435940e-03 | 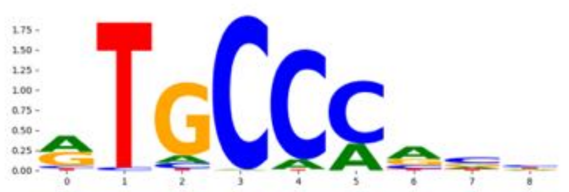 | Hic1 | 1.441250e-02 | 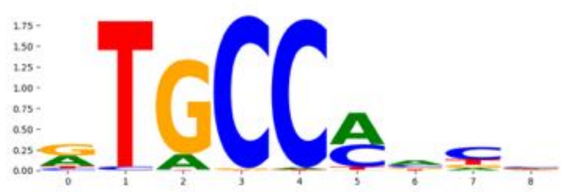 | Zkscan6 | 8.077890e-01 | 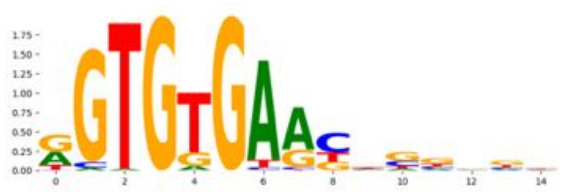 |
